# The Species-specific Acquisition and Diversification of a Novel Family of Killer Toxins in Budding Yeasts of the Saccharomycotina

**DOI:** 10.1101/2020.10.05.322909

**Authors:** Lance R. Fredericks, Mark D. Lee, Angela M. Crabtree, Josephine M. Boyer, Emily A. Kizer, Nathan T. Taggart, Samuel S. Hunter, Courtney B. Kennedy, Cody G. Willmore, Nova M. Tebbe, Jade S. Harris, Sarah N. Brocke, Paul A. Rowley

## Abstract

Killer toxins are extracellular antifungal proteins that are produced by a wide variety of fungi, including *Saccharomyces* yeasts. Although many *Saccharomyces* killer toxins have been previously identified, their evolutionary origins remain uncertain given that many of the se genes have been mobilized by double-stranded RNA (dsRNA) viruses. A survey of yeasts from the *Saccharomyces* genus has identified a novel killer toxin with a unique spectrum of activity produced by *Saccharomyces paradoxus*. The expression of this novel killer toxin is associated with the presence of a dsRNA totivirus and a satellite dsRNA. Genetic sequencing of the satellite dsRNA confirmed that it encodes a killer toxin with homology to the canonical ionophoric K1 toxin from *Saccharomyces cerevisiae* and has been named K1-like (K1L). Genomic homologs of K1L were identified in six non-*Saccharomyces* yeast species of the Saccharomycotina subphylum, predominantly in subtelomeric regions of the yeast genome. The sporadic distribution of these genes supports their acquisition by horizontal gene transfer followed by diversification, with evidence of gene amplification and positive natural selection. When ectopically expressed in *S. cerevisiae* from cloned cDNAs, both K1L and its homologs can inhibit the growth of competing yeast species, confirming the discovery of a new family of biologically active killer toxins. The phylogenetic relationship between K1L and its homologs suggests gene flow via dsRNAs and DNAs across taxonomic divisions to enable the acquisition of a diverse arsenal of killer toxins for use in niche competition.

## Introduction

Many different species of fungi have been observed to produce proteinaceous killer toxins that inhibit the growth of competing fungal species [1–7]. The killer phenotype was reported in the budding yeast *Saccharomyces cerevisiae* in 1962, when Bevan *et al*. observed that spent culture medium had antifungal properties [8]. The potential future application of killer toxins as novel fungicides has led to the discovery of many different killer yeasts with varying specificities and toxicities [9,10]. In the *Saccharomyces* yeasts, including several commonly used laboratory strains, it is estimated that 9-10% are able to produce killer toxins [11,12]. Despite the number of known killer yeasts that have been identified, a complete understanding of the diversity of killer toxins and their evolutionary history is lacking, even within the *S. cerevisiae*, which has been used as a model organism to study killer toxins for decades.

In general, killer toxin production by *S. cerevisiae* is most often enabled by infection with double-stranded RNA (dsRNA) totiviruses of the family *Totiviridae* [13–15]. Totiviruses that infect *Saccharomyces* yeasts are approximately 4.6 kbp in length and only encode two proteins, Gag and Gag-pol (by a programmed −1 frameshift). These proteins are essential for the assembly of virus particles and the replication of viral RNAs [16,17]. Totiviruses therefore enable killer toxin production by acting as helper viruses for the replication and encapsidation of ‘M’ satellite dsRNAs, which often encode killer toxins. These satellite dsRNAs are not limited to *Saccharomyces* yeasts, as they have been identified within other yeasts of the phylum Ascomycota (i.e. *Zygosaccharomyces bailii, Torulaspora delbrueckii*, and *Hanseniaspora uvarum* [18–20]) and the phylum Basidiomycota (*Ustilago maydis* [21,22]). In the Ascomycota, the organization of dsRNA satellites is similar, with all sequenced dsRNA satellites encoding a 5’ terminal sequence motif with the consensus of G(A)4-6, one or more central homopolymeric adenine (poly(A)) tracts, and a 3’ UTR containing packaging and replication *cis*-acting elements [16]. In all known satellite dsRNAs, killer toxin genes are positioned upstream of the central poly(A) tract and encode a single open reading frame. Identifying and characterizing dsRNAs is challenging, as the sequencing of dsRNAs currently requires specialized techniques for nucleic acid purification and conversion to cDNAs [23–25]. This has limited our understanding of the diversity of dsRNA-encoded killer toxins within fungi.

In *Saccharomyces* yeasts, there are eight known satellite dsRNA-encoded killer toxins that have been identified in *Saccharomyces* yeasts (K1, K2, K28, Klus, K21, K45, K62, and K74), with the majority found in the species *S. paradoxus* [26–30]. Owing to their early identification and distinct mechanisms of action, most functional studies of killer toxins have focused on the *S. cerevisiae* killer toxins K1 and K28. *Saccharomyces*-associated killer toxin genes appear to be evolutionarily diverse and unrelated by nucleotide and amino acid sequence. Despite the lack of homology, there are similarities in the posttranslational modifications that occur during killer toxin maturation prior to extracellular export of the active toxin. Killer toxins are expressed as pre-processed toxins (preprotoxins) with hydrophobic signal peptides that are required for extracellular secretion [31]. These signal peptides are cleaved by a signal peptidase complex in the endoplasmic reticulum. In the case of K1 and K28 toxins, the resulting protoxins are glycosylated and then crosslinked by disulfide bonds in the endoplasmic reticulum. Disulfide bonds in killer toxins are critical for both protein stability and toxicity [32–34]. The disulfide-linked protoxins are further cleaved by carboxypeptidases in the Golgi network to yield mature toxins that are secreted by exocytosis [35]. Mature K1 and K28 toxins can be described as α/β heterodimers that are linked by interchain disulfide bonds. Once outside of the producer cell, mature killer toxins can exert their antifungal activities upon competing fungi.

The K1 toxin was the first killer toxin to be discovered in *S. cerevisiae* and the mechanism of action has been studied extensively (reviewed in [36]). K1 is an ionophoric toxin that attacks the cell membrane of susceptible yeast cells and is mechanistically similar to the K2 toxin [37]. Interaction of K1 with a susceptible cell occurs in a two-step process that involves the initial energy-independent binding of the a- and β-domains of the toxin to the β-1,6-D-glucan polysaccharide of the yeast cell wall [31,34,38,39]. After binding surface glucans, K1 translocates to the cell membrane where it interacts with a secondary receptor, Kre1p [40]. Although there is still uncertainty on the exact mechanism of action of K1, it is likely that intoxication is caused by a-domain dependent formation of membrane channels and the subsequent selective leakage of monovalent cations from the cytoplasm [41]. K1 is lethal even at low concentrations, causing the apoptosis of sensitive cells [42]. Importantly, killer toxin immunity is provided by the immature protoxin by a mechanism that is not well understood. For K1, this activity has been mapped to the a-subunit and 31 amino acids of the adjacent g-subunit that is usually removed during toxin maturation [39,43].

In this study we describe the identification of a novel killer toxin encoded by a satellite dsRNA in *S. paradoxus*. This killer toxin has low primary sequence identity to the canonical K1 toxin produced by *S. cerevisiae* but has a similar secondary structure and domain organization. Due to its relatedness to K1, we have named this new killer toxin K1-like (K1L). This is the first example of a dsRNA-encoded killer toxin from *Saccharomyces* yeasts that has significant homology to a larger family of DNA-encoded “K1 killer toxin-like” (*KKT*) genes within the Saccharomycotina. Cloning and ectopic expression of *K1L* and *KKT* genes confirmed that they are functional extracellular antifungal toxins. These proteins represent a new family of killer toxins that are both diverse and show signs of rapid protein evolution. This work provides insights into the expansion and horizontal transfer of killer toxin genes in yeasts, whether they are encoded upon DNAs or mobilized and replicated as dsRNAs by viruses.

## Methods

### Killer phenotype assays

Killer toxin production by yeasts was measured using killer yeast agar plates (YPD agar plates with 0.003% w/v methylene blue at pH 4.6), as described previously [23]. Toxin production was identified by either a zone of growth inhibition or methylene blue-staining of the susceptible lawn yeasts. The pH optima of killer toxins produced by different yeasts was measured on killer yeast agar plates adjusted to pH values of 4.0, 5.0 and 5.5. The diameter of growth inhibition zones was measured with calipers

### Killer toxin enrichment

Strains of killer yeast were grown in 2 mL of YPD medium (pH 4.6) overnight at room temperature with vigorous shaking (250 rpm). The culture was centrifuged at 3,100 × g for 5 min followed by filtration of the culture medium through a 0.22 μm filter. Filtered growth medium was added 1:1 with 4°C supersaturated ammonium sulfate solution and mixed by inversion and incubated on ice for 3 h. The precipitated proteins were collected by centrifugation at 20,800 × g for 10 min at 4°C. The supernatant was then removed, and the precipitated proteins suspended in 10 μL of YPD pH 4.6. Killer toxins were incubated either at room temperature, or heat-inactivating at 98°C for 2 min before treating lawns of susceptible yeasts spread onto killer assay agar plates.

### Curing *Saccharomyces* yeasts of dsRNAs

Cycloheximide, anisomycin, and high temperatures were used to create strains of yeasts that lack satellite dsRNAs. Yeasts were first grown overnight in 2 mL of liquid YPD medium before 1 μL of cell suspension was transferred to YPD agar with either cycloheximide (0.4 - 5.0 μM) or anisomycin (0.8 μM). Yeast cultures were incubated for 2-5 days at 23°C to recover surviving cells. Curing satellite dsRNAs using temperature involved incubating yeast cultures on YPD agar for 2-5 days at 30°C, 37°C, or 40°C. Growing cells were streaked onto YPD agar plates from the heat-treated agar plates and were incubated for an additional 2-3 days at 23°C. The colonies resulting from chemical or temperature treatment were analyzed for the loss of killer toxin production by replica plating onto killer assay agar plates seeded with a killer toxin sensitive yeast strain.

### Short read sequencing of satellite dsRNAs

The protocol followed for the preparation of dsRNAs, cDNAs, Nextera Illumina libraries, and sequence analysis were the same as we previously reported, with several amendments detailed below [23]. The extracted dsRNAs were not incubated with oligo d(T)25 magnetic beads and 2× LTE buffer was replaced with 2× STE (500 mM NaCl; 20 mM Tris-HCl, pH 8.0; 30 mM EDTA, pH 8.0). Reads were cleaned with fastp and assembled with SPAdes 3.13.0 [44,45]. HTStream (https://github.com/ibest/HTStream) was used to clean the reads with stringent parameters. SPAdes assembler v3.11.1 was used to assemble reads using default parameters. The contigs produced for each dataset were used to build a bowtie2 index and were mapped to create BAM files that were subsequently visualized using Geneious version 8.1 (https://www.geneious.com). Sequence reads were deposited to the NCBI Sequence Read Archive with the accession number: TBA.

### Sanger sequencing of SpV-M1L

Reverse transcriptase PCR was used to generate overlapping DNAs that represented the genetic sequence of the satellite dsRNA SpV-M1L. The approximate molecular weight of these DNAs was determined by agarose gel electrophoresis and capillary electrophoresis ((Fragment Analyzer, Agilent Technologies Inc, La Jolla, CA, USA). DNAs were cloned using the pCR-Blunt II-TOPO vector and subjected to Sanger sequencing (See File S3 for the full list of primers used).

### Cloning of genome-encoded killer toxin genes

Genomic DNAs were extracted from *K. africana, N. dairenensis, N. castellii, T. phaffii, P. membranifaciens* using the method of Hoffman and Wilson (1987) and were used as templates for PCR (see table S1 for the full list of primers used) [46]. Killer toxin genes were cloned into pCR8 by TOPO-TA cloning (Thermo Fisher) and the DNA sequences were confirmed by Sanger sequencing. The K1L gene was commercially synthesized (GeneArt by Thermo Fisher) and used as a PCR template to amplify K1L. The PCR-derived K1L gene was cloned into pCR8 by TOPO-TA cloning and confirmed via Sanger sequencing. All killer toxin genes were sub-cloned using Gateway technology into the destination vector pAG426-Gal-ccdB for ectopic expression in either *S. cerevisiae* or *S. paradoxus* [47,48].

### Expression of K1L and related homologs from the Ascomycota

For ectopic expression of killer toxins, plasmids encoding toxin genes were used to transform either *S. cerevisiae* BY4741 or a customized non-flocculant tractable derivative of *S. paradoxus* A12 (named A12C) [49]. Transformants were selected on complete medium lacking uracil. To assay toxin expression, a single colony of each transformed strain was used to inoculate a series of consecutive overnight cultures in 2 mL of complete medium lacking uracil first with dextrose, then raffinose, and finally galactose at 30°C with shaking (250 rpm). The optical density of the final 2 mL culture was normalized to an OD600 of 1.0 and 1 mL was centrifuged at 3,000 × g for 5 min. The supernatant was removed, and the cell pellet was disrupted by gentle agitation. 2.5 μL of the resulting cell slurry was used to inoculate YPD and YPG plates (with 0.003% w/v methylene blue, pH 4.6) seeded with a killer toxin-susceptible yeast strain. Inoculated plates were incubated for 48-72 h at 25°C until killer toxin production was visible (24-72 h).

### Phylogenetic analyses

Killer toxin gene sequences were aligned using MUSCLE and manually trimmed to represent the most confident alignment of the a-domain. MEGA (version 7) was used for phylogenetic analysis using neighbor-joining and maximum likelihood methodologies. The optimal model for amino acid substitution was determined as the Whelan and Goldman model with a gamma distribution. 500 bootstrap replicates were used to construct a phylogenetic model with the highest log-likelihood.

## Results

### The Identification of New Strains of Killer Toxin-Producing Yeasts

A total of 110 strains of *Saccharomyces* yeasts were obtained from the USDA Agricultural Research Service (ARS) culture collection and screened to identify the production of novel killer toxins. The first screen used eight yeasts from four different species as indicators of toxin production on “killer assay media” (YPD, pH 4.6 with methylene blue) and found that 22% (24 strains) could inhibit the growth of at least one strain of yeast spread as a lawn (File S1). To identify the types of killer toxins based on their unique spectrum of activities, 13 of the killer yeasts were further screened against 45 indicator lawns of yeasts. Four strains of *S. cerevisiae* that have been previously described in the literature to produce killer toxins of unknown types were also included (NCYC1001, NCYC190, CYC1058, and CYC1113) [50–52]. To facilitate the classification of different toxin types, yeasts that produce K1 (BJH001), K28 (MS300c), and K74 (Y8.5) killer toxins, and a non-killer yeast (*S. cerevisiae* 1116) were included for comparison. The degree of growth inhibition by each killer yeast was scored qualitatively based on the appearance of zones of growth inhibition and methylene blue staining of the surrounding indicator strain on agar plates (Fig. 1 and File S2).

**Fig. 1.**
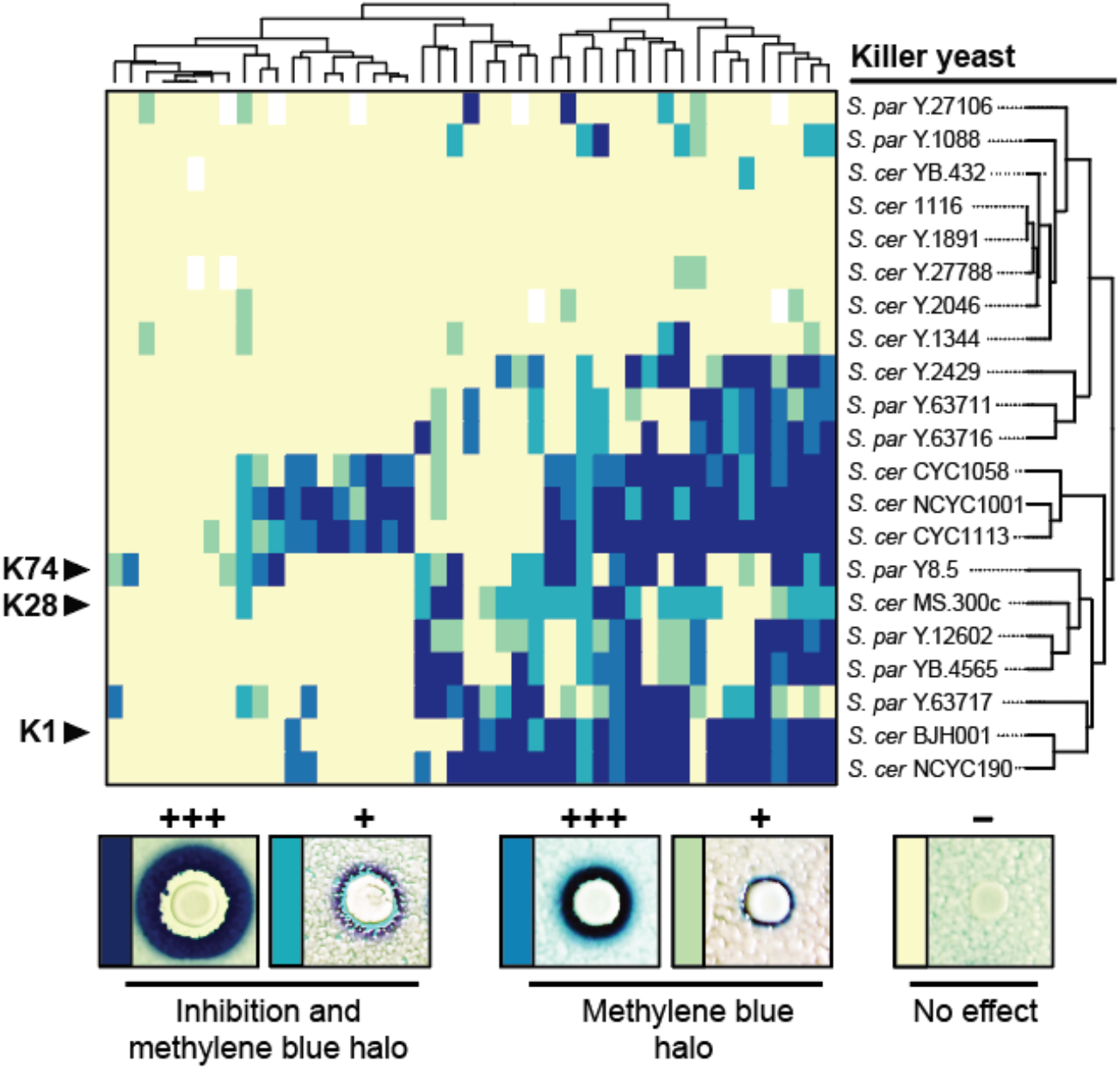
The strain and species specificity of killer toxins produced by *Saccharomyces* yeasts. A total of 21 killer yeasts were assayed for killer toxin production on agar plates seeded with 47 different indicator strains. Killer toxin activity was qualitatively assessed based on the presence and size of zones of growth inhibition or methylene blue staining around killer yeasts. Darker colors on the cluster diagram represent a more prominent killer phenotype with yellow indicating no detectable killer phenotype. The non-killer yeast strain *S. cerevisiae* 1116 was used as a negative control. Results were analyzed using the R package gplots to cluster killer yeast and indicator yeasts based on killer toxin cell tropism and susceptibility, respectively.

The known strain- and species-specificity of killer toxins have been previously used to biotype different yeasts [53–57] and to identify groups of unknown killer toxins [3,58]. Cluster analysis grouped the killer yeasts based on their ability to inhibit growth and revealed that no two strains of killer yeast have the exact same spectrum of antifungal activity. Clustering revealed that that the antifungal specificity of *S. cerevisiae* NCYC190 is closely correlated with that of the K1 killer yeast *S. cerevisiae* BJH001, with 91% identical interactions with competing lawn strains. The killer toxins K28 (MS300c) and K74 (Y8.5) have similar activities, even though they are different types of killer toxins [27,59]. *S. cerevisiae* CYC1113. CYC1058, and NCYC1001 have the broadest spectrum of activity but do not cluster with either K1, K28, or K74 killer yeasts. The killer toxin activity of *S. paradoxus* Y-63717 clusters with K1 (49% identical interactions), showing both gain and loss of function. This result is particularly intriguing because evidence of K1 production by *S. paradoxus* is inconsistent between different research groups with recent studies suggesting that K1 toxins are unique to *S. cerevisiae* [12,60,61]. The remaining killer yeasts that were identified have weaker inhibitory activities with weak clustering with known killer yeasts.

### Killer Yeasts Harbor Satellite dsRNAs that Encode Proteinaceous Killer Toxins

The production of killer toxins by yeasts is often associated with the presence of satellite dsRNAs that are maintained by totiviruses [17,36]. To determine if the killer phenotype correlates with the presence of dsRNAs, cellulose chromatography was used to selectively purify dsRNAs from 19 killer yeasts (including *S. cerevisiae* BJH001 as a positive control [23]). The analysis of extracted nucleic acids revealed that 68% of the killer yeasts contained dsRNAs indicative of totiviruses (~4.6 kbp) and satellite dsRNAs (<2 kbp) (Fig. 2A; top). The remaining killer yeasts were found to either contain only totiviruses (16%) or no dsRNAs (16%), which suggests that killer toxin production by these strains is genome encoded (Fig. S1). To confirm that the observed satellite dsRNAs encode killer toxin genes, killer yeasts were treated with either cycloheximide, anisomycin, or incubated at elevated temperatures to select for the loss of dsRNAs [62,63]. The majority of killer yeast (86%) lost their killer phenotype after exposure to chemical or thermal insult (Fig. 2A; bottom). Analysis of the dsRNAs within yeast strains that had lost killer toxin production showed the loss of satellite dsRNAs, but with the maintenance of totivirus dsRNAs (Fig. 2A; bottom). The same treatments were unable to select for the loss of the killer phenotype in five representative strains that lacked dsRNA satellites (Y-1344, Y-2046, YB-432, Y-1891, and Y-27788). Relative to the wild type strains, most of the cured strains exhibited an elevated copy number of totivirus dsRNAs. This phenomenon has been previously observed and is attributed to the fitness cost of parasitism by satellite dsRNAs [64].

**Fig. 2.**
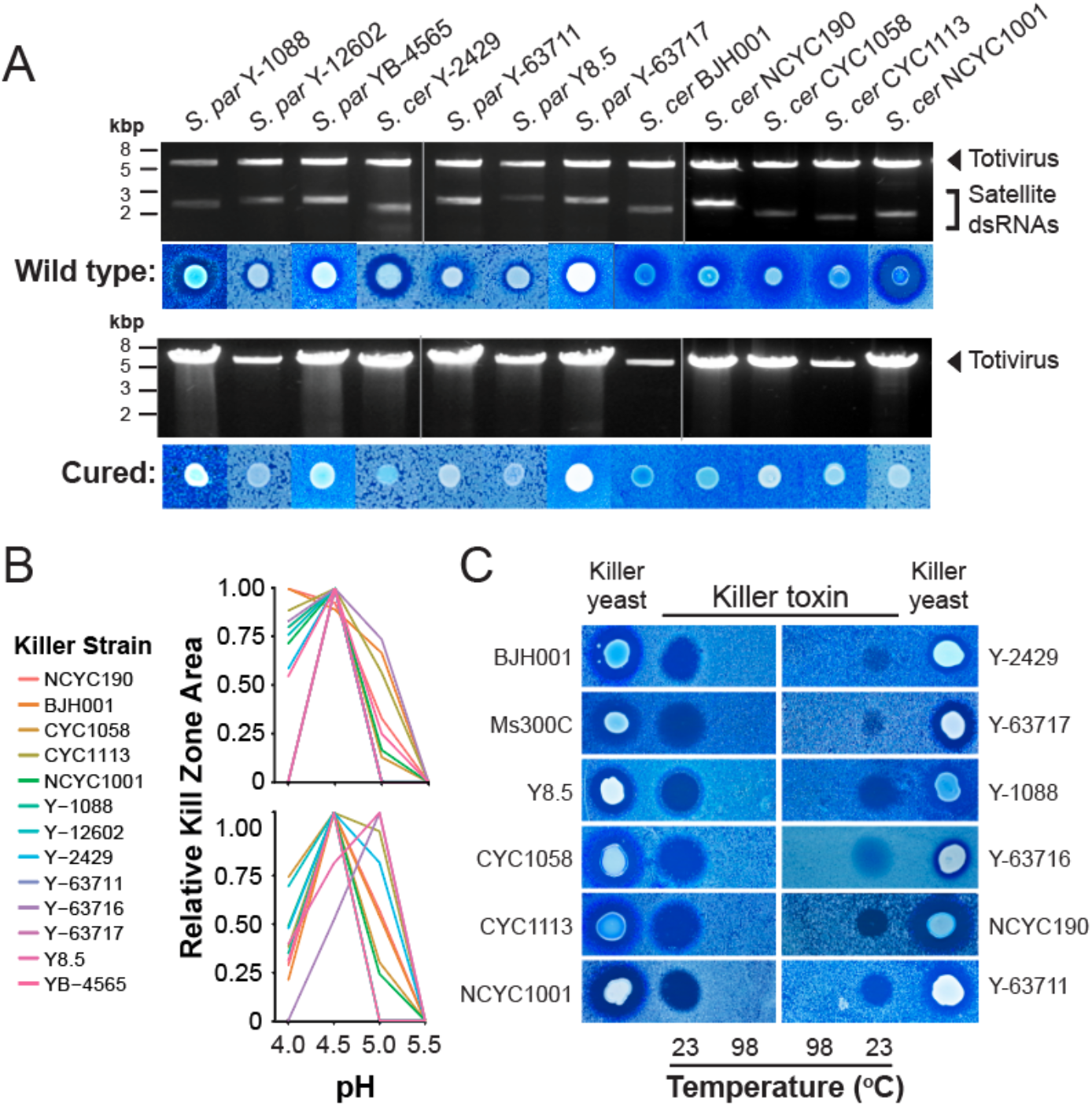
Analysis of dsRNAs present within killer yeasts and the biological properties of their killer toxins. (A) The extraction and analysis of dsRNAs from totiviruses and associated satellites in *Saccharomyces* yeasts that produce killer toxins (top panel) or have lost the killer phenotype by exposure to cycloheximide, anisomycin, or elevated temperatures (bottom panel). (B) The relative pH optimum of killer toxins against the indicator yeasts *S. cerevisiae* YB-4237 (top) and DSM70459 (bottom). (C) Enrichment and concentration of killer toxins from spent growth media by ammonium sulfate precipitation and their loss of inhibitory activity after incubation at 98°C.

The dsRNA-encoded killer toxins identified have a pH optimum between 4.5 and 5, with no inhibitory activity at pH 5.5 (Fig. 2B). To confirm that the identified killer toxins are proteinaceous, each was purified by ammonium sulfate precipitation and used to challenge susceptible yeasts. Zones of inhibition were clearly visible on confluent lawns of yeast cells for all of the killer toxins tested (Fig. 2C). The inhibitory activities of these precipitates were heat-labile, and the toxicity was lost after incubation at 98°C for 2 minutes. Together, these data suggest that these killer toxins have similar biochemical characteristics to known proteinaceous killer toxins despite their differing inhibitory effects towards yeasts.

### The Discovery of a New Killer Toxin Produced by S. paradoxus

To identify the unknown killer toxins produced by killer yeasts, dsRNAs were purified and subjected to a short-read sequencing pipeline for dsRNAs [23]. BLASTn analysis of *de novo* assembled contigs revealed that dsRNAs within strains CYC1058 and NCYC1001 encode canonical K2 toxins and NCYC190 a canonical K1 toxin (Fig. S2). The contigs derived from the dsRNAs of Y-63717 assembled into 125 different contigs, with six >750 bp in length and a coverage score >1,000 (Fig. 3A). BLASTn analysis of these high-quality contigs identified the dsRNA genome of the totivirus L-A-45 from *S. paradoxus* N-45 with 100% coverage and 95.5% nucleotide identity [27]. However, the remaining contigs did not match the nucleotide sequence of any known killer toxin in *Saccharomyces* yeasts. A combination of 5’ and 3’ RACE, reverse transcriptase PCR, and capillary electrophoresis was used to assemble the complete sequence of the putative dsRNA satellite from Y-63717 (Fig. 3B and S3). The novel satellite dsRNA is approximately 2371 bp in length with a single open reading frame (ORF) that encodes a protein of 340 amino acids. The 5’ ORF is positioned upstream of a central poly(A) tract of ~220 bp (Fig. 3B and S3). The 5’ terminus has a nucleotide sequence of 5’-GAAAAA that is found in many satellites dsRNAs (Fig. S3) and is predicted to fold into a large stem-loop structure (Fig. S4). Downstream of the poly(A) tract in the 3’ untranslated region (UTR) there are elements of secondary structure that are indicative of replication (terminal recognition element; *TRE*) and packaging signals (viral binding site; *VBS*) that have been well characterized in the canonical M1 satellite dsRNA from *S. cerevisiae* (ScV-M1) (Fig. 3B and S3) [65–67]. In addition to RNA secondary structures, there are also two direct repeats of the sequence motif named “Downstream of Poly(A)” (*DPA*; 5’-CTCACCYTGAGNHTAACTGG-3’) that is found in different satellite dsRNAs isolated from *S. paradoxus* (M45, M74, and M62), *S. cerevisiae* (M1 and Mlus), *Zygosaccharomyces bailii* (MZb), and *Torulaspora delbrueckii* (Mbarr) (Fig. S3) [19,68].

**Fig. 3.**
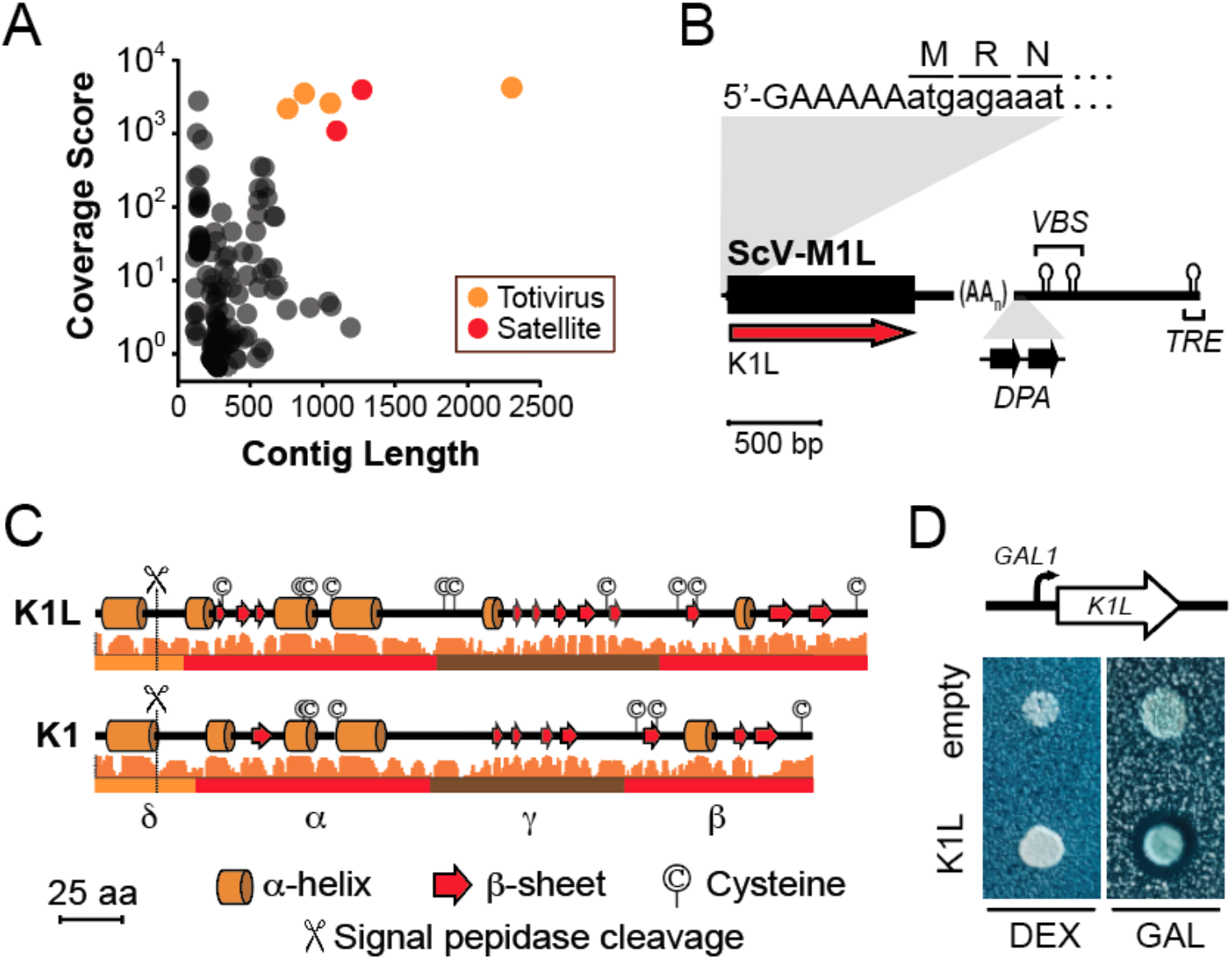
Short-read sequencing and analysis of the K1L killer toxin from *S. paradoxus* Y-63717. (A) Sequence contigs after *de novo* assembly of sequence reads represented by contig coverage score and contig length. BLASTx analysis was used to annotate contigs as similar to totiviruses or satellite dsRNAs. (B) Schematic of the organization of the SpV-M1L satellite dsRNA (C) Jpred secondary structure prediction of the K1 killer toxin and K1L killer toxin from Sp-M1L with confidence score plotted as a histogram. Predicted domain boundaries are drawn below the secondary structure prediction. (D) Ectopic expression of K1L from a plasmid in the non-killer yeast *S. paradoxus* A12 induced by the presence of galactose in the growth media.

The length and positioning of the 5’ ORF of the satellite dsRNA in Y-63717 strongly suggests that it encodes a killer toxin (Fig. 2). A PSI-BLAST search of the NCBI database with two iterations found that the putative killer toxin has weak homology to the canonical K1 toxin from *S. cerevisiae* (99% coverage, 21% amino acid identity, e-value 4 x 10^-17^) (Fig. 4A). Based on this homology, the putative killer toxin was named K1L (K1-like) and the dsRNA satellite was named Saccharomyces paradoxus virus M1-like (SpV-M1L). The organization of the functional domains of K1L appears to be similar to K1 based on the secondary structure, conserved cysteine residues, and the predicted signal peptidase cleavage sites (Fig. 3C) [69]. K1L contains ten cysteine residues, two of which are likely important for interchain disulfide linkage (Cys91) and killer toxin immunity (Cys257) based on their alignment with cysteines from K1 [32]. To confirm that the K1L is an active killer toxin, it was ectopically expressed by the non-killer strain *S. paradoxus* A12C using a galactose inducible promoter. A well-defined zone of growth inhibition was visible when the strain was grown on galactose-containing media (Fig. 3D). No K1L toxin expression was observed when cells were plated on dextrose-containing growth media. Together, these data confirm the identification of a new dsRNA satellite in *Saccharomyces* yeasts and a novel killer toxin related to K1.

**Fig. 4.**
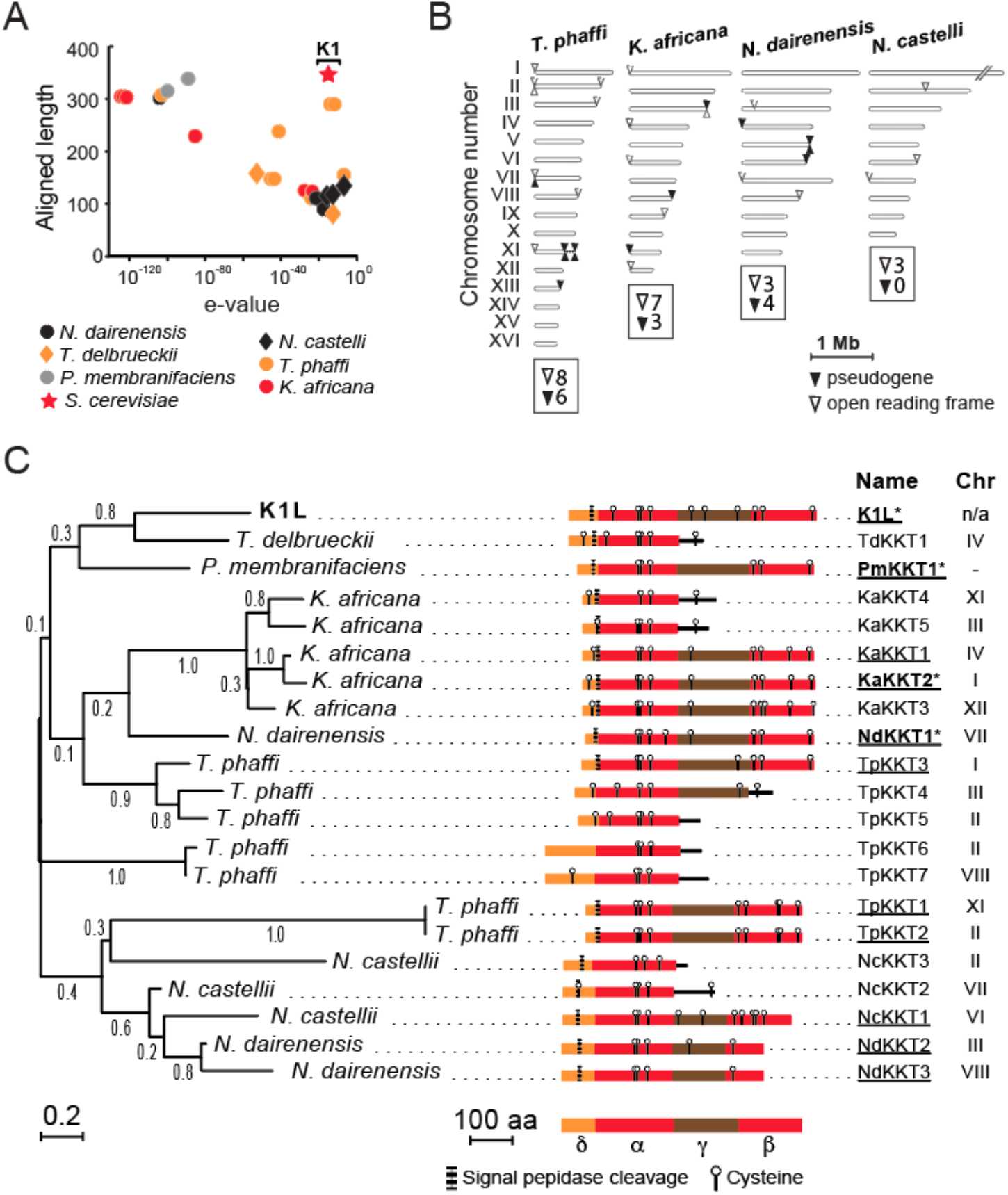
*K1L* genes are broadly distributed across the subphylum Saccharomycotina and are primarily subtelomeric in their genomic location. (A) PsiBLAST analysis of the killer toxin encoded upon the satellite dsRNA from *S. paradoxus* Y-63717. (B) Location of *KKT* genes and pseudogenes on the linear representation of the chromosomes of four yeasts of the *Saccharomycetaceae*. Chromosome I of *N. castellii* did not contain any *KKT* genes and was truncated due to its large size for clarity (C) Unrooted maximum likelihood phylogeny of the aligned a-domain of 21 KKT proteins from six species of yeast and one dsRNA satellite. Numerical values represent the bootstrap support for the placement of each node. The domain organization of each protein is illustrated and annotated based on the four-domain structure of K1. Underlined killer toxin names represent those that were cloned and functionally tested for antifungal activity. *Killer toxins with confirmed antifungal activities.

### K1L Homologs are Found in Yeasts of the Saccharomycotina

The PSI-BLAST search that identified K1L as a homolog of K1 also identified 24 hypothetical “K1-like Killer Toxin” (*KKT*) genes in six diverse species of non-*Saccharomyces* yeasts from the subphylum Saccharomycotina. The species *Kazachstania africana, Naumovozyma castellii, Naumovozyma dairenensis, Tetrapisispora phaffii*, and *Pichia membranifaciens* encode *KKT* genes that closely matched K1L (aligned >300 amino acids, e-value <10^-80^) and represent yeasts from the families *Saccharomycetaceae* and *Pichiaceae* (Fig. 4A and Table S1) [70]. Importantly, genomic *KKT* genes appear to be unique to these particular species and absent from other related yeasts (i.e. *Kazachstania naganishii, Tetrapisispora blattae*, and *Pichia kudriavzevii*). The length of all *KKT* ORFs was between 153-390 amino acids with 11 of the KKT proteins being similar in length to K1L (~340 amino acids) and an amino acid identity between 25-38% (Fig. 4A). In addition, BLASTn was used to identify 14 additional pseudogenes that, in some species, outnumber intact *KKT* genes (Fig. 4B and Table S1). All *KKT* genes and related pseudogenes are found in multiple copies that vary in frequency between different yeast species and are mostly located within subtelomeric regions (within ~20 kb of the assembled chromosome ends) (Fig. 4B). Of the 38 *KKT* genes and pseudogenes found within six different species, only two are positioned away from the subtelomeric regions in the yeasts *N. dairenensis* and *N. castellii* (Fig. 4B). Analysis of the chromosomal position of these two genes revealed that their insertions are unique to each species, are absent from other related species at the syntenic chromosomal location, and are inserted close to tRNAs (Fig. S5). It was noted that six of the *KKT* genes and pseudogenes from *K. africana* contained a characteristic “GAAAAA” sequence motif close to the start codon of each ORF (Fig. S5).

To ascertain the evolutionary history and relatedness of K1L and its homologs, a multiple amino acid sequence alignment was constructed. The most confident alignment was achieved between the putative a-domain of each protein and included eight truncated proteins with premature stop codons. Phylogenetic analysis was performed using maximum likelihood (Fig. 4C) and neighbor-joining (Fig. S6) methodologies with 500 bootstrap iterations as implemented by MEGA [71]. The tree topologies of the two phylogenetic models are in general agreement and show the distinct clustering of KKT killer toxins from each species (Fig. 4C and S6). BLASTn analysis and nucleotide alignment of the 5’ and 3’ UTRs of the *KKT* genes from *K. africana* indicates that flanking nucleotide sequence is between 83-94% identical over ~2,000 bp. There is also evidence of gene duplication of *KKT* within *N. dairenensis* based on their close phylogenetic relationships and similar untranslated regions. The acquisition of *KKT* genes in *T. phaffii* appears to have occurred on three separate occasions resulting in three distinct clades of genes that have significant sequence divergence from each other and other *KKT* genes. Each clade appears to be composed of 2-3 closely related paralogs, suggesting recent gene duplication events of a single ancestral gene. To ascertain the evolutionary trajectory of *KKT* paralogs, the rate of accumulation of nonsynonymous (dN) and synonymous (dS) mutations were calculated for *KKT* paralogs that could be confidently aligned by their nucleotide sequences. When dN/dS was calculated for all codons over the evolutionary history of the sequence pairs, all three pairs of paralogs have evolved under purifying selection since their duplication (dN/dS = 0.72, 0.41, 0.23) [72]. However, a domain-resolution approach using a sliding window to calculate dN/dS indicates evidence of positive selection (dN/dS >1) in the α- and/or g-domains (Fig. S7).

### A New Family of Antifungal Killer Toxins

The relatedness of the *KKT* genes to the killer toxin K1L suggests that they encode antifungal killer toxins. To assay for killer toxin production by the yeasts that encode *KKT* genes, *P. membranifaciens*, *N. dairenensis*, *N. castellii*, *T. delbrueckii*, *T. phaffii*, and *K. africana* were used to challenge 19 different yeast on killer assay plates. With the exception of *N. castellii*, all of the *KKT*-encoding yeast species produced killer toxins that caused growth inhibition of at least one other yeast (Fig. 5A and B). Each of these killer yeasts was also immune to its own killer toxin, but susceptible to those produced by other *KKT*-encoding yeasts. The production of killer toxins by these species is consistent with the previously reported killer activity of *P. membranifaciens*, *T. delbrueckii*, and *T. phaffii* [19,73,74]. There was no evidence of satellite dsRNAs in any of the *KKT*-encoding yeasts, except for the detection of an unknown high molecular weight dsRNA within *P. membranifaciens* NCYC333 (Fig. S8). The differences in killer toxin production by different strains of *P. membranifaciens* indicated that there could be strain-specific differences in *KKT* genes. The published genome sequence of *P. membranifaciens* Y-2026 revealed a large central deletion in the g-domain of its *KKT* gene (Fig. S9). Sanger sequencing of the same *KKT* gene from *P. membranifaciens* Y-2026 acquired directly from the NRRL culture collection failed to identify the same deletion, instead there was an indel within the g-domain that caused the truncation of the killer toxin gene (Fig. S9). Sequencing of the *K1L* gene from *P. membranifaciens* NCYC333 confirmed a full-length *K1L* gene that correlated with robust killer toxin production by the strain. However, Y-2026 was still able to produce killer toxins, suggesting the production of other antifungal molecules by *P. membranifaciens*.

**Fig. 5.**
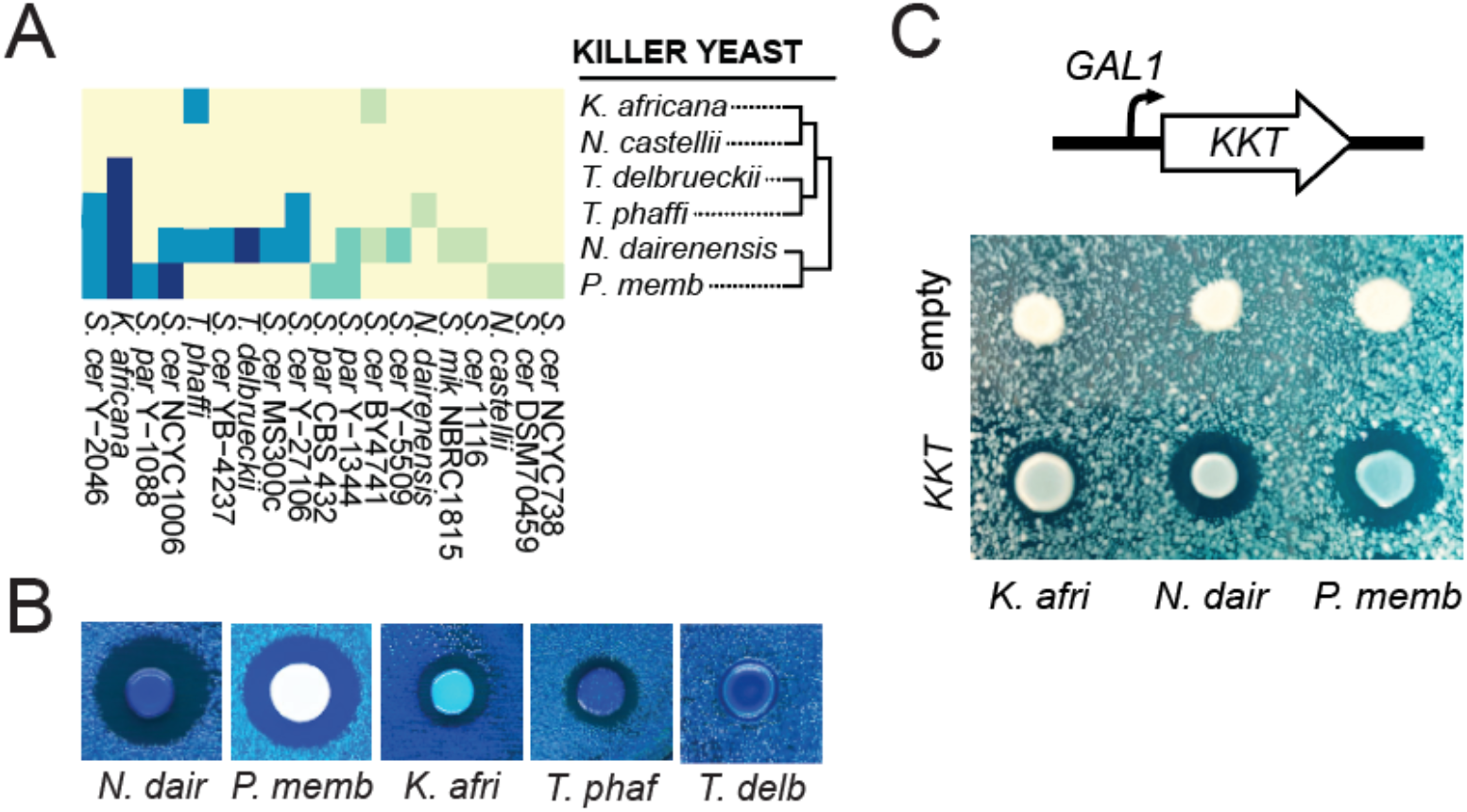
The antifungal activity of *K1L* toxins cloned from *K. africana, N. dairenensis*, and *P. membranifaciens* when ectopically expressed by *S. cerevisiae*. (A) *KKT*-encoding yeasts were assayed for killer toxin production on killer assay agar plates seeded with indicator strains. Killer toxin production was judged by the presence of zones of growth inhibition or methylene blue staining (see Fig. 1). (B) Representative pictures of killer toxin production by different species of yeast from the Saccharomycotina. (C) Galactose-dependent ectopic expression of *KKT* genes from *K. africana* (KaKKT2)*, P. membranifaciens* (PmKKT1) and *N. dairenensis* (NdKKT1) can inhibit the growth of *K. africana*.

Although *P. membranifaciens*, *N. dairenensis*, *T. delbrueckii*, *T. phaffii*, and *K. africana* are killer yeasts (Fig. 5A and B), it was unclear whether *KKT* genes were directly responsible for the observed production of killer toxins. Indeed, *T. phaffii* has been reported to express an antifungal glucanase and K2 killer toxin-related genes have been found in the genome of *K. africana* [68,73]. To demonstrate that *KKT* genes are active killer toxins with antifungal activities, 10 full-length *KKT* genes were cloned into galactose inducible expression vectors (labelled in Fig. 4C).

Active killer toxin production from a non-killer strain of *S. cerevisiae* transformed with *KKT* genes was assayed against 13 lawns of yeasts using galactose containing agar plates. The majority of *KKT* genes did not cause any noticeable growth inhibition or methylene blue staining of competing yeasts (Fig. S10). However, killer toxins from *P. membranifaciens* NCYC333, *K. africana*, and *N. dairenensis* were able to create visible zones of growth inhibition (Fig. 5C). No growth inhibition was observed when the genes were not expressed by plating cells on dextrose (Fig. S10), or when *S. cerevisiae* was transformed with an empty vector control (Fig. 5C). Altogether, these data show that *KKT* genes encode active killer toxins and confirm the discovery of a new family of genome and dsRNA-encoded antifungal proteins in the Saccharomycotina.

## Discussion

The most significant finding of this study is the discovery of a novel satellite dsRNA that encodes a killer toxin related to K1 and a larger family of DNA-encoded homologs in yeasts. The relatedness of killer toxins encoded on dsRNAs and DNAs suggests that the origins of K1L are outside of the *Saccharomyces* genus, with killer toxin gene mobilization and interspecific transfer by dsRNAs. Many of these killer toxins have been shown to be biologically active and are diverse in their amino acid sequences with evidence of their rapid evolution by gene duplication and elevated rates of non-synonymous mutations. This demonstrates the likely benefits of killer toxin acquisition and the ongoing mobilization of these genes between divergent species of yeasts. The more specific implications of our findings are discussed below.

### Horizontal Acquisition and Copy Number Expansion of Killer Toxins in Fungi

KKT genes have most likely been acquired by horizontal gene transfer because of their sporadic distribution and lack of common ancestry in closely related yeast species. Moreover, the lack of relatedness of *KKT* genes in these species suggests that they have independent origins. Fungi are known to acquire foreign DNAs from other species of fungi [75–80] and bacteria [81,82]. The interspecific capture of DNAs derived from retrotransposons, viruses, and plasmids has also been observed [83–86]. Specifically, genome integrated copies of dsRNA-encoded killer toxins homologous to KP4, K1, K2, Klus, and Kbarr have been found within bacteria and fungi, indicating gene flow between dsRNAs and DNAs across taxonomic divisions [68,87]. However, the vast majority of these putative killer toxins are uncharacterized, and it remains unclear as to whether they are biologically active.

Phylogenetic evidence suggests that cross-species transmission of viruses between fungi has occurred on multiple independent occasions between fungi [88,89]. Laboratory experiments have also successfully demonstrated extracellular [90,91] and interspecific virus transfer [27]. In particular for yeasts, mating and hybridization between different species has been frequently observed, and is a mechanism for gene introgression, as well as for the acquisition of retrotransposons and plasmids. The close association of many yeast species in natural and anthropic habitats may increase the likelihood of horizontal gene transfer or invasion by dsRNA viruses and satellites [92–94]. Specifically, the satellite dsRNA that was identified in this study (named SpV-M1L) and an unrelated satellite dsRNA (SpV-M45) are both found within sympatric Far Eastern yeast strains of *S. paradoxus* [27,95]. The parasitism of L-A-45 by both of these satellite dsRNAs in different strains suggests that they were acquired by horizontal gene transfer and evolved distinct mechanisms to enable their replication and packaging by the same totivirus. Unlike *Saccharomyces* yeasts, the presence of active RNAi within the *KKT*-encoding yeasts would hinder the horizontal acquisition of dsRNA viruses from other yeasts, which could explain the abundance of genome encoded killer toxins [96,97]. However, these yeasts can be infected with dsRNA viruses as *T. delbrueckii* can support the replication of the putative totivirus TdV-LAbarr and an associated satellite [19]. Invasion of these species and the evasion of active RNAi by viruses would enable the broad mobilization of killer toxins and viral sequences encoded upon dsRNAs.

To capture killer toxin genes from dsRNAs, the erroneous reverse transcription of mRNAs by endogenous retrotransposons would allow their insertion into the yeast genome by the retrotransposon integrase protein [98]. The ancestor of the *KKT* genes from *K. africana* could have been captured directly from dsRNAs as they all encode a 5’UTR “GAAAAA” motif close to the start codon that is characteristic of a satellite dsRNAs. The 5’ UTR is often conserved during mRNA capture by retrotransposons in addition to a short (20-50 bp) poly(A) tail, which we were unable to identify in the 3’ UTRs of the *KKT* genes from *K. africana* [98]. While the majority of *KKT* genes are subtelomeric in their location, we have observed that non-subtelomeric *KKT* genes from *N. dairenensis* and *N. castellii* have been uniquely inserted into genomic loci near tRNA genes. The genomic integration of retrotransposon cDNAs is selectively targeted to tRNA genes and many extrachromosomal nucleic acids are identified at loci adjacent to tRNAs [85]. This suggests the potential mobilization of these genes by retrotransposons and direct insertion by integrase or cellular DNA repair mechanisms [99]

### The Benefits of Killer Toxin Acquisition

Acquisition of foreign nucleotide sequences can be associated with adaptation to a specific environmental niche, including genes associated with nutrient acquisition, virulence, stress response, and interference competition (e.g. allelopathy) [77,80,85]. The acquisition, expansion, and rapid evolution of *KKT* genes could represent selection for a diverse arsenal of killer toxins to improve competitive fitness. The production of killer toxins by different species of yeasts has been consistently shown to provide a competitive advantage, particularly in a spatially structured environment at an optimal pH for killer toxin activity [100–104]. However, competition between different killer yeasts selects for locally adapted populations that are immune to the predominant killer toxins in a specific environmental niche [7]. Furthermore, laboratory evolution of killer yeast populations has shown that the selective pressure of killer toxin exposure increases the prevalence of killer toxin resistance [105]. The rise of killer toxin resistance within a population would perhaps drive the acquisition of new killer toxins or the subfunctionalization of existing killer toxins to maintain a selective advantage. The majority of *KKT* genes are found within the subtelomeric regions of chromosomes that would facilitate gene expansion due to elevated rates of homologous recombination between telomeric repeat sequences. As has been noted for the subtelomeric *MAL* gene family, gene duplication enables evolutionary innovation that is also evident in *KKT* paralogs by elevated signatures of positive selection [106]. Other fungal killer toxin families have also undergone copy number expansion and are experiencing elevated rates of non-synonymous substitutions [87,107,108]. Both KP4-like and Zymocin-like killer toxin families appear to have roles during antagonistic interactions with plants and fungi, respectively, which could drive the continued evolution of novel killer toxins.

In addition to the expansion of *KKT* genes in yeasts, there is also gene loss and pseudogenization. *KKT* gene inactivation is biased towards truncations that leave the a-domain and a small portion of the g-domain. This same a/g region of K1 is the minimal sequence required for functional killer toxin immunity that would provide a selective advantage by protecting yeasts from exogenous killer toxins related to *KKT* [39,43]. *KKT* yeasts (with the exception of *P. membranifaciens*) all encode C-terminally truncated *KKT* genes and are mostly resistant to other *KKT* killer yeasts. However, despite encoding several truncated *KKT* genes, *K. africana* appears to be naturally susceptible to many of the killer toxins produced by other *KKT* encoding yeasts, including its own killer toxin when ectopically expressed by *S. cerevisiae*. Subtelomeric Sir-dependent gene silencing could account for this apparent susceptibility under laboratory conditions, preventing the constitutive expression of both full-length and truncated *KKT* genes and associated immunity functions by *K. africana* [109,110].

There are now nine evolutionarily distinct dsRNA-encoded killer toxins that have been discovered within *Saccharomyces* yeast. K1L and its homologs represent a unique example of the mobilization and subtelomeric expansion of killer toxin genes in different species. The clear diversity of dsRNAs within *Saccharomyces* yeasts and the known prevalence of killer yeasts suggests that more killer toxins await future discovery and characterization.

## Supporting information

Fie S3

## Acknowledgements

We would like to acknowledge the NRRL Agricultural Research Service culture collection for providing the Rowley laboratory with a diverse collection of *Saccharomyces* yeasts. We would also like to thank Dr. Antonia Santos for the CYC yeasts (Complutense Yeast Collection, Complutense University of Madrid), Prof. Manfred J. Schmitt (Saarland University, Saarbrücken, Germany) for *S. cerevisiae* MS300c, and Dr. Gianni Liti for the genetically tractable strains of *S. paradoxus* and *S. cerevisiae*. We would also like to acknowledge Dr. Gianni Liti and Dr. Scott Minnich for helpful comments on the draft manuscript.

## SUPPLEMENTARY FILES

**Fig. S1.**
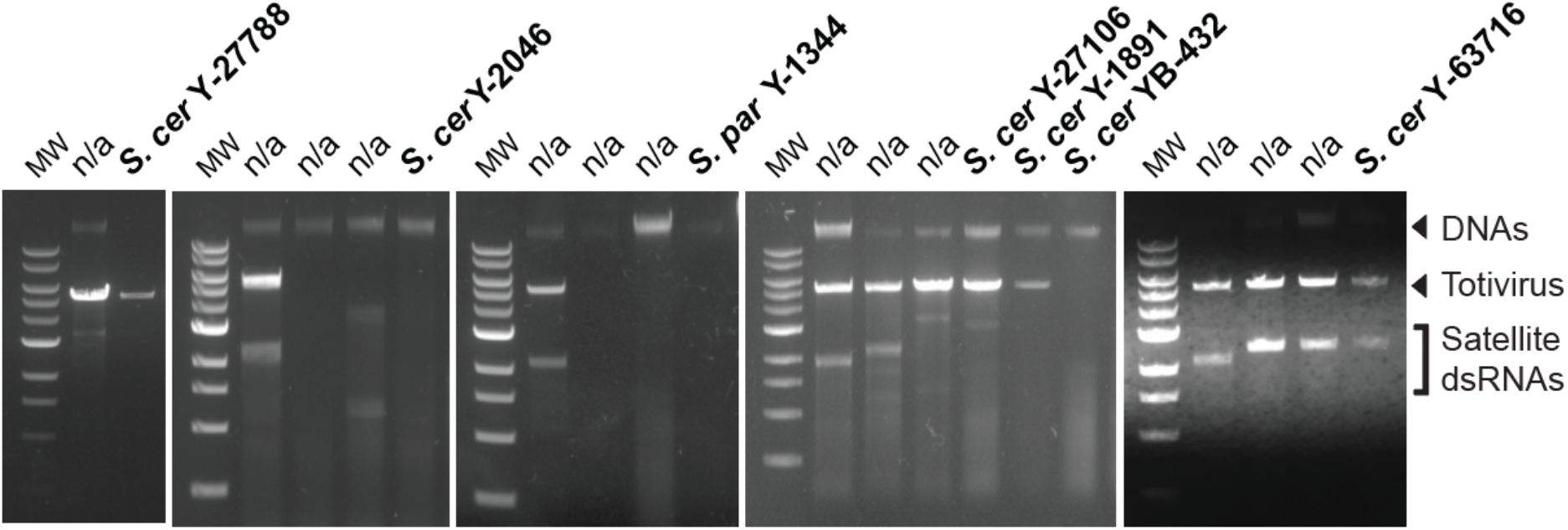
Double-stranded RNA enrichment and analysis of nucleic acids from killer yeasts. Agarose gel electrophoresis is used to show the diversity of dsRNA in killer yeasts. Satellite dsRNAs are labeled as dsRNAs that are smaller than the associated totivirus dsRNAs.

**Fig. S2.**
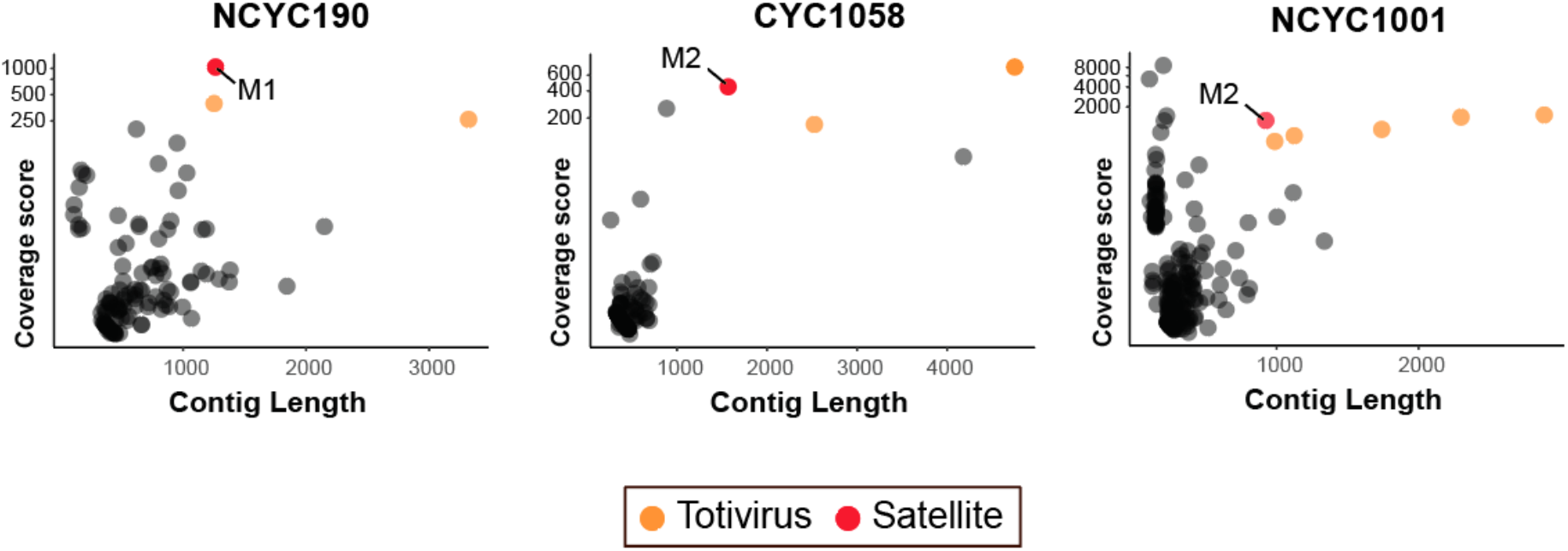
Coverage and contig length of NGS of dsRNAs from different strains of yeasts. Each scatter plot represents all contigs generated after *de novo* assembly of sequence reads after assembly. M satellites are labeled according to their relatedness to other previously described sequences.

**Fig. S3.**
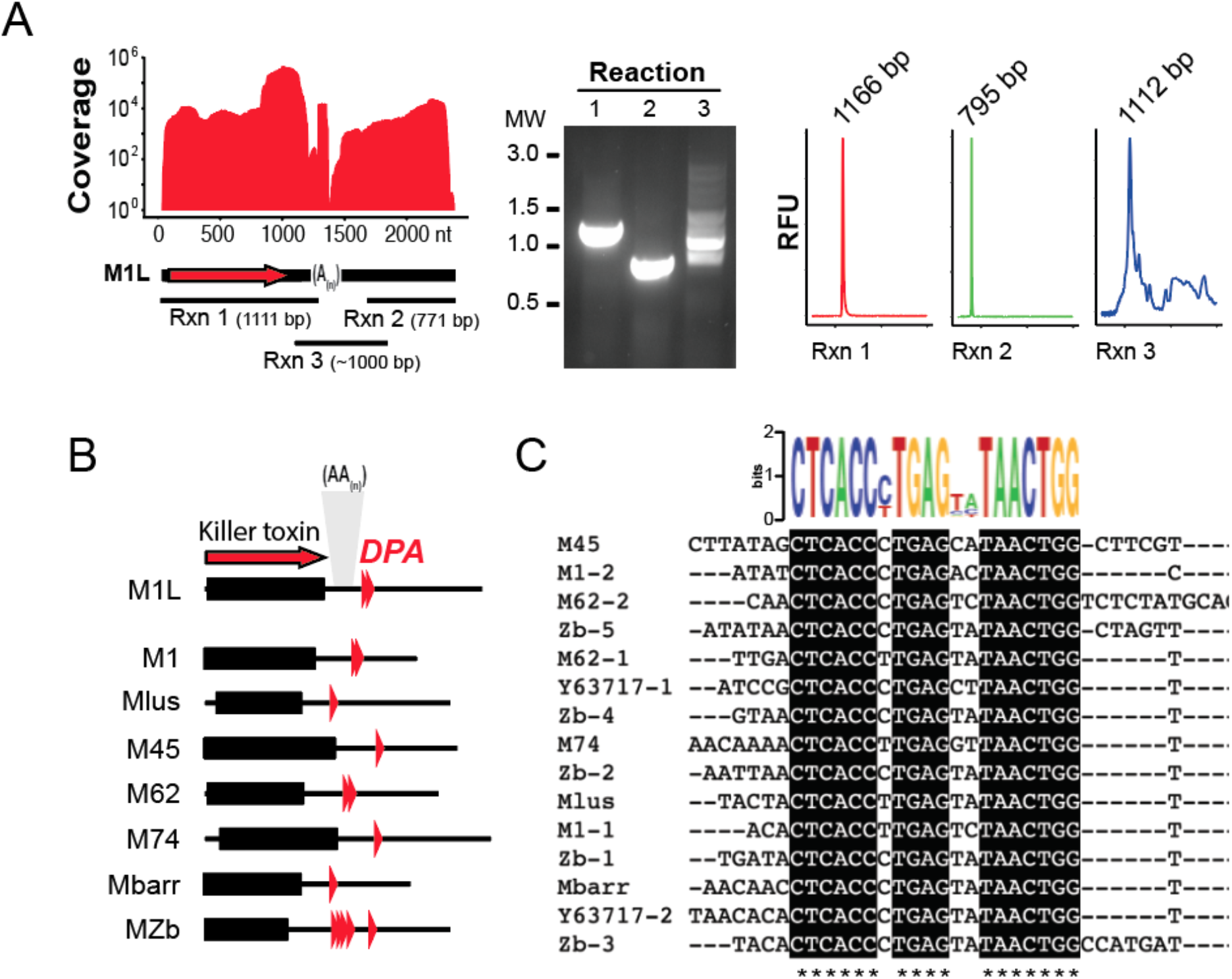
Sequence analysis of the dsRNA satellite M1L from *S. paradoxus*. (A) Coverage of the assembled short reads for the assembly of M1L and the positioning the expected product from three reverse transcription PCR reactions to amplify portions of K1L ORF, the 3’ UTR, and across the internal poly(A) tract. Actual PCR products and their estimated sizes as determined from fragment analysis are shown. (B) The positioning of the repeated *DPA* element is represented relative to the genomes of eight dsRNA satellites. (C) The consensus sequence derived from 15 *DPA* elements is shown as a sequence logo and multisequence alignment.

**Fig. S4.**
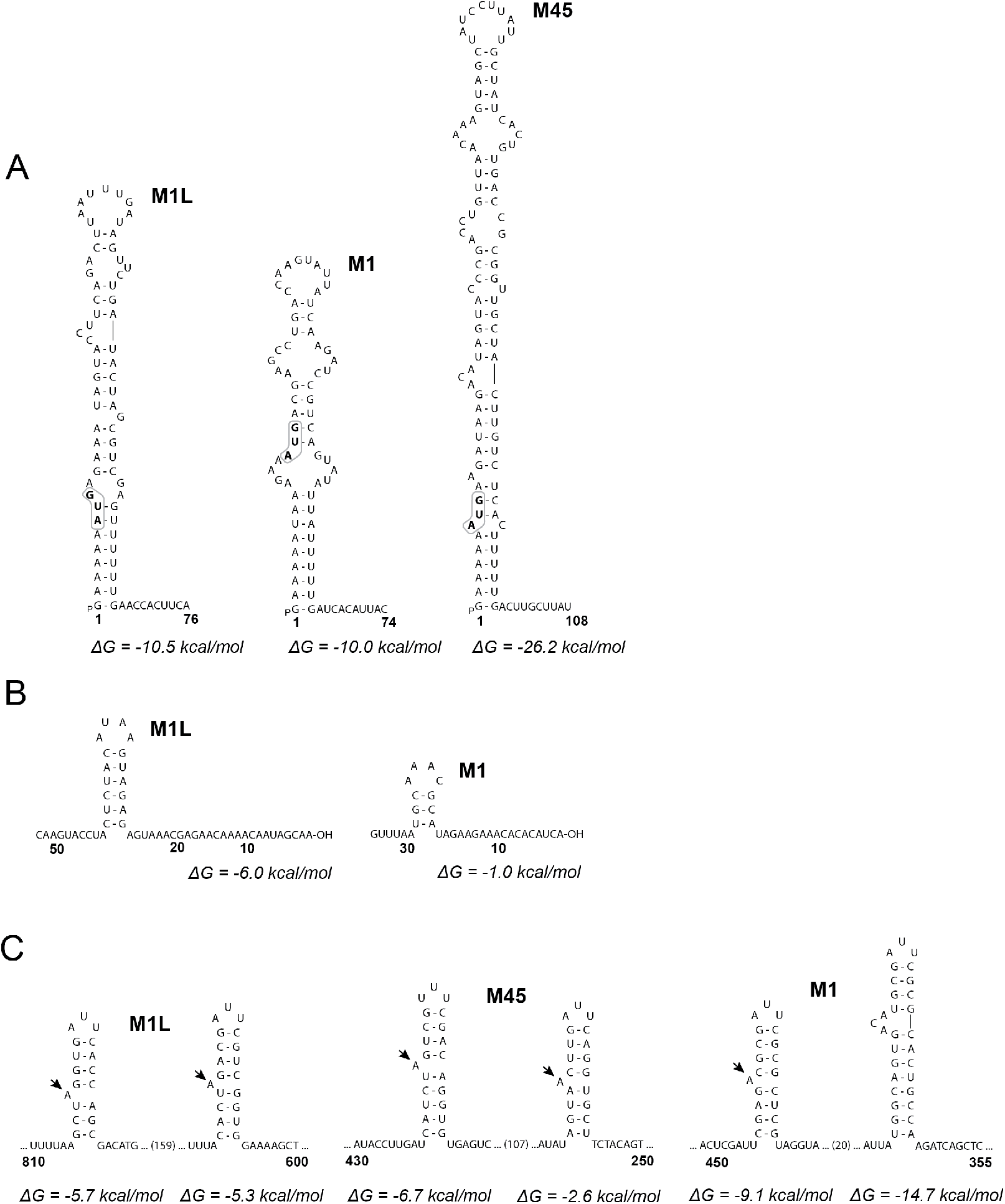
Secondary structure predictions of M1L, M1, and M45 (+) strand 5’ and 3’ ends. (A) Secondary prediction of the 5’ terminal structures. Start codons for the translation of preprotoxin synthesis are highlighted. Numbers represent nucleotides from the 5’ terminus. (B) Putative replication signal represented as a stem-loop at the 3’ end of M1L and M1 satellite. Numbers represent distance from the 3’ terminal nucleotide. (C) Putative viral particle binding sites with a 5’ facing ‘A’ bulge present in the stem-loops (indicated by an arrow). Numbers represent distance from the 3’ terminal nucleotide.

**Fig. S5.**
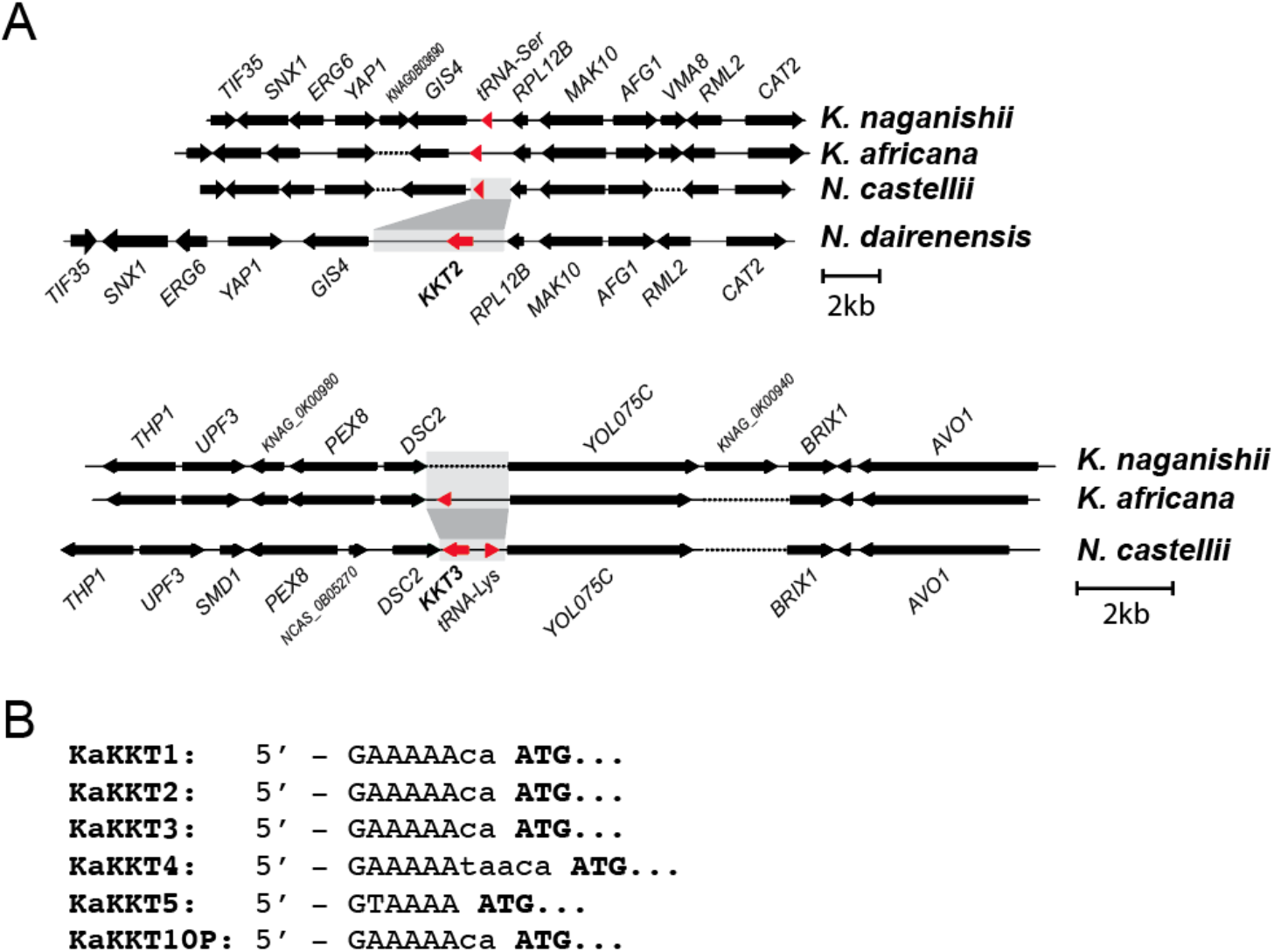
Unique genomic insertions of *KKT* genes in *N. dairenensis* and *N. castellii*. (A) *N. dairenensis KKT2* inserted into chromosome III and *N. castellii KKT3* inserted into chromosome II. Genes flanking *KKT* insertions are colored green and demonstrate synteny between related genomes. Single red triangles represent tRNA genes. Brocken lines represent gaps in synteny that were inserted for clarity. (B) 5’ UTR sequence from *KKT* genes and one pseudogene identified within *K. africana*.

**Fig. S6.**
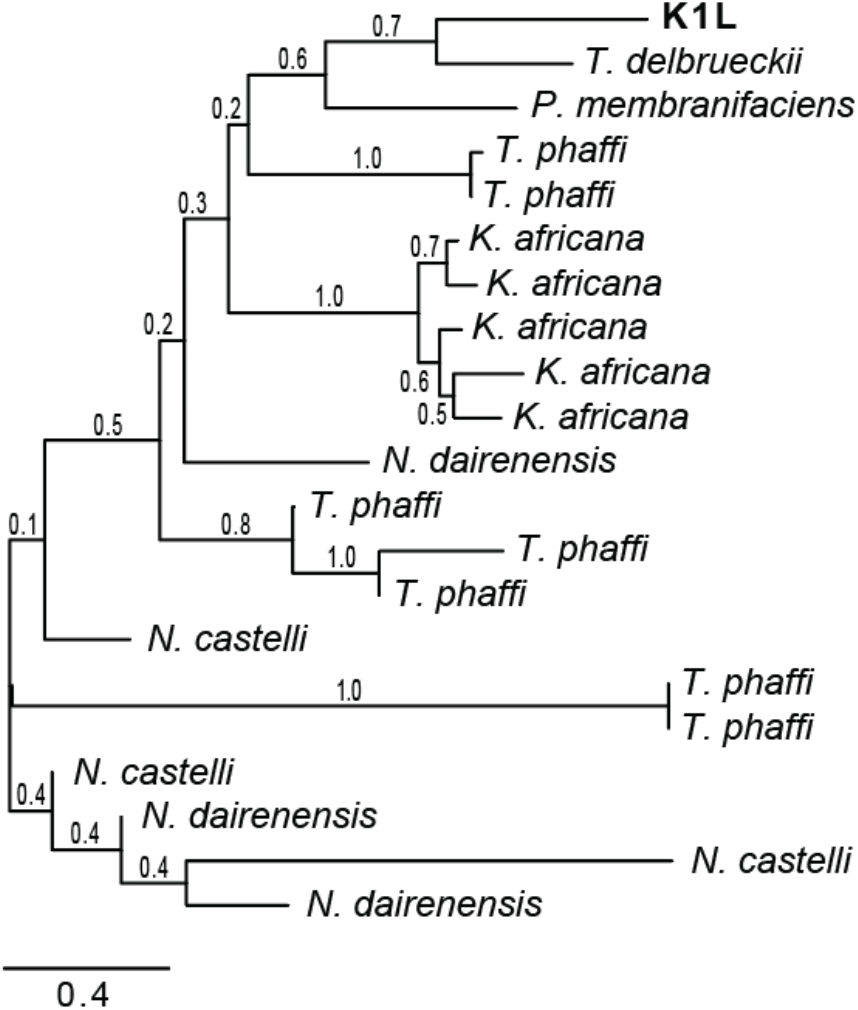
Phylogenetic model of the evolutionary relationship between *KKT* proteins using the neighbor-joining method. Unrooted neighbor-joining phylogeny of the aligned a-domain of 21 *KKT* proteins from six species of yeast and one dsRNA satellite dsRNA. Numerical values represent the bootstrap support for the placement of each node.

**Fig. S7.**
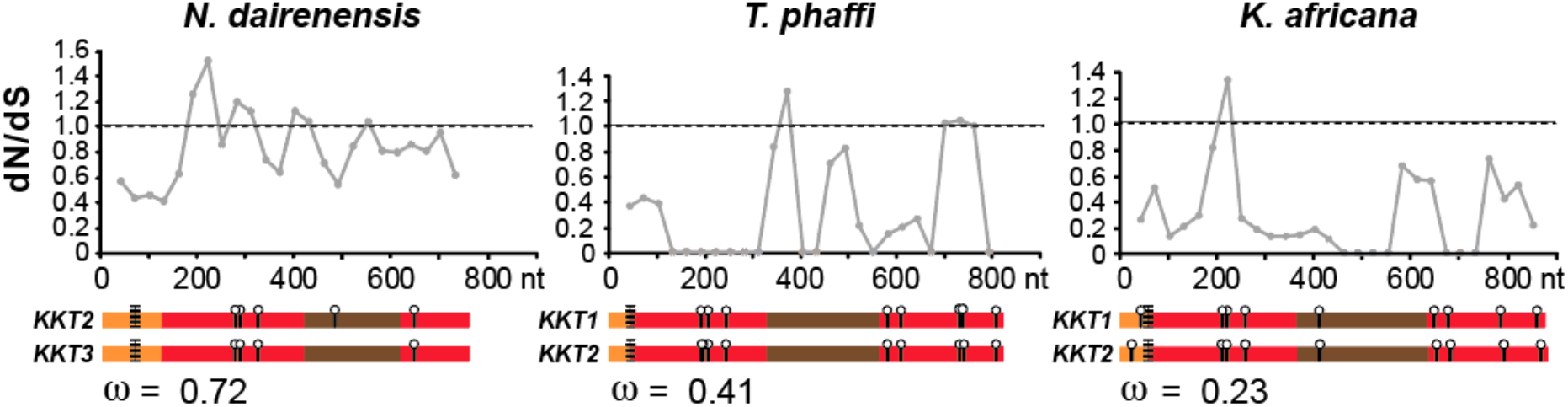
Paralogous *KKT* genes are evolving under positive selection in different species of yeasts. Three sliding window dN/dS calculations are shown for the comparison of three pairs of closely related genes in three yeast species. The X-axis represents the nucleotide (nt) number of each gene and is shown in the context of the predicted domain organization of each pair of genes. Omega values represent the whole gene dN/dS value for each gene pair.

**Fig. S8.**
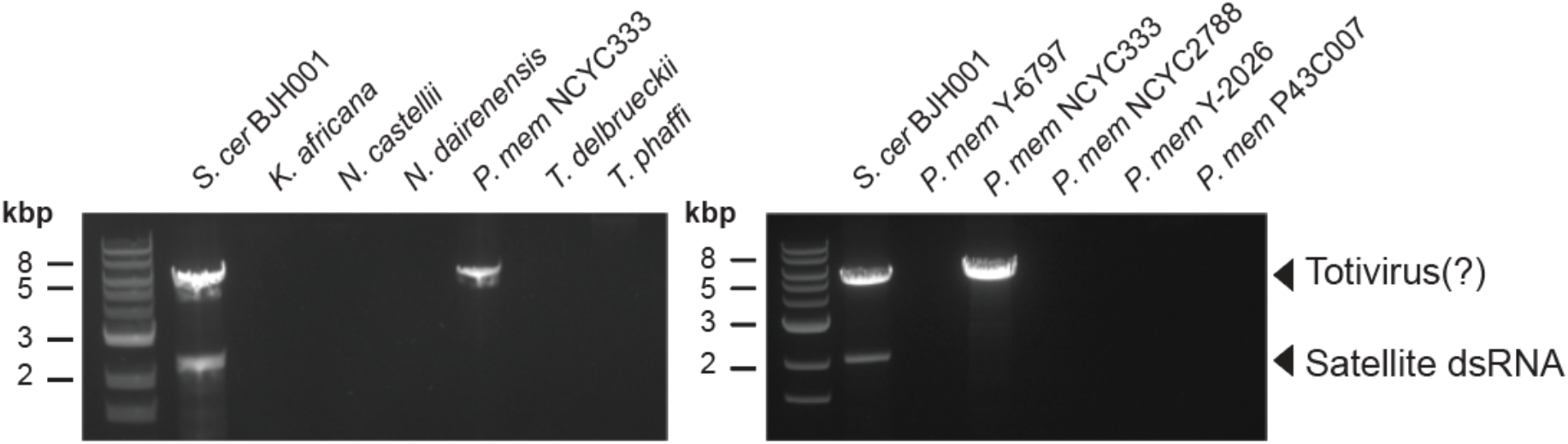
The absence of satellite dsRNAs within killer yeasts of the Saccharomycotina. Agarose gel electrophoresis of dsRNAs extracted from different killer yeasts. Stained bands in lane 2 represent canonical totivirus and satellite dsRNAs from *S. cerevisiae*.

**Fig. S9.**
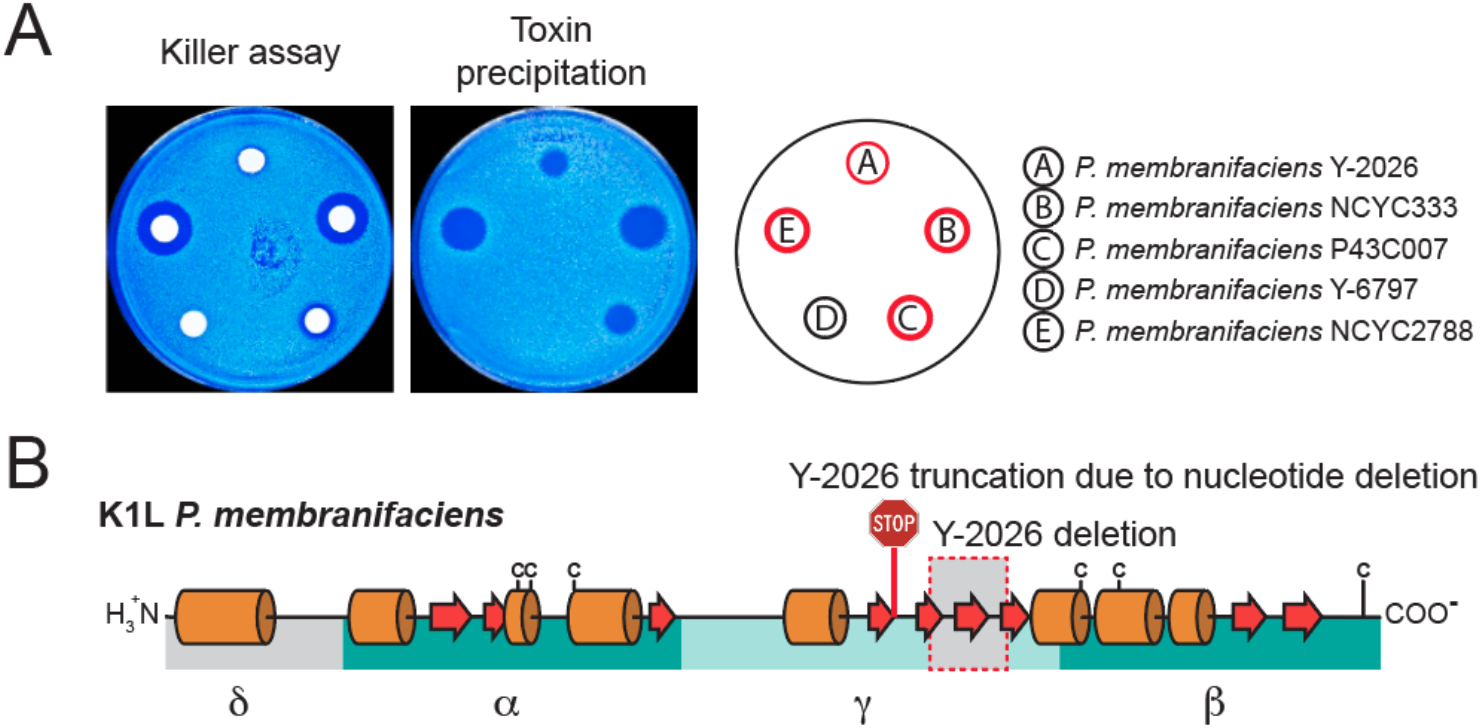
Strain-specific production of killer toxins by *P. membranifaciens*. (A) Killer toxin production and partial purification from *P. membranifaciens* (B) Mutations within *KKT1* in the context of the proteins secondary structure organization (as predicted by Jpred) from strain Y-2026 compared to full-length active killer toxin sequenced from strain NCYC333. Arrows represent beta-sheets and cylinders represent alpha helices. “c” represents cysteine residues.

**Fig. S10.**
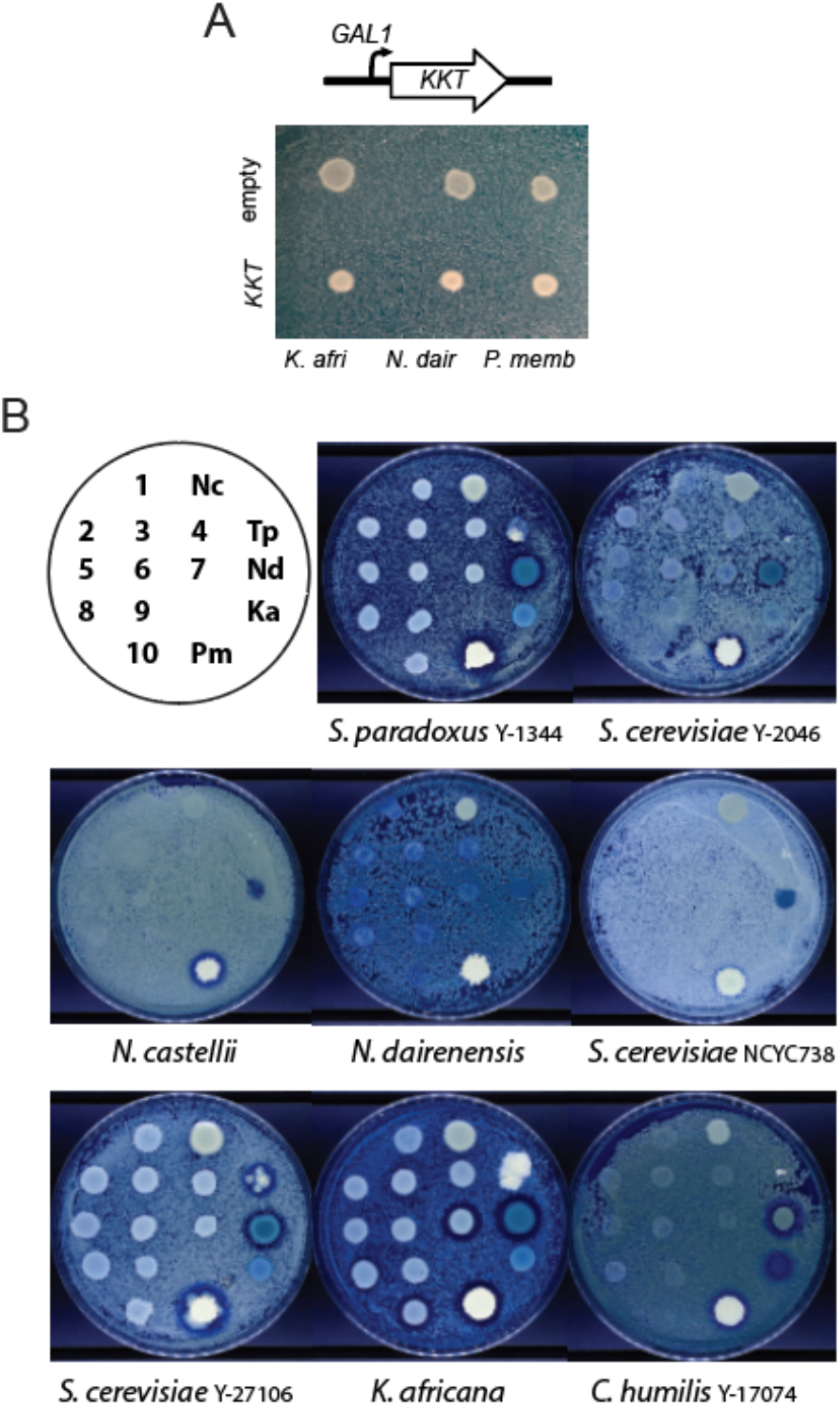
Ectopic expression of KKT genes causes the inhibition of K. africana and is dependent on the induction of KKT expression. (A) Galactose-dependent ectopic expression of *KKT* genes by *S. cerevisiae* on agar plates seeded with different species of yeasts. Key: 1. pUI114, 2. pUI109 (T.p1), 3-pUI110 (T.p2), 4-pUI111 (T.p3), 5. pUI112 (N.d1), 6-pUI113 (N.d2), 7-pML115 (N.d3), 8. pML117 (K.a1), 9. pML118 (K.a2), 10. pML116 (P.m), Nc. *N. castellii* NCYC2898, Nd. *N. dairenensis* NCYC777, Tp. *T. phaffii* Y-8282, Pm. *P. membranifaciens* NCYC333.

**Table S1.** Species that encode genome encoded killer toxins that are homologous to K1L.

| Species | Strain | Gene | Protein | Chr | Position | nt | aa |
| --- | --- | --- | --- | --- | --- | --- | --- |
| *K. africana* | CBS 2517 | KaKKT1 | XP_003956791.1 | 4 | 4933-5865 | 932 | 310 |
|  |  | KaKKT3 | XP_003959753.1 | 12 | 3868-4800 | 932 | 310 |
|  |  | KaKKT2 | XP_003954584.1 | 1 | 14695-15627 | 932 | 310 |
|  |  | KaKKT5 | XP_003956787.1 | 3 | 1322627-1323511 | 884 | 153 |
|  |  | KaKKT4 | XP_003959210.1 | 9 | c592072-591575 | 885 | 165 |
|  |  | KaKKT6 | XP_003957738.1 | 6 | 4855-5505 | 650 | 216 |
|  |  | KaKKT7P | - | 8 | 738788-737853 | 935 | - |
|  |  | KaKKT8P | - | 3 | 1320455-1320080 | 375 | - |
|  |  | KaKKT9P | - | 11 | 29171-30101 | 930 | 95 |
| *N. castellii* | CBS 4309 | NcKKT1 | XP_003677247.1 | 6 | 820175-821086 | 911 | 303 |
|  |  | NcKKT2 | XP_003677251.1 | 7 | 4472-5032 | 560 | 186 |
|  |  | NcKKT3 | XP_003674985.1 | 2 | 979503-979970 | 467 | 155 |
| *N. dairenensis* | CBS 421 | NdKKT1 | XP_003671029.2 | 7 | 263-1186 | 923 | 307 |
|  |  | NdKKT2 | XP_003669012.1 | 3 | 214531-215346 | 815 | 271 |
|  |  | NdKKT3 | XP_003671820.1 | 8 | 997352-998167 | 815 | 271 |
|  |  | NdKKT4P | - | 5 | 1179525-1178607 | 918 | - |
|  |  | NdKKT5P | - | 6 | 1136398-1135718 | 680 | - |
|  |  | NdKKT7P | - | 6 | 1135727-1135598 | 129 | - |
| *T. phaffii* | CBS 4417 | TpKKT3 | XP_003683531.1 | 1 | 8390-9322 | 932 | 310 |
|  |  | TpKKT1 | XP_003687580.1 | 11 | 11192-10317 | 875 | 291 |
|  |  | TpKKT5 | XP_003684118.1 | 2 | 7999-8806 | 807 | 161 |
|  |  | TpKKT6 | XP_003684596.1 | 2 | 1153004-1153633 | 629 | 209 |
|  |  | TpKKT7 | XP_003686950.1 | 8 | 749558-750160 | 602 | 200 |
|  |  | TpKKT2 | XP_003684117.1 | 2 | 1052-1927 | 875 | 291 |
|  |  | TpKKT4 | XP_003685085.1 | 3 | 1089707-1090513 | 806 | 268 |
|  |  | TpKKT8 | XP_003686288.1 | 7 | 6094-5576 | 518 | 172 |
|  |  | TpKKT12P | - | 11 | 512438-515245 | 807 | - |
|  |  | TpKKT9P | - | 11 | 521694-523935 | 241 | - |
|  |  | TpKKT10P | - | 11 | 518825-521023 | 198 | - |
|  |  | TpKKT11P | - | 13 | 451100-453480 | 380 | - |
|  |  | TpKKT13P | - | 11 | 520515-520628 | 113 | - |
|  |  | TpKKT13P | - | 11 | 520515-520628 | 113 | - |
|  |  | TpKKT14P | - | 7 | 3752-3575 | 177 | - |
| *T. delbrueckii* | CBS 1146 | TdKKT1 | XP_003680807.1 | 4 | 11668-12189 | 522 | 173 |
| *P. membranifaciens* | KS47-1 | PmKKT1 | GAV30688.1 | n/a | c2248-1259 | 990 | 390 |

**File S1. A large-scale screen to identify killer yeasts in the *Saccharomyces* genus.**

**File S2. Image data illustrating the susceptibility of 53 strains of yeast to a selection of potent killer toxins produced by *Saccharomyces* yeasts.**

**File S3. SignalP and TargetP predictions for K1, K1L and KKT proteins.**

**File S4. Supplementary file listing all primers, plasmids, and yeast strains used in this study.**

**File S5. Supplementary FASTA file with the DNA sequences of all plasmids used in this study.**

