## Supplementary material for "The Species-specific Acquisition and Diversification of a Novel Family of Killer Toxins in Budding Yeasts of the Saccharomycotina": File S1

**File S1. Image data of killer phenotypes exhibited by strains of *Saccharomyces* yeasts as summarized in figure 1.** The organization of a large collection of strains of *Saccharomyces* yeasts arrayed on killer assay agar plates. Each killer yeast was assayed for toxin activity against 8 different lawns of susceptible yeast strains. Black squares indicate detectable killer toxin production.

| | Sensitive Lawns | | | | | | | |
|---|---|---|---|---|---|---|---|---|
| Yeasts Tested for Killer Phenotype | K12 | FY4 | BY4741 | UWOPS 83-787.3 | CBS 432 | CBS7001 | SSS 104 | NCYC 2729 |
| Y-580 | | | | | | | | |
| Y-670 | | | | | | | | ■ |
| Y-846 | | | | | | | | |
| Y-969 | | | | | | | | |
| Y-972 | | | | | | | | |
| Y-1374 | | | | | | | | |
| Y-11845 | | | | | | | | |
| Y-12624 | | | | | | | | |
| Y-12646 | | | | | | | | |
| Y-12648 | | | | | | | | |
| Y-17034 | | | | | | | | |
| Y-27339 | | | | | | | | |
| Y-27470 | | | | | | | | |
| Y-48770 | | | | | | | | |
| Y-63707 | | | | | | | | |
| Y-63718 | | | | | | | | |
| YB-254 | | | | | | | | |
| YB-432 | | | | | | | | ■ |
| Y-567 | | | | | | | | |
| Y-851 | | | | | | | | |
| Y-852 | | | | | | | | |
| Y-897 | | | | | | | | |
| Y-898 | | | | | | | | |
| YB-908 | | | | | | | | ■ |
| Y-954 | | | | | | | | |
| Y-975 | | | | | | | | |
| Y-976 | | | | | | | | |
| Y-977 | | | | | | | | ■ |
| Y-1018 | | | | | | | | |
| Y-1089 | | | | | | | | |
| Y-1285 | | | | | | | | |
| Y-1301 | | | | | | | | |
| Y-1370 | | | | | | | | |
| Y-1428 | | | | | | | | |
| Y-1429 | | | | | | | | |
| Y-1430 | | | | | | | | |
| Y-1436 | | | | | | | | |
| Y-1438 | | | ■ | | | | | ■ |
| Y-1536 | | | | | | | | |
| Y-1540 | | | | | | | | |
| YB-1773 | | | | | | | | |
| Y-1891 | | | ■ | | | | | ■ |
| Y-2044 | | | | | | | | |
| Y-2045 | | | | | | | | |
| Y-2046 | | | ■ | | | | | |

| | | | | | | | | | |
|---|---|---|---|---|---|---|---|---|---|
| | Y-2204 | | | | | | | | |
| | Y-2205 | | | | | | | | |
| | Y-2429 | ■ | | ■ | | | | | ■ |
| | Y-2430 | | | | | | | | |
| | Y-2432 | | | | | | | | |
| | Y-2434 | | | | | | | | |
| | YB-4237 | | | ■ | | | | | |
| | YB-4255 | | | | | | | | |
| | YB-4634 | | | | | | | | |
| | YB-4635 | | | | | | | | |
| | Y-5508 | | | | | | | | |
| | Y-5509 | ■ | | ■ | | | | | |
| | Y-5510 | | | | | | | | |
| | Y-7327 | | | | | | | | |
| | Y-7328 | | | | | | | | |
| | Y-7567 | | | | | | | | |
| | Y-10988 | | | | | | | | |
| | Y-11875 | | | | | | | | |
| | Y-12842 | | | | | | | | |
| | Y-17009 | | | | | | | | ■ |
| | Y-17898 | | | ■ | | | | | |
| | Y-27105 | ■ | | | | | | | |
| | Y-27106 | | | | | | | ■ | |
| | Y-27437 | | | | | | | | |
| | Y-27788 | | | ■ | | | | | |
| | Y-27796 | | | | | | | | |
| | y-63703 | | | | | | | | |
| | Y-63748 | | | | | | | | |
| | Y-63749 | | | | | | | | |
| | Y-27340 | | | | | | | | |
| | Y-27341 | | | | | | | | |
| | Y-27342 | | | ■ | | | | | |
| | Y-27471 | | | | | | | | |
| | Y-63704 | | | ■ | | | | | |
| | Y-63705 | | | | | | | | |
| | Y-63706 | | | | | | | | |
| | Y-788 | | | | | | | | |
| | Y-863 | | | | | | | | |
| | Y-911 | | | | | | | | |
| | Y-1088 | ■ | | ■ | | | | ■ | |
| | Y-1344 | | | ■ | | | | | |
| | Y-1356 | | | | | | | | |
| | Y-1548 | | | | | | | | |
| | Y-1912 | | | | | | | | |
| | Y-2038 | | | | | | | | |
| | YB-2047 | | | | | | | | |
| | YB-4137 | | | | | | | | |
| | YB-4565 | | | ■ | | | | | |
| | Y-5688 | | | | | | | | |
| | Y-6177 | | | | | | | | |
| | Y-6179 | | | | | | | | |
| | Y-11842 | | | | | | | | |
| | Y-12602 | | | ■ | | | | | |
| | Y-17218 | | | | | | | | |
| | Y-17353 | | | | | | | | |
| | Y-63708 | | | | | | | | |
| | Y-63709 | | | | | | | | |
| | Y-63710 | | | | | | | | |
| | Y-63711 | | | ■ | | | | ■ | |
| | Y-63712 | | | | | | | | |

| | | | | | | | | | |
|---|---|---|---|---|---|---|---|---|---|
| | Y-63713 | | | | | | | | |
| | Y-63714 | | | | | | | | |
| | Y-63715 | | | ■ | | | | ■ | ■ |
| | Y-63716 | ■ | | | | | | | ■ |
| | Y-63717 | | | ■ | | | | | ■ |

Key

<table>
<tr><td colspan="6">Y-1436</td></tr>
<tr><td colspan="3">Y-2045</td><td colspan="3">Y-1438</td></tr>
<tr><td colspan="2">Y-2044</td><td colspan="2">Y-2046</td><td colspan="2">Y-1536</td></tr>
<tr><td colspan="3">Y-1891</td><td colspan="3">Y-1540</td></tr>
<tr><td colspan="6">YB-1773</td></tr>
</table>

BY4741

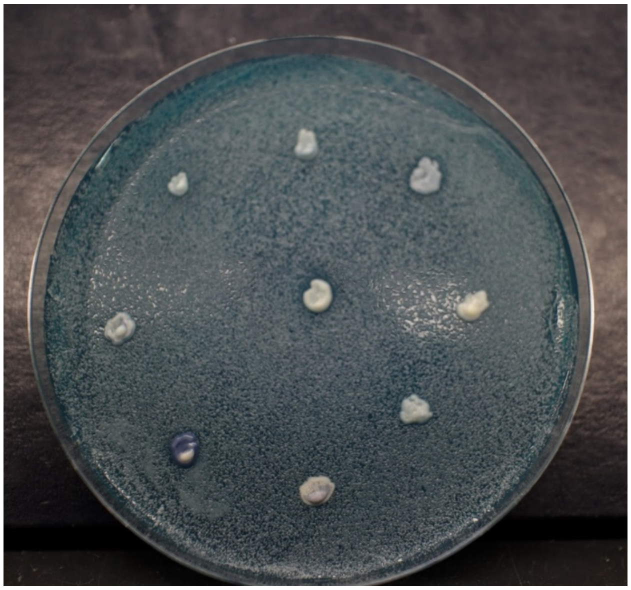

NCYC 2729

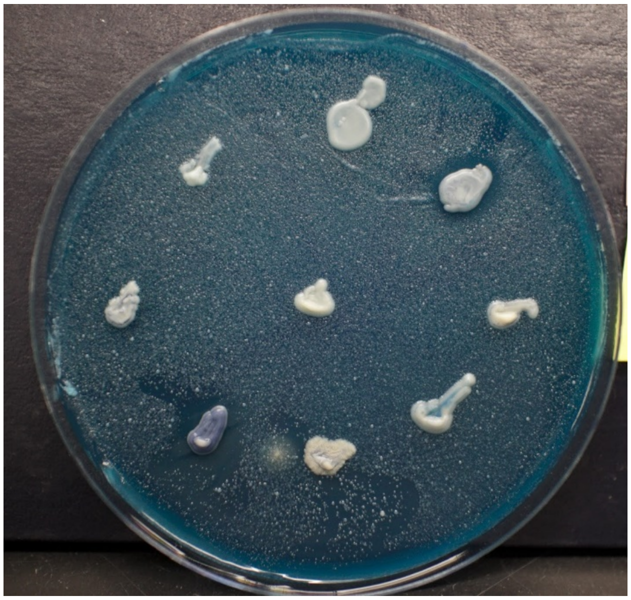

Key

<table>
<tr><td colspan="6">Y-2204</td></tr>
<tr><td colspan="3">YB-4255</td><td colspan="3">Y-2205</td></tr>
<tr><td colspan="2">YB-4237</td><td colspan="2">YB-4634</td><td colspan="2">Y-2429</td></tr>
<tr><td colspan="3">Y-2434</td><td colspan="3">Y-2430</td></tr>
<tr><td colspan="6">Y-2432</td></tr>
</table>

K 12

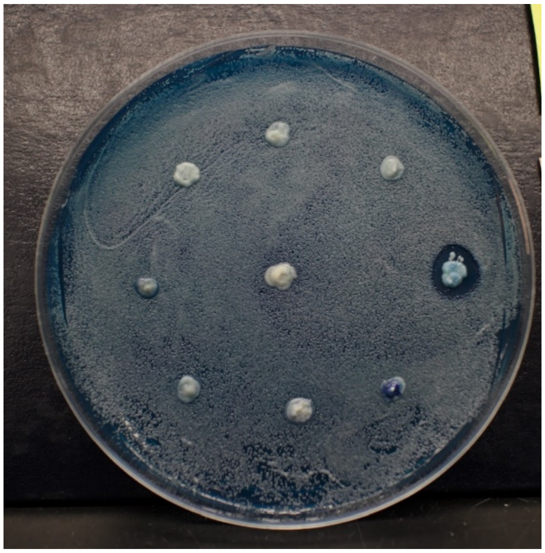

BY4741

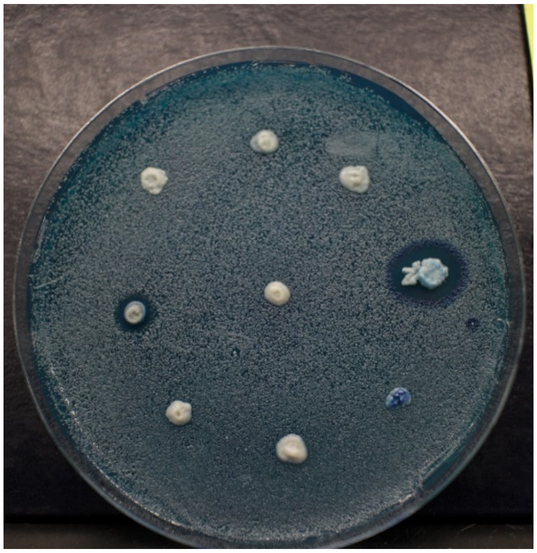

SSS 104

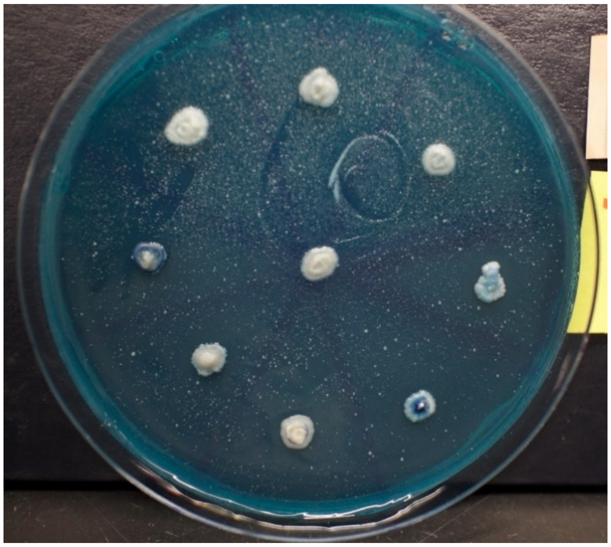

NCYC 2729

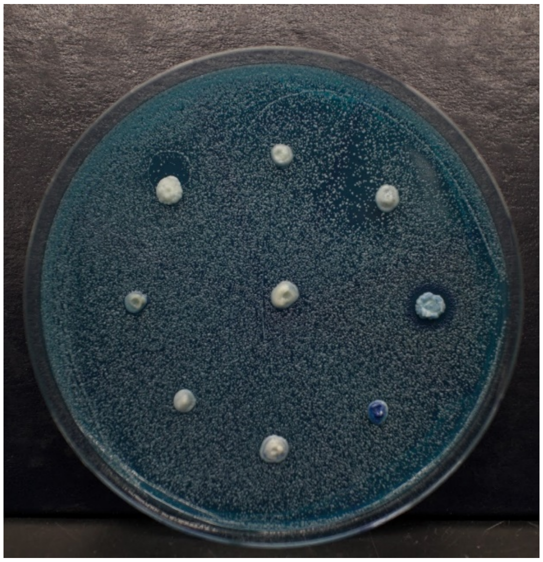

Key

<table>
<tr><td colspan="6">YB-4635</td></tr>
<tr><td colspan="3">Y-10988</td><td colspan="3">Y-5508</td></tr>
<tr><td colspan="2">Y-7567</td><td colspan="2">Y-11875</td><td colspan="2">Y-5509</td></tr>
<tr><td colspan="3">Y-7328</td><td colspan="3">Y-5510</td></tr>
<tr><td colspan="6">Y-7327</td></tr>
</table>

K12

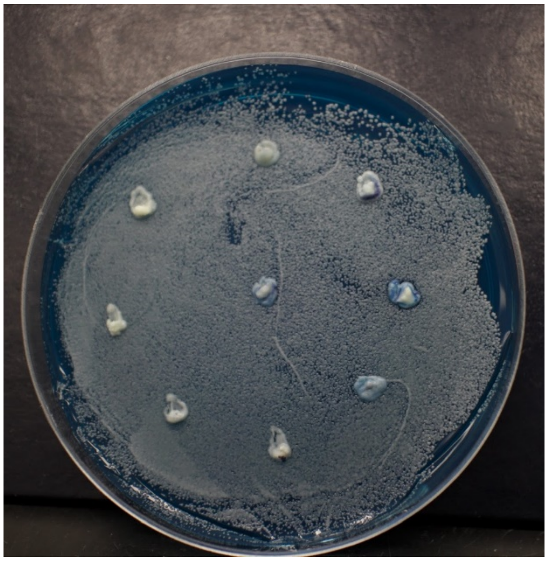

BY4741

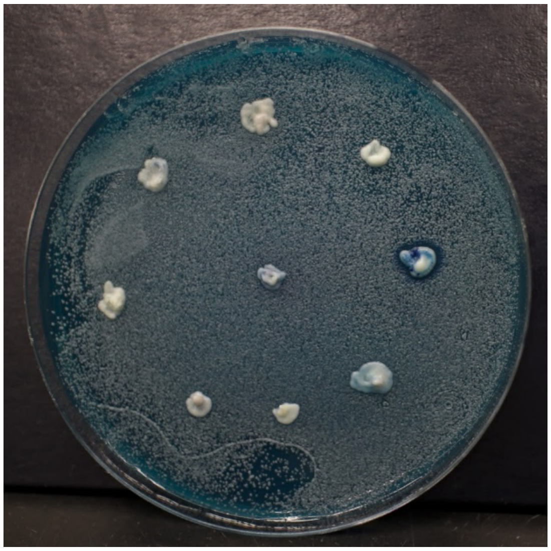

Key

<table>
<tr><td colspan="6">Y-12842</td></tr>
<tr><td colspan="3">Y-27796</td><td colspan="3">Y-17009</td></tr>
<tr><td colspan="2">Y-27788</td><td colspan="2">Y-63703</td><td colspan="2">Y-17898</td></tr>
<tr><td colspan="3">Y-27437</td><td colspan="3">Y-27105</td></tr>
<tr><td colspan="6">Y-27106</td></tr>
</table>

K12

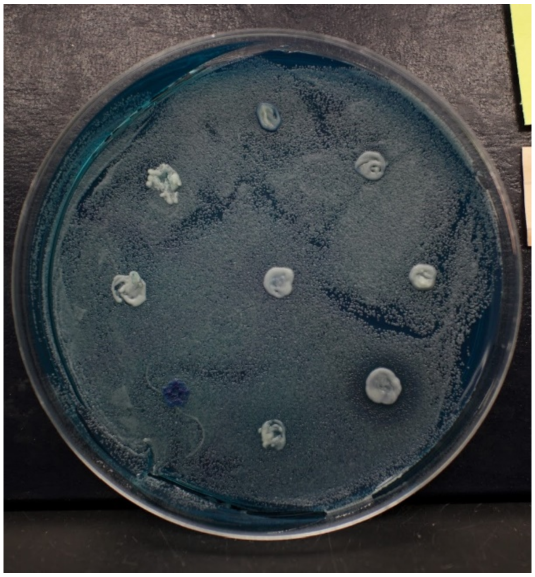

SSS 104

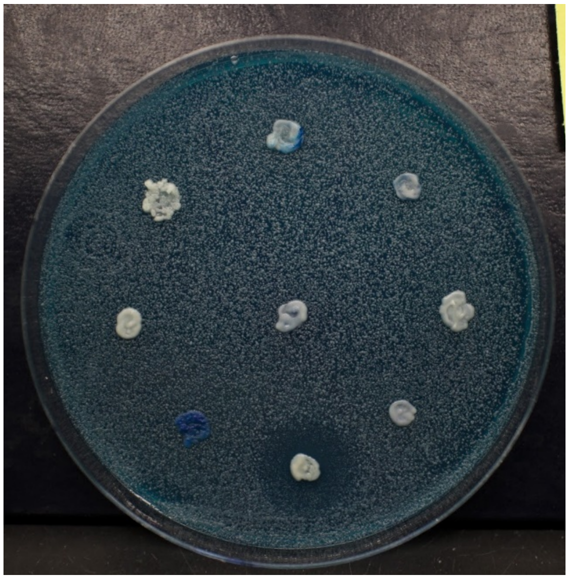

NCYC 2729

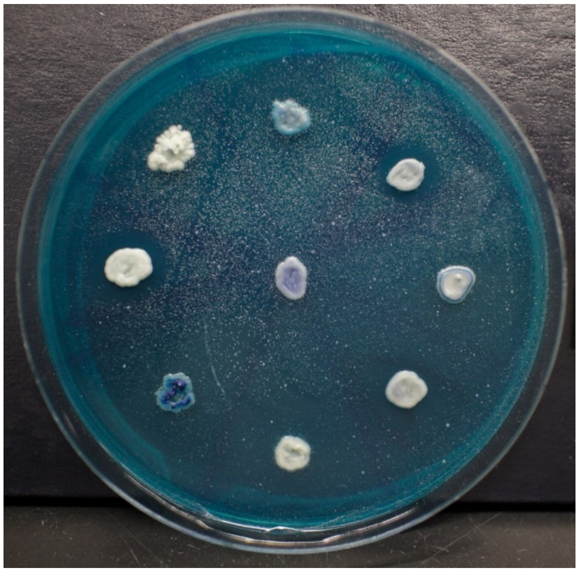

BY4741

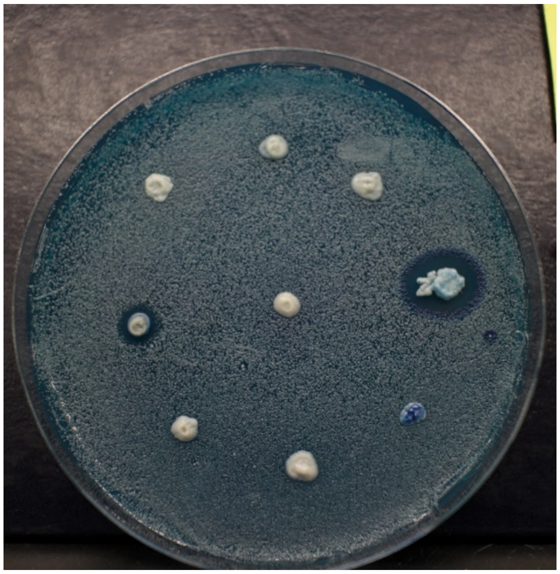

Key

| | | | | |
|---|---|---|---|---|
| Y-63748 | | | | |
| Y-63705 | | | Y-63749 | |
| **Y-63704** | | Y-63706 | | Y-27340 |
| Y-27471 | | | Y-27341 | |
| **Y-27342** | | | | |

BY4741

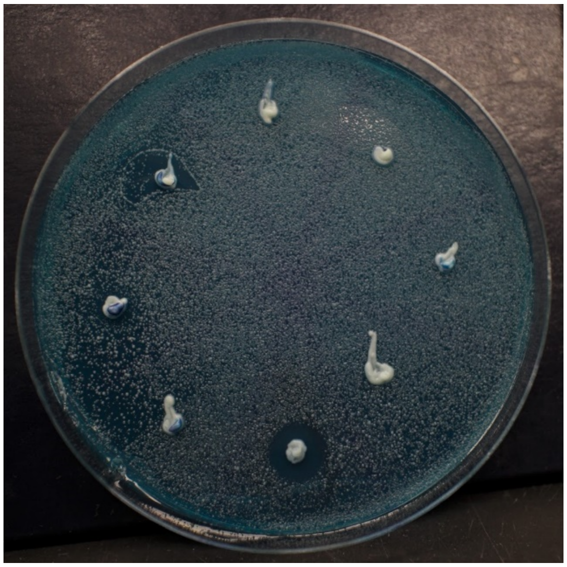

Key

<table>
<tr><td colspan="6">Y-788</td></tr>
<tr><td colspan="3">Y-1912</td><td colspan="3">Y-863</td></tr>
<tr><td colspan="2">Y-1548</td><td colspan="2">Y-2038</td><td colspan="2">Y-911</td></tr>
<tr><td colspan="3">Y-1356</td><td colspan="3">Y-1088</td></tr>
<tr><td colspan="6">Y-1344</td></tr>
</table>

K12

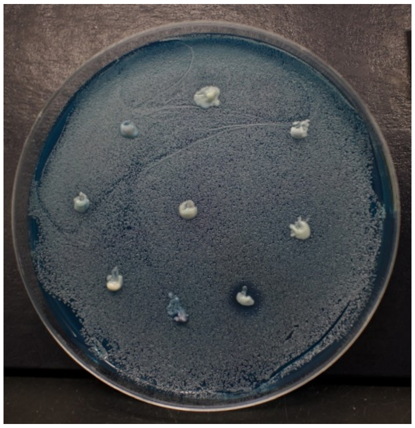

BY4741

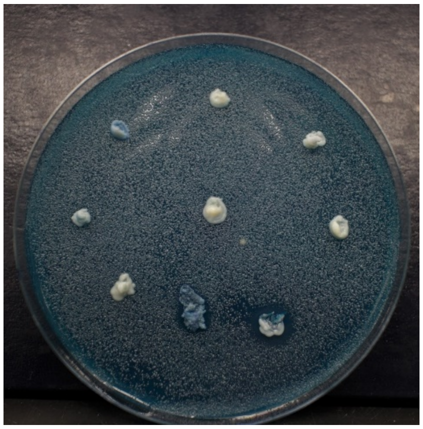

SSS 104

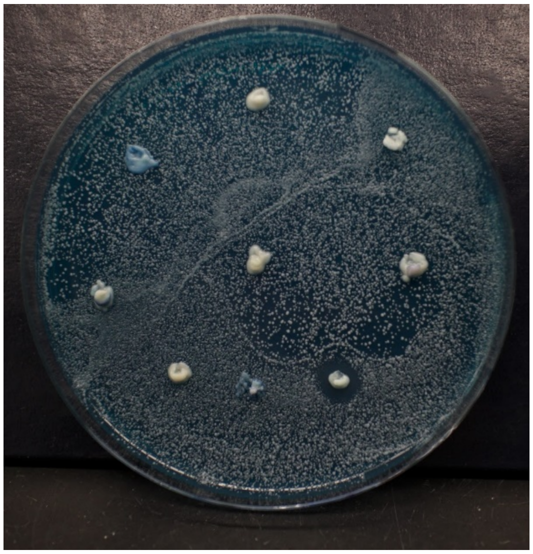

Key

<table>
<tr><td colspan="6">YB-2047</td></tr>
<tr><td colspan="3">Y-12602</td><td colspan="3">YB-4137</td></tr>
<tr><td colspan="2">Y-11842</td><td colspan="2">Y-17218</td><td colspan="2">YB-4565</td></tr>
<tr><td colspan="3">Y-6179</td><td colspan="3">Y-5688</td></tr>
<tr><td colspan="6">Y-6177</td></tr>
</table>

BY4741

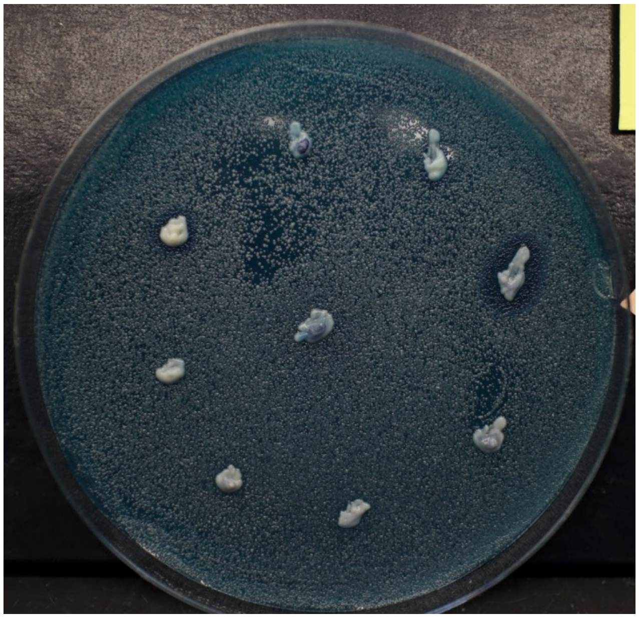

Key

<table>
<tr><td colspan="6">Y-17353</td></tr>
<tr><td colspan="3">Y-63714</td><td colspan="3">Y-63708</td></tr>
<tr><td colspan="2">Y-63713</td><td colspan="2">Y-63715</td><td colspan="2">Y-63709</td></tr>
<tr><td colspan="3">Y-63712</td><td colspan="3">Y-63710</td></tr>
<tr><td colspan="6">Y-63711</td></tr>
</table>

BY4741

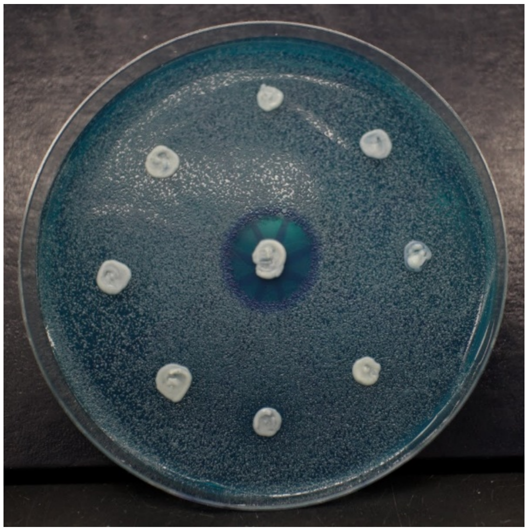

SSS 104

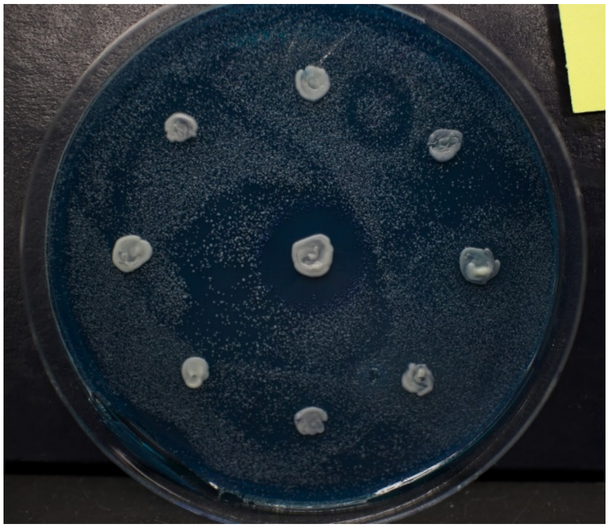

NCYC 2729

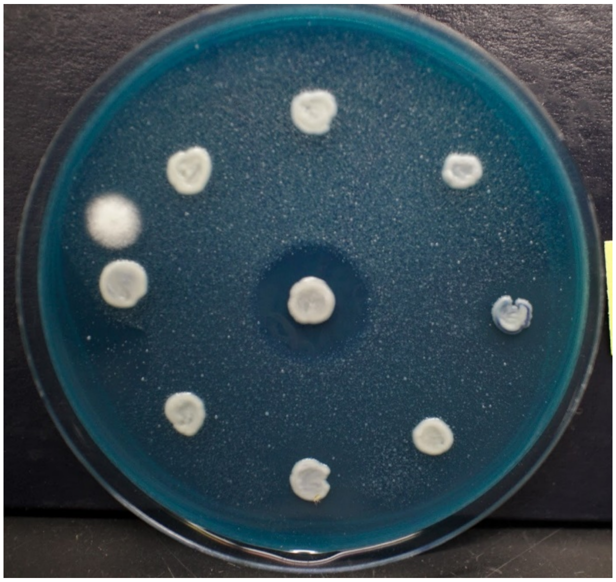

Key

<table>
<tr><td colspan="6">Y-63716</td></tr>
<tr><td colspan="3">1200</td><td colspan="3">Y-63717</td></tr>
<tr><td colspan="2">1130</td><td colspan="2">1133</td><td colspan="2">851</td></tr>
<tr><td colspan="3">1119</td><td colspan="3">859</td></tr>
<tr><td colspan="6">838</td></tr>
</table>

K12

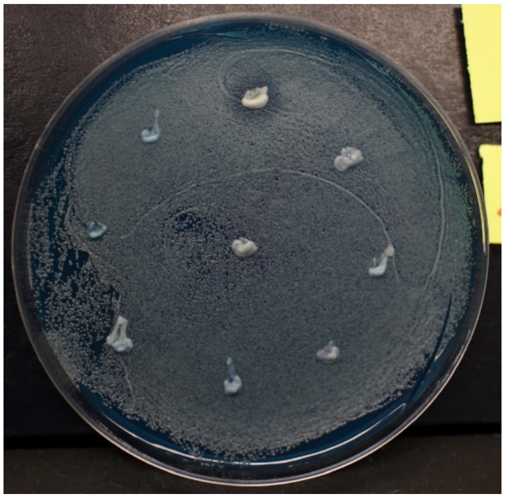

SSS 104

BY4741

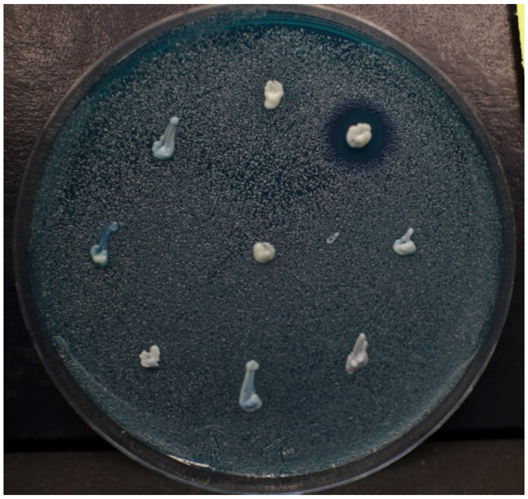

NCYC 2729

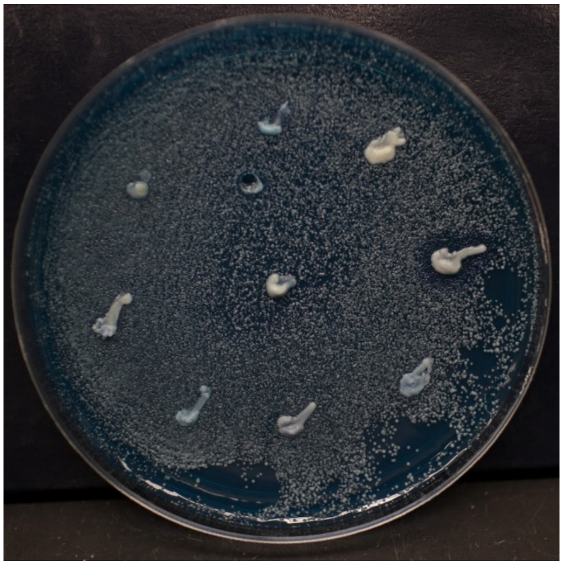

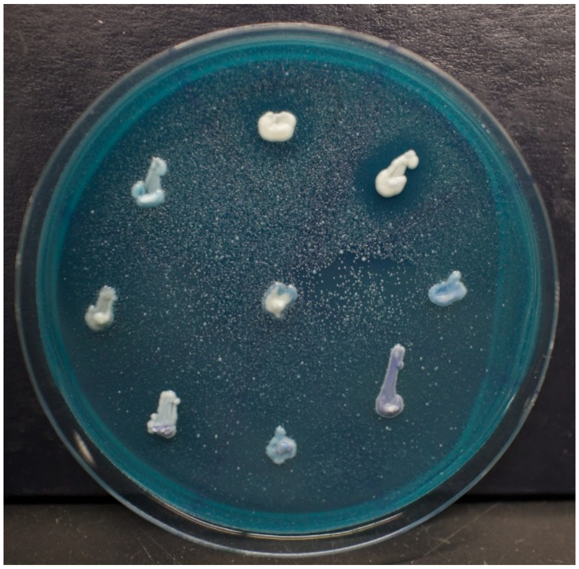

Key

<table>
<tr><td colspan="6">Y-580</td></tr>
<tr><td colspan="3">Y-12624</td><td colspan="3">Y-670</td></tr>
<tr><td colspan="2">Y-11845</td><td colspan="2">Y-12646</td><td colspan="2">Y-846</td></tr>
<tr><td colspan="3">Y-1374</td><td colspan="3">Y-969</td></tr>
<tr><td colspan="6">Y-972</td></tr>
</table>

NCYC 2729

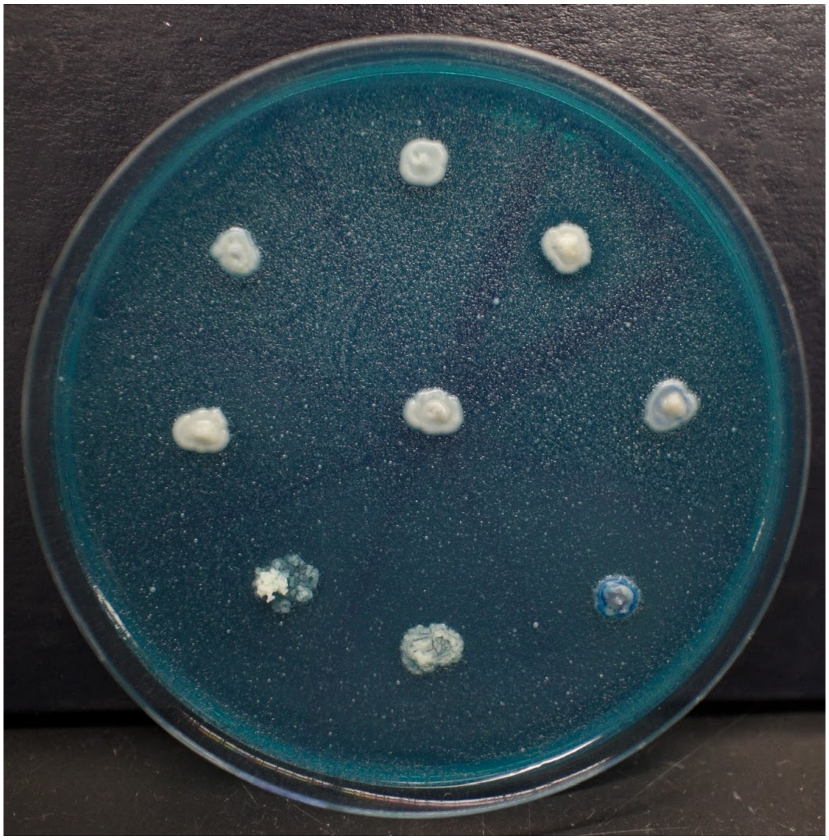

Key

| | | | | |
|---|---|---|---|---|
| Y-12648 | | | | |
| YB-254 | | | Y-17034 | |
| Y-63718 | | **YB-432** | | Y-27339 |
| Y-63707 | | | Y-27470 | |
| Y-48770 | | | | |

NCYC 2729

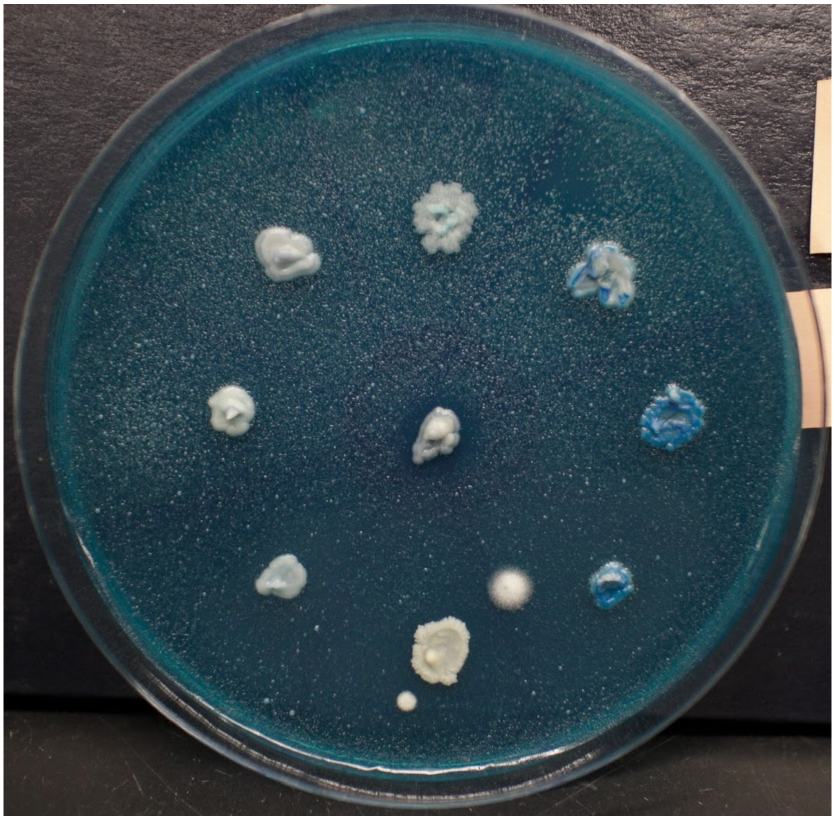

Key

| | | |
|---|---|---|
| Y-567 | | |
| Y-975 | | Y-851 |
| Y-954 | Y-976 | Y-852 |
| **YB-908** | | Y-897 |
| Y-898 | | |

K12

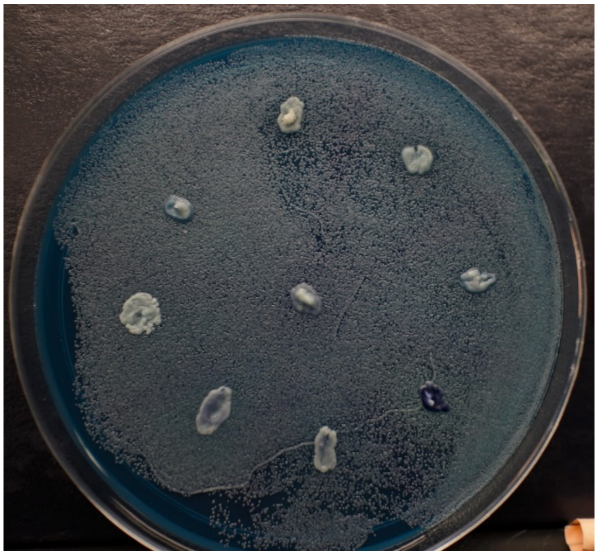

FY4

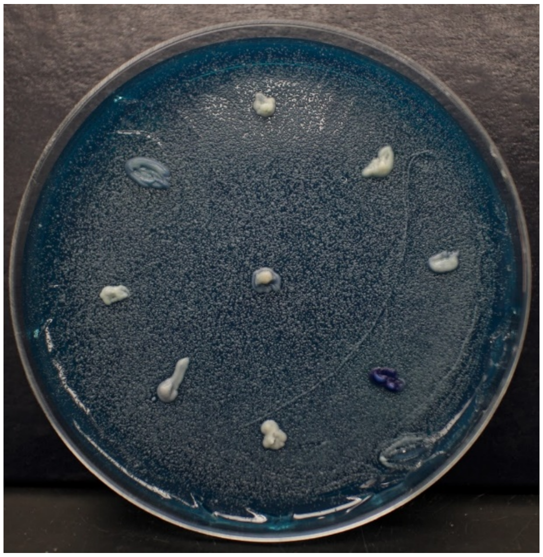

CBS 7001

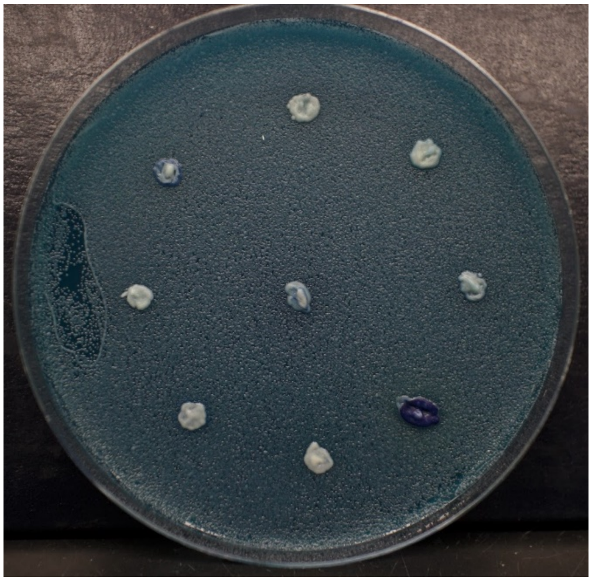

NCYC 2729

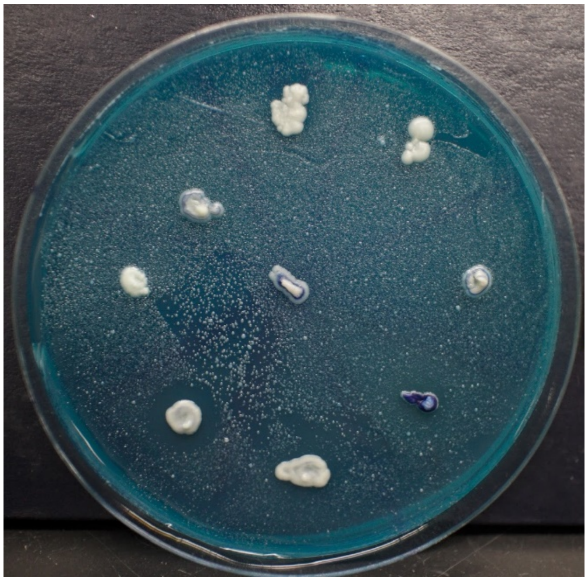

Key

<table>
<tr><td colspan="6">Y-977</td></tr>
<tr><td colspan="3">Y-1429</td><td colspan="3">Y-1018</td></tr>
<tr><td colspan="2">Y-1428</td><td colspan="2">Y-1430</td><td colspan="2">Y-1089</td></tr>
<tr><td colspan="3">Y-1370</td><td colspan="3">Y-1285</td></tr>
<tr><td colspan="6">Y-1301</td></tr>
</table>

NCYC 2729

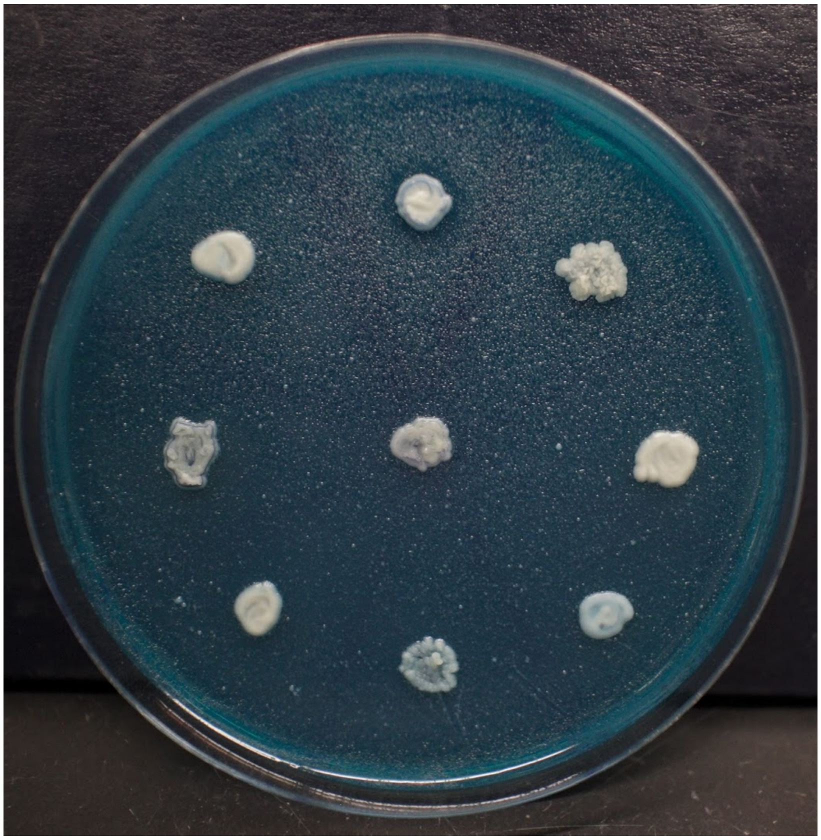
