## Supplementary material for "The Species-specific Acquisition and Diversification of a Novel Family of Killer Toxins in Budding Yeasts of the Saccharomycotina": File S2

**File S2. Image data of killer phenotypes exhibited by strains of *Saccharomyces* yeasts as summarized in figure 1.** The organization of the 21 strains of *Saccharomyces* yeasts arrayed on killer assay agar plates. Each killer yeast was assayed for toxin activity against 45 different lawns of susceptible yeast strains. n/a: not applicable.

**PLATE KEY (Pages 2-13)**

| | 1 | 2 | 3 | 4 | 5 | 6 | 7 | 8 |
|---|---|---|---|---|---|---|---|---|
| A | n/a | - | BJH001 | - | NCYC 190 | - | n/a | - |
| B | - | Y-2429 | - | 1116 | - | NCYC 1001 | - | MS300C |
| C | CYC 1058 | - | CYC 1113 | - | n/a | - | n/a | - |
| D | - | Y8.5 | - | Y-63717 | - | Y-63716 | - | Y-63711 |
| E | Y-12602 | - | YB-4565 | - | Y-1088 | - | n/a | - |
| F | - | n/a | - | - | - | - | - | - |

Lawn: BJH001

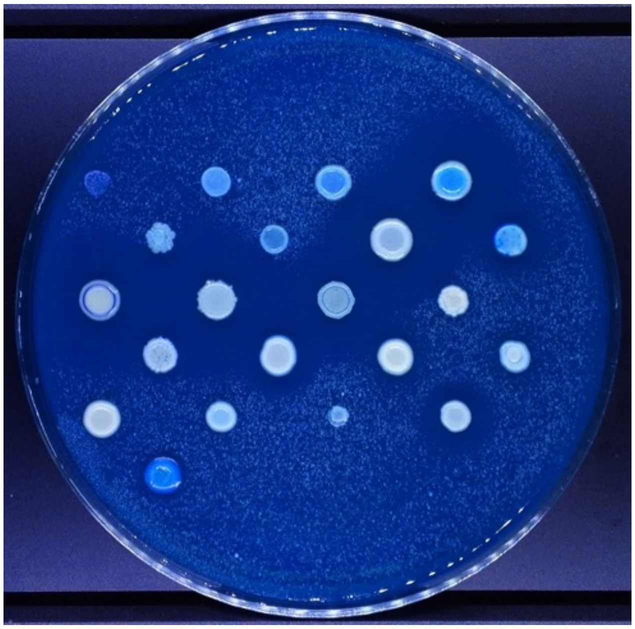

Lawn: NCYC 738

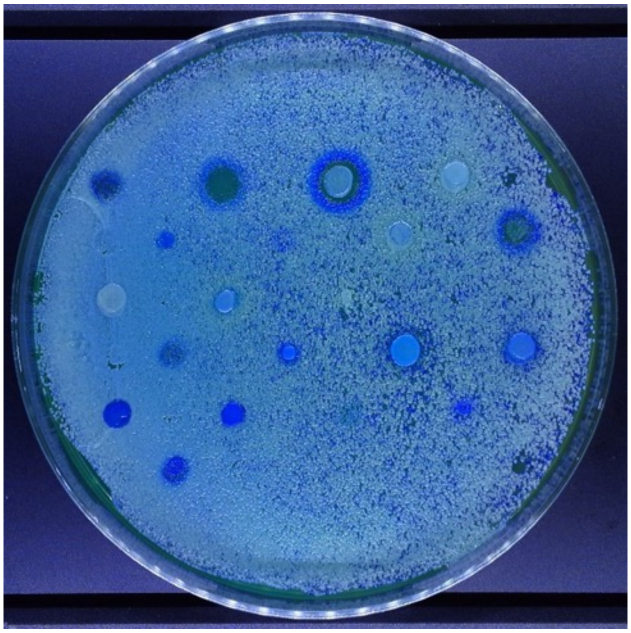

Lawn: DSM70459

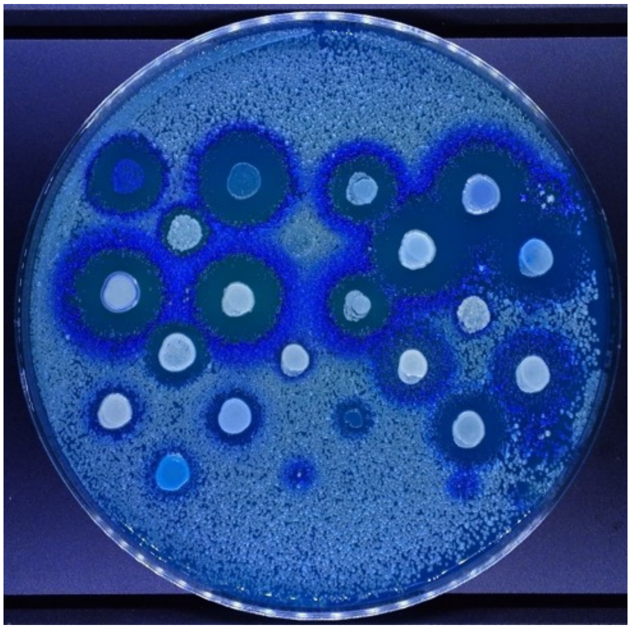

Lawn: DBVPG 1373

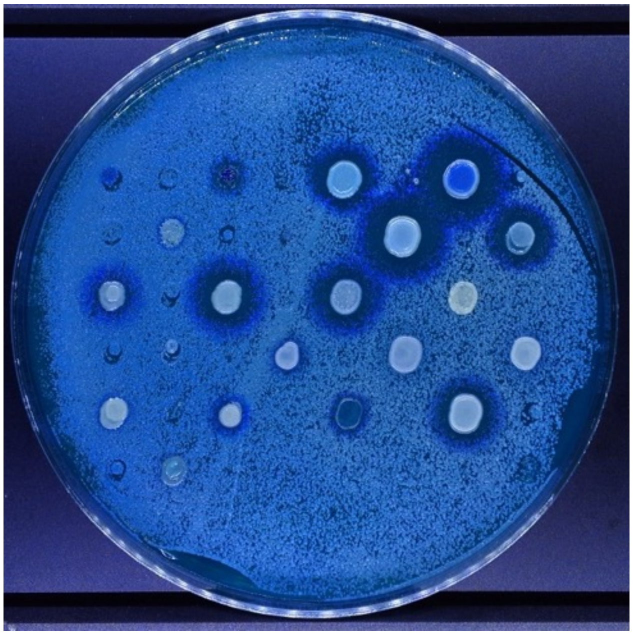

Lawn: Y-27788

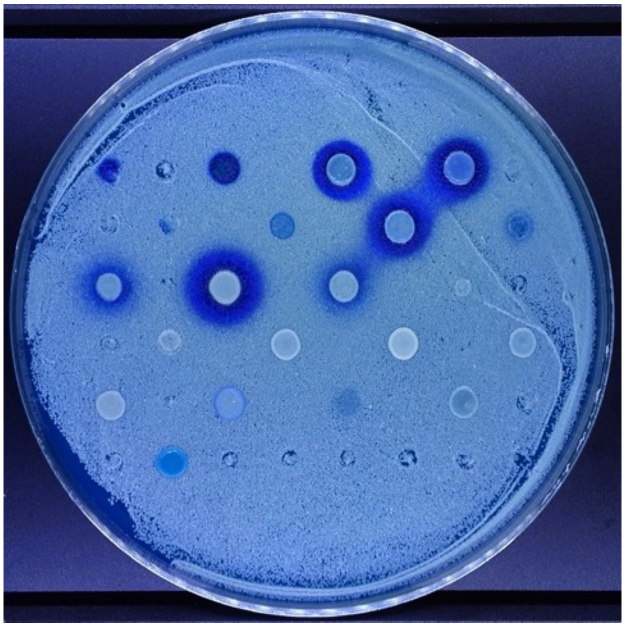

Lawn: Y-27106

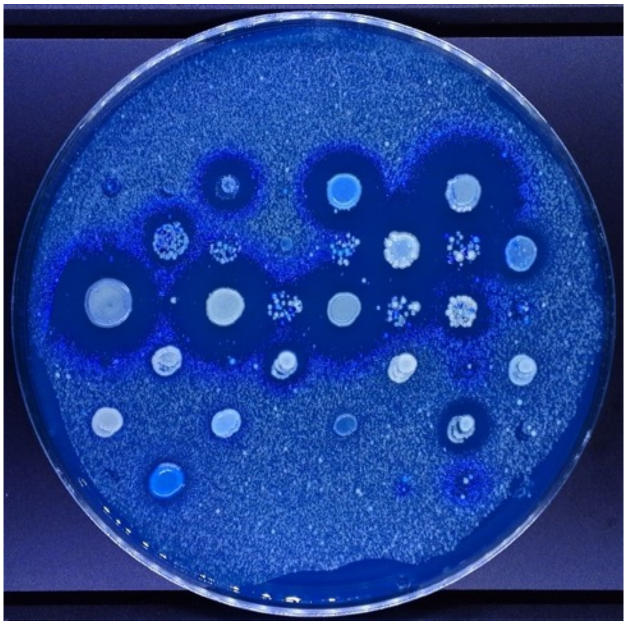

Lawn: Y-5509

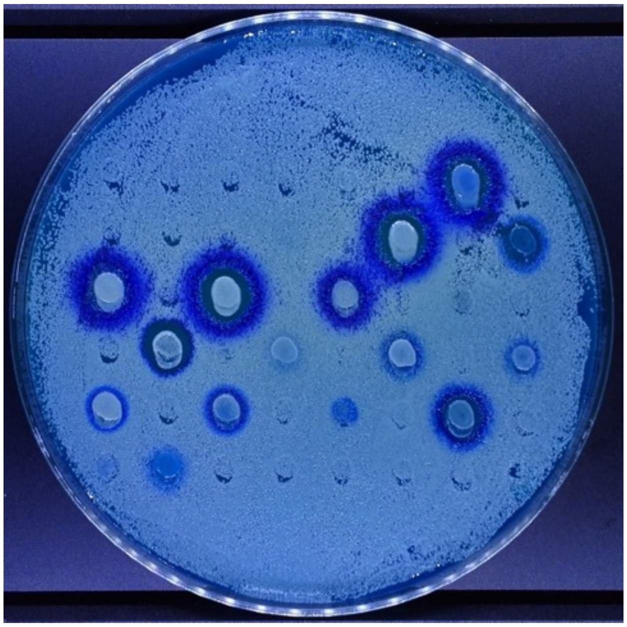

Lawn: Y-1891

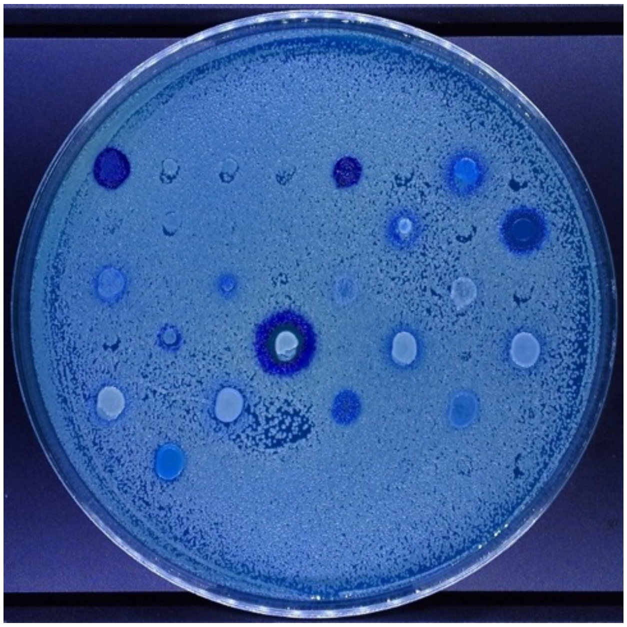

Lawn: YB 432

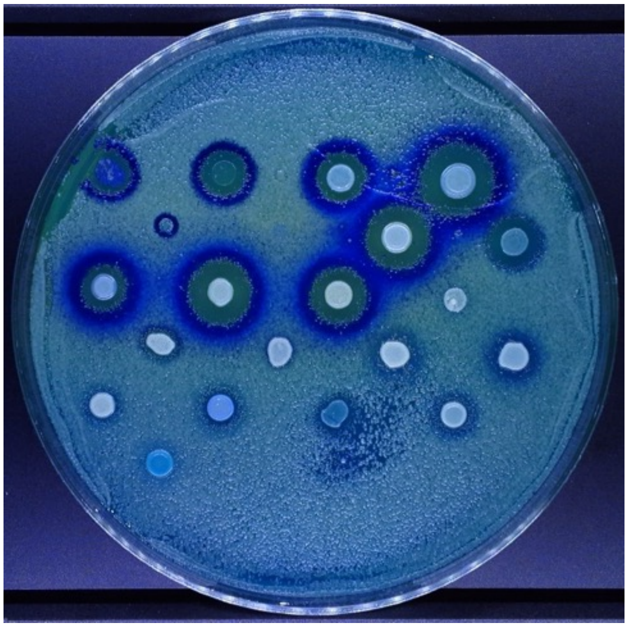

Lawn: NCYC 1001

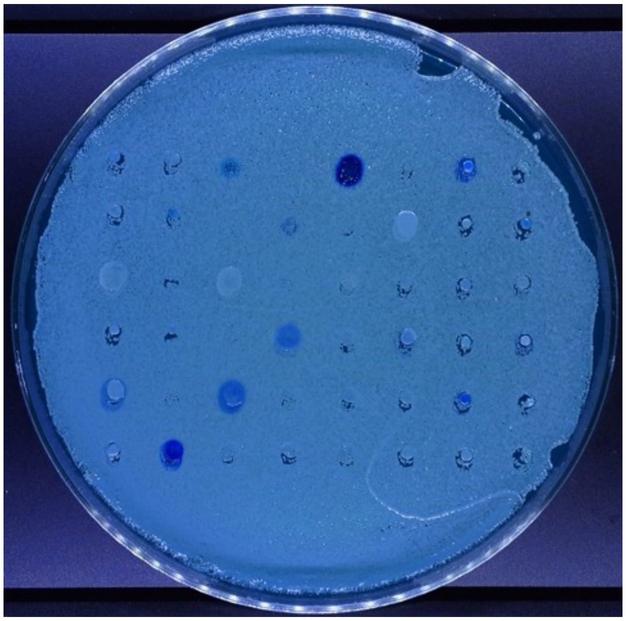

Lawn: NCYC 1006

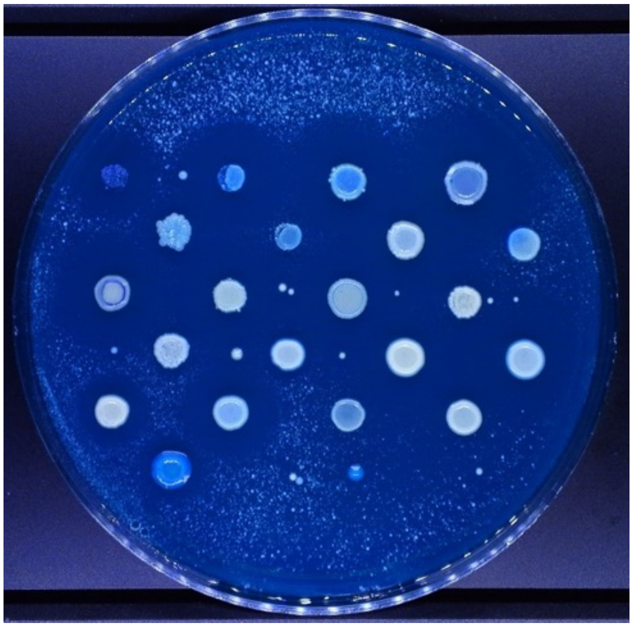

Lawn: Ms300C

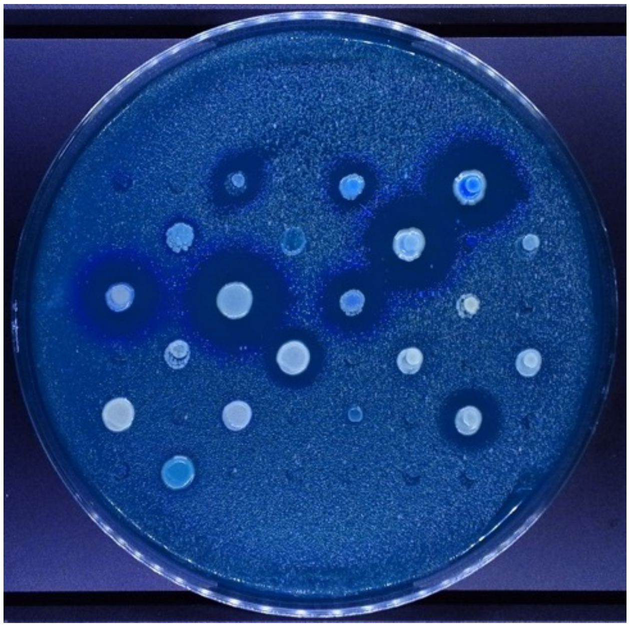

Lawn: CYC 1058

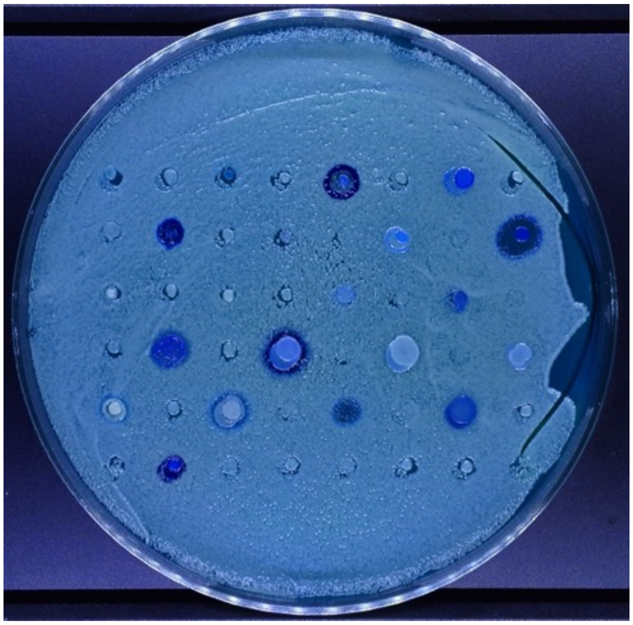

Lawn: CYC 1113

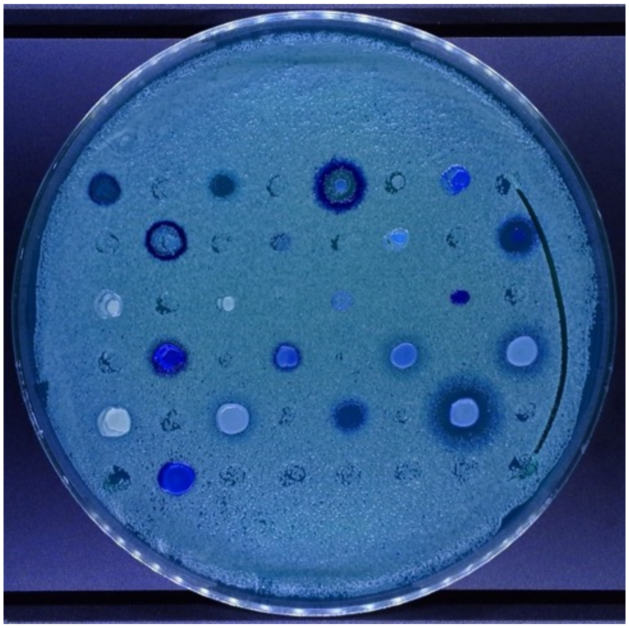

Lawn: CYC 1170

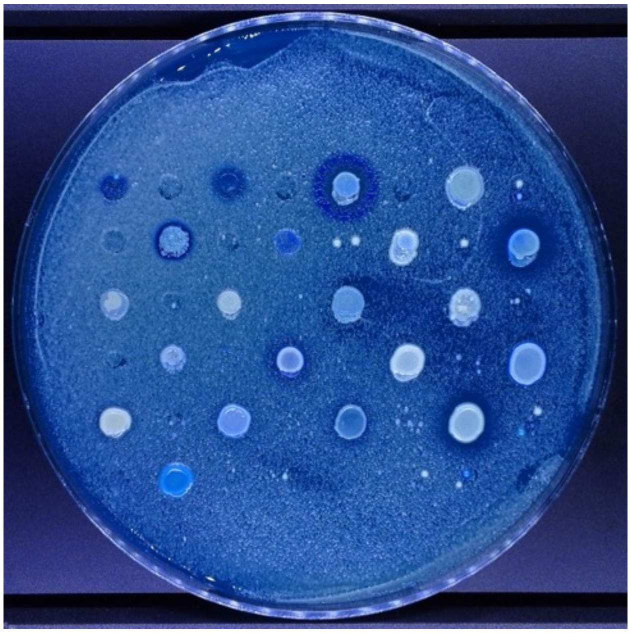

Lawn: CYC 1172

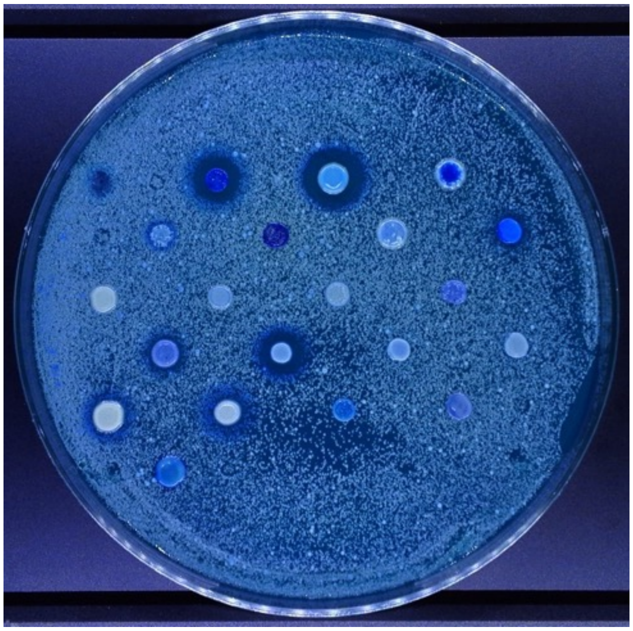

Lawn: Y-1088

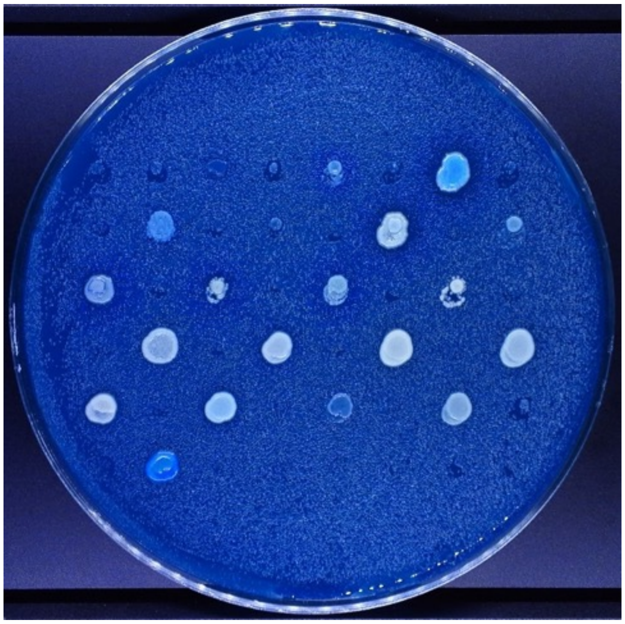

Lawn: Y-27342

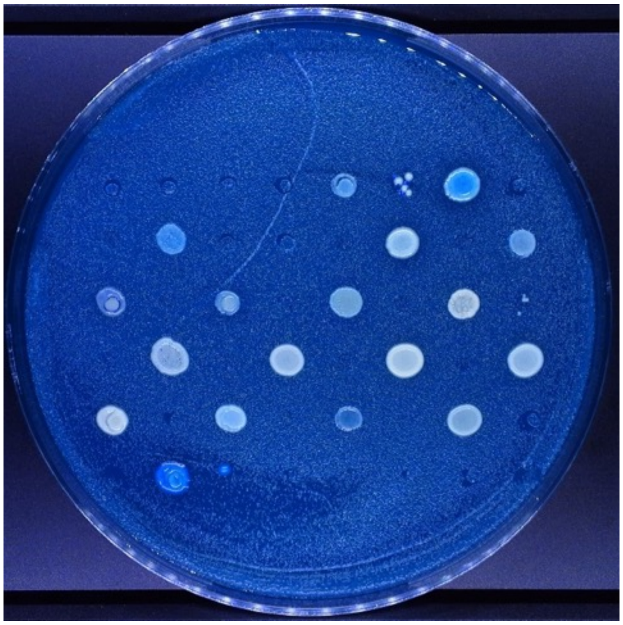

Lawn: Y-2046

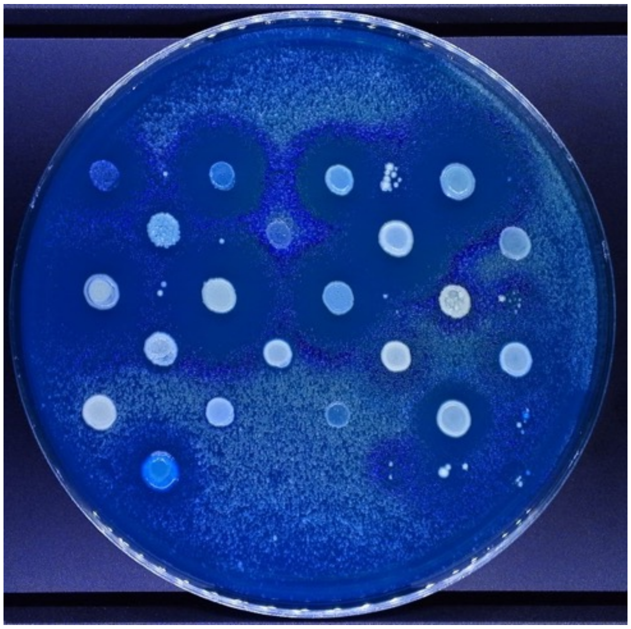

Lawn: Y-1344

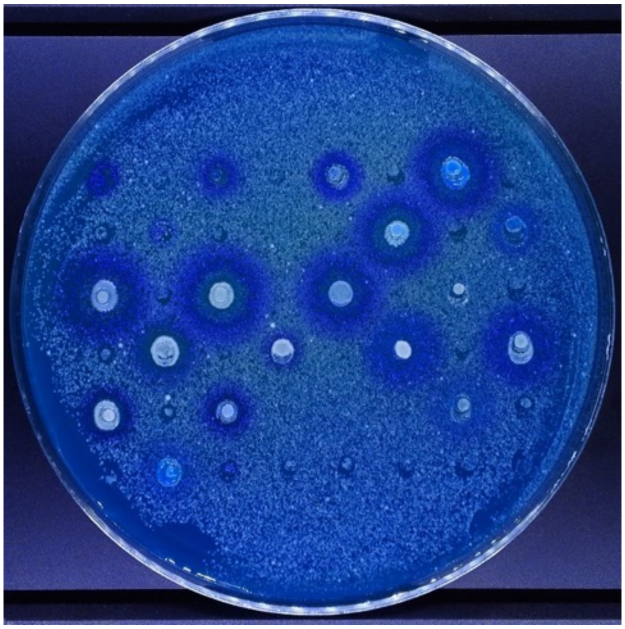

Lawn: NCYC 777

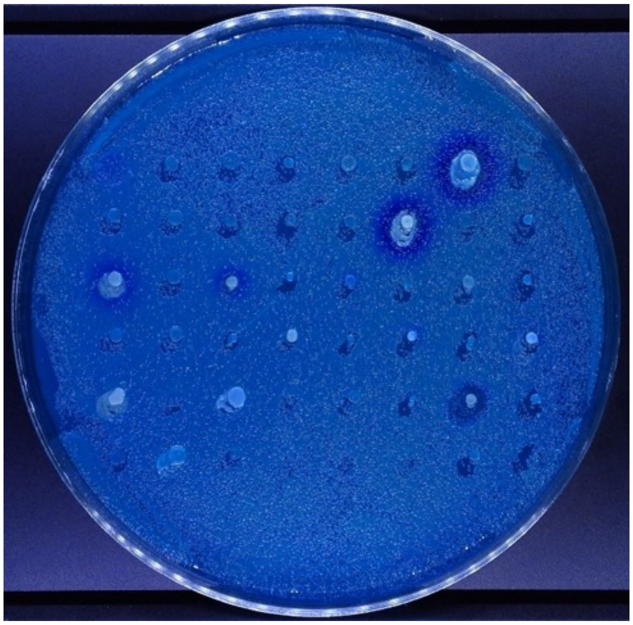

Lawn: NCYC 2898

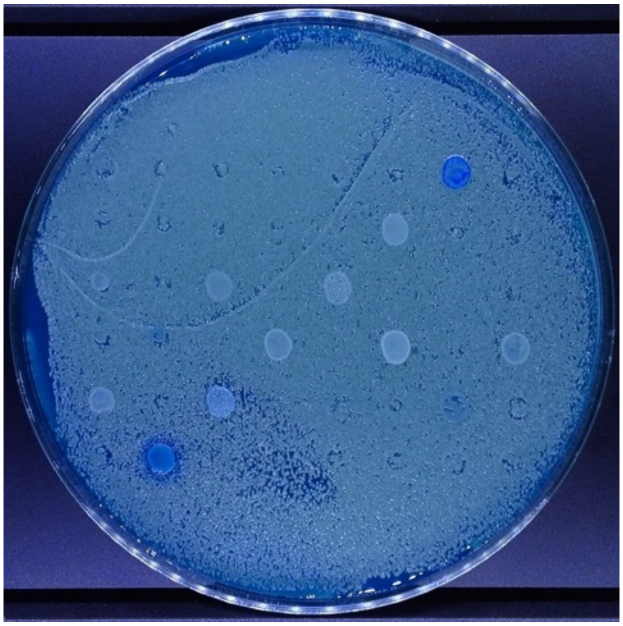

Lawn: 2729

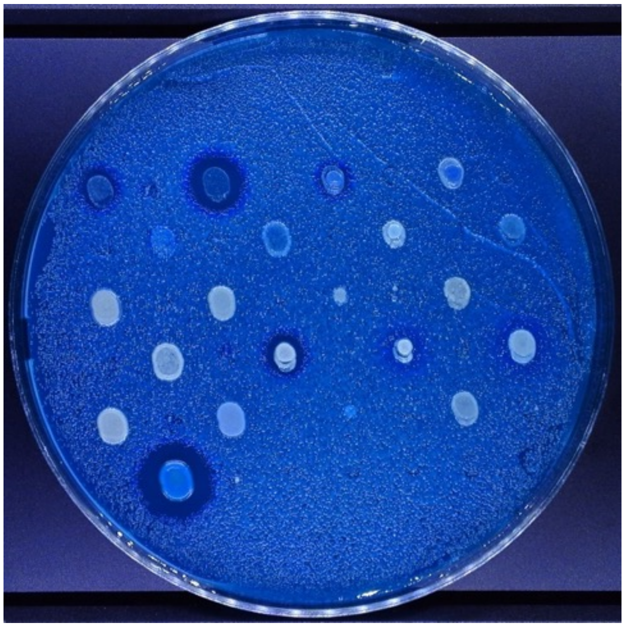

Lawn: FY4

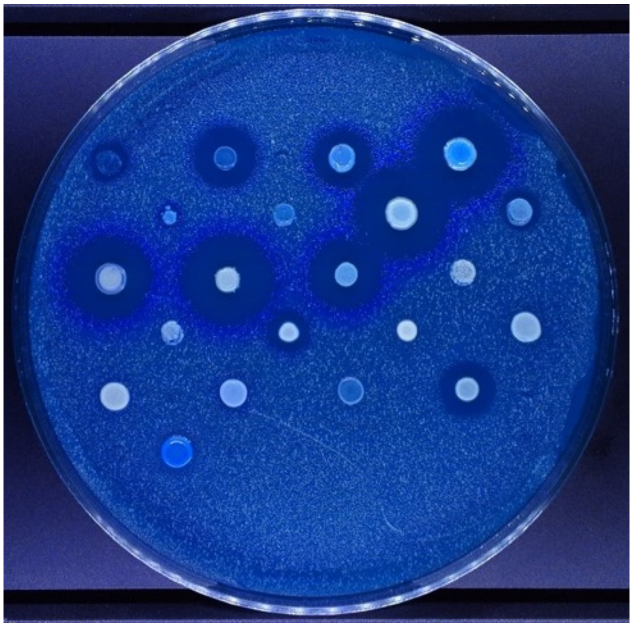

Lawn: K12

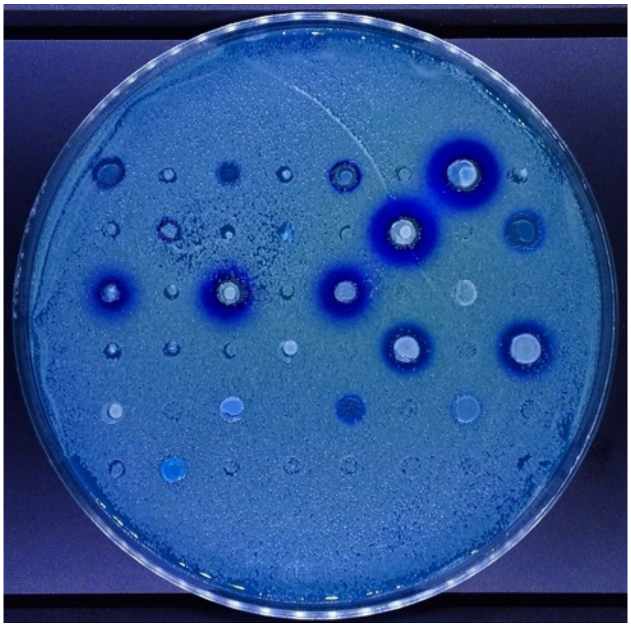

Lawn: BY4741

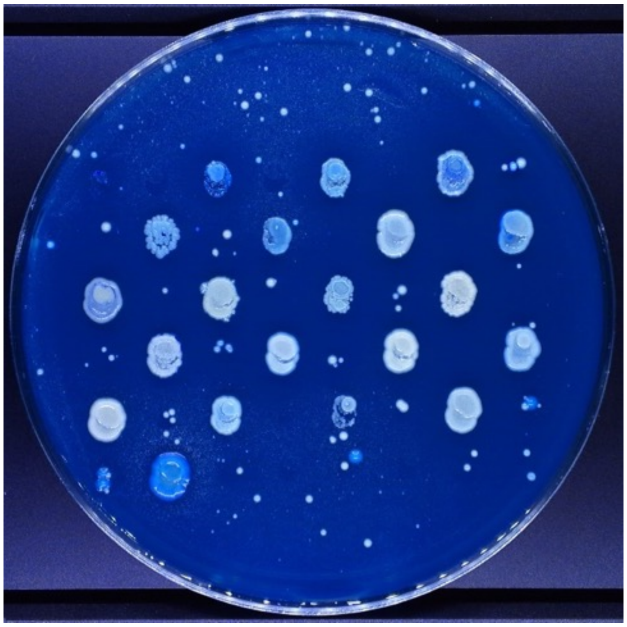

Lawn: DBVPG 6765

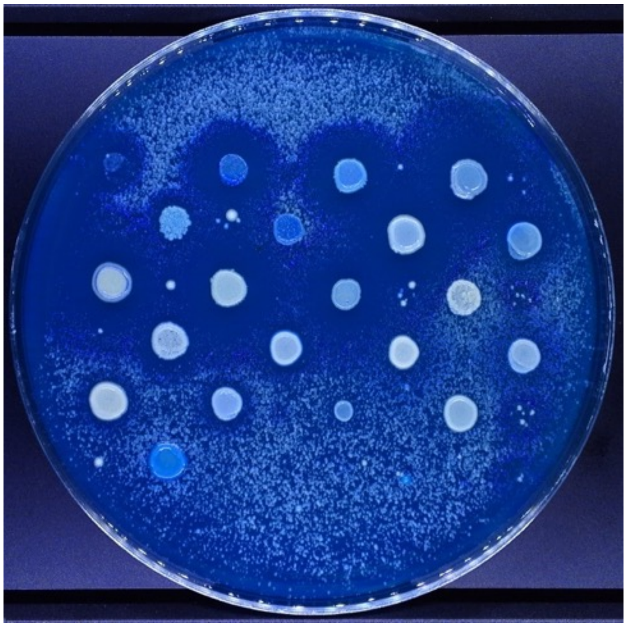

Lawn: CYC 1102

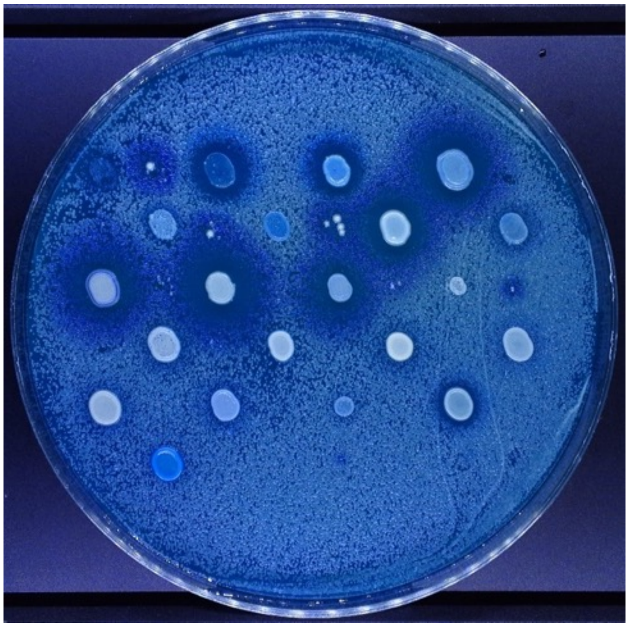

Lawn: YB-4237

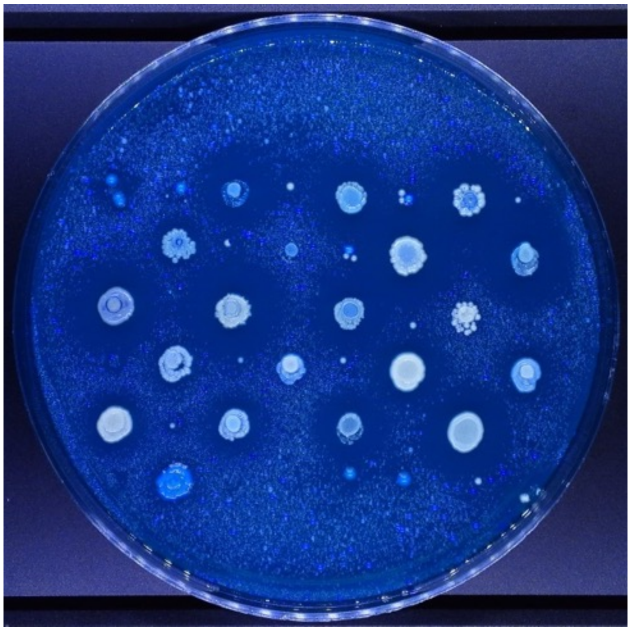

Lawn: CBS 432

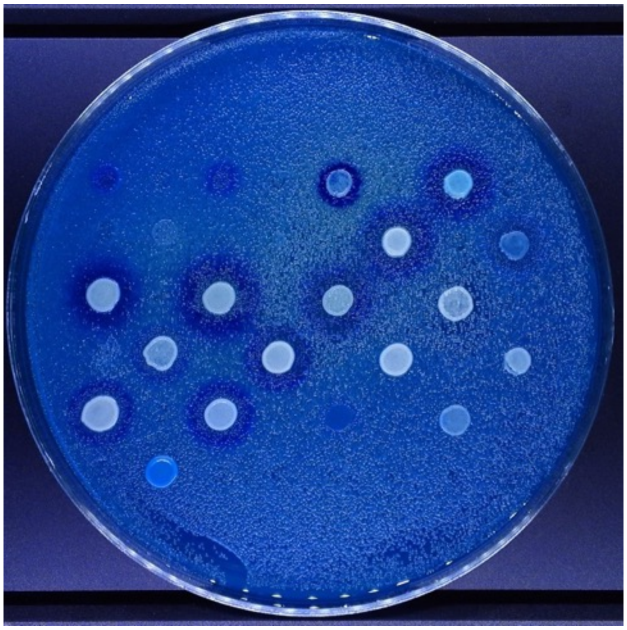

Lawn: A12C

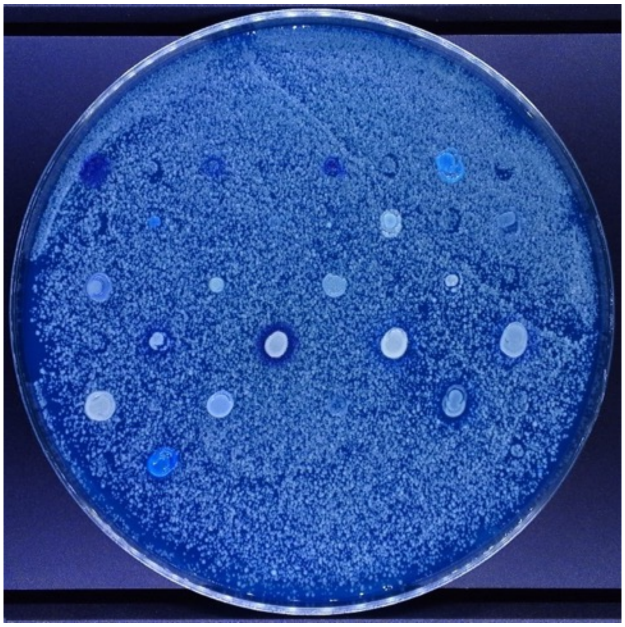

Lawn: NBRC 1815

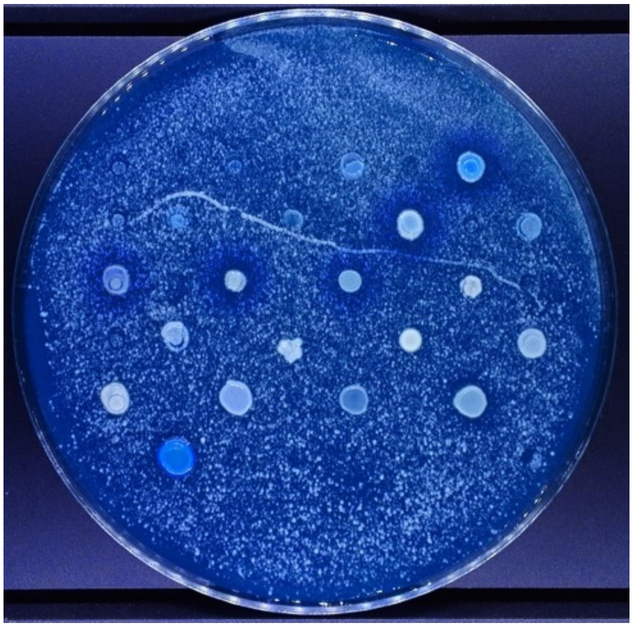

Lawn: SSS 211

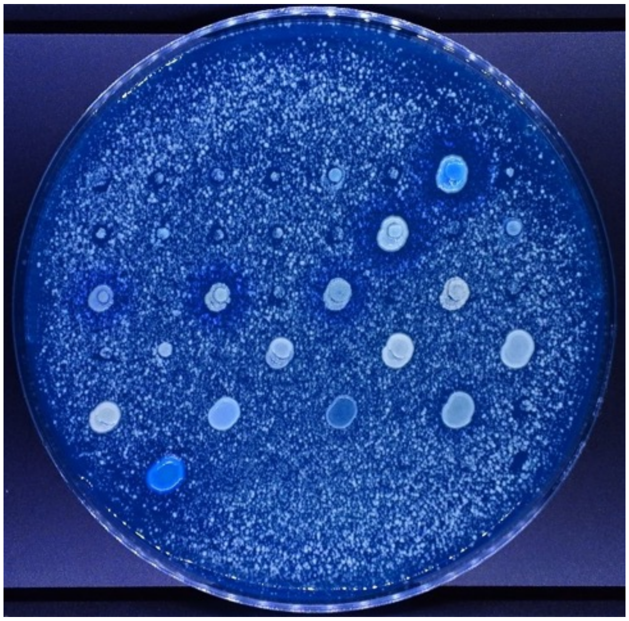

Lawn: NBRC 1802

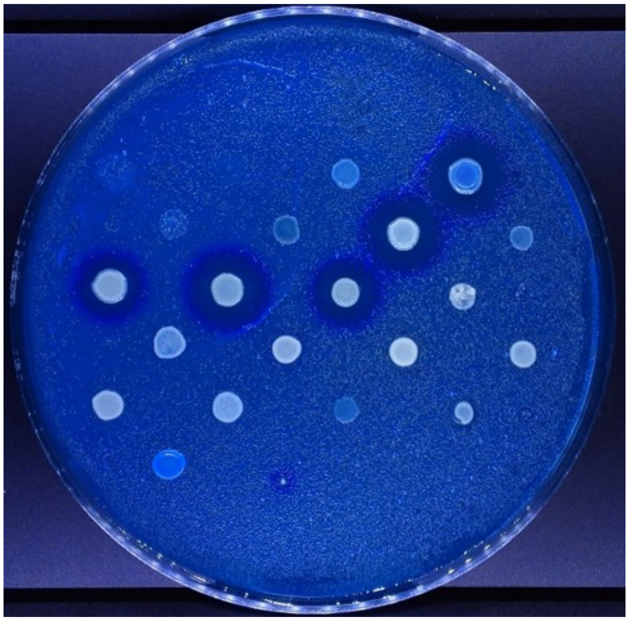

Lawn: CBS 7001

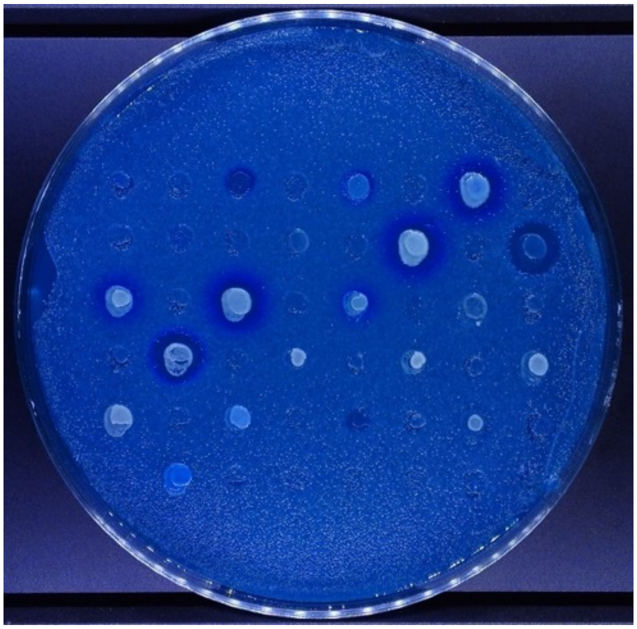

Lawn: SSS 104

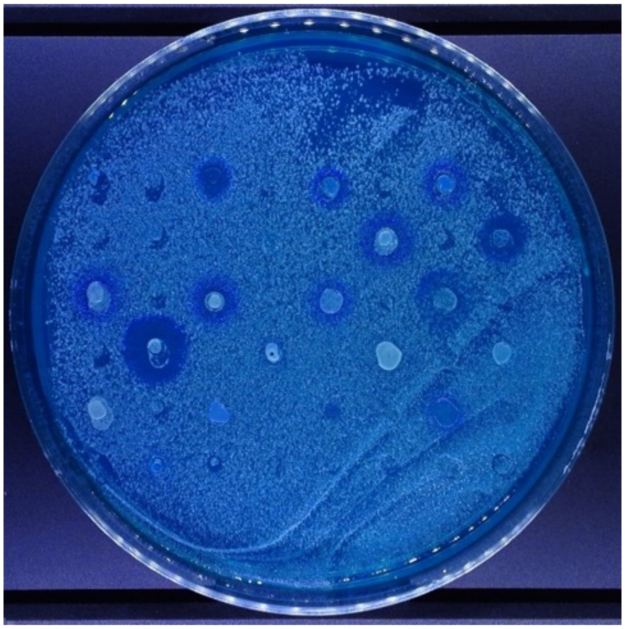

Lawn: Y-63711

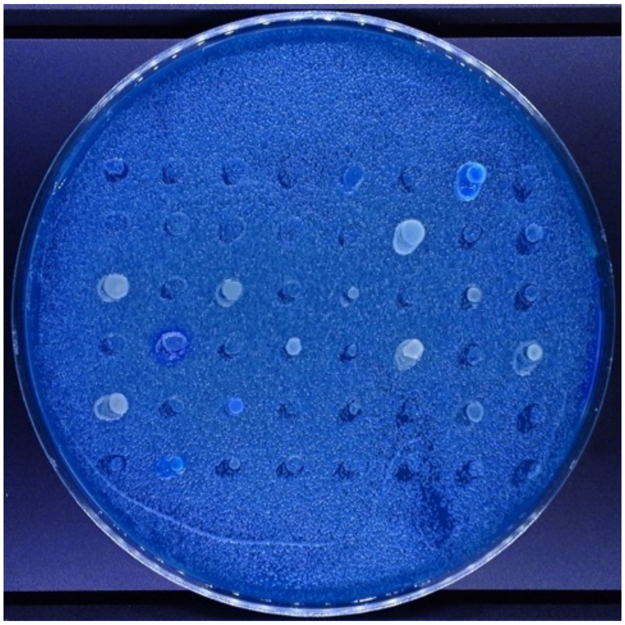

Lawn: Y-63716

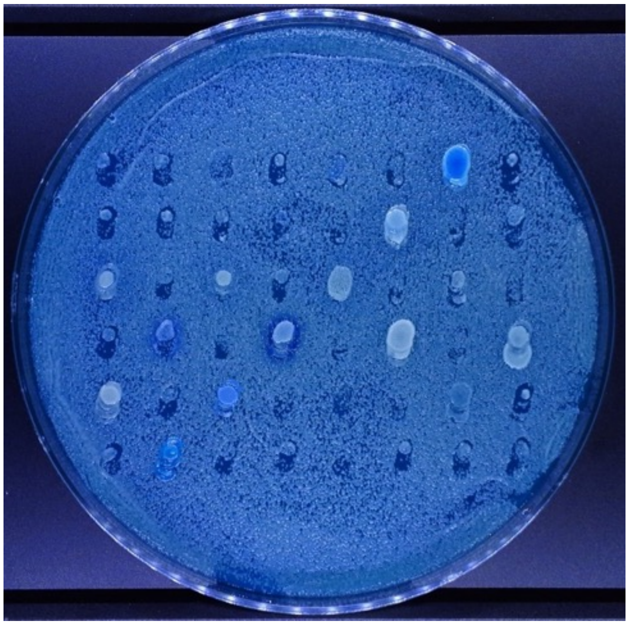

Lawn: YB-4565

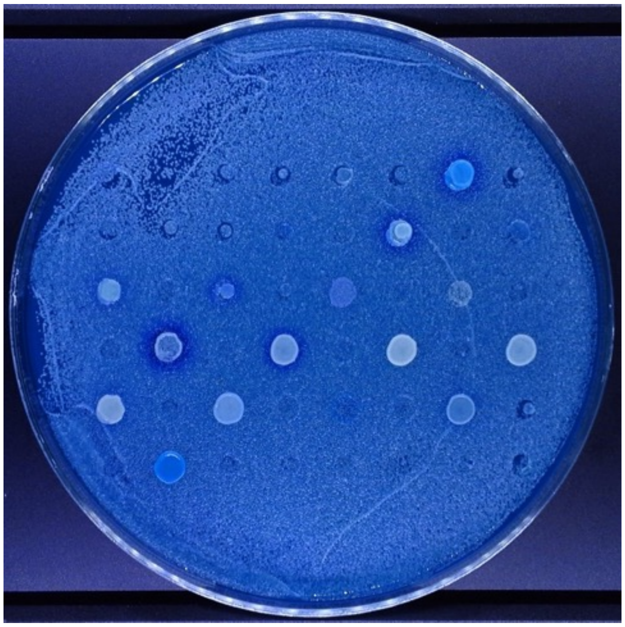

Lawn: 1116

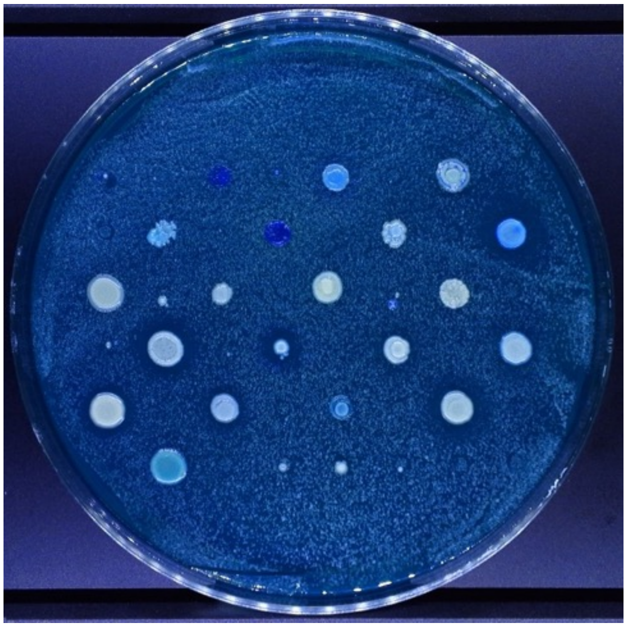

Lawn: Y-2429

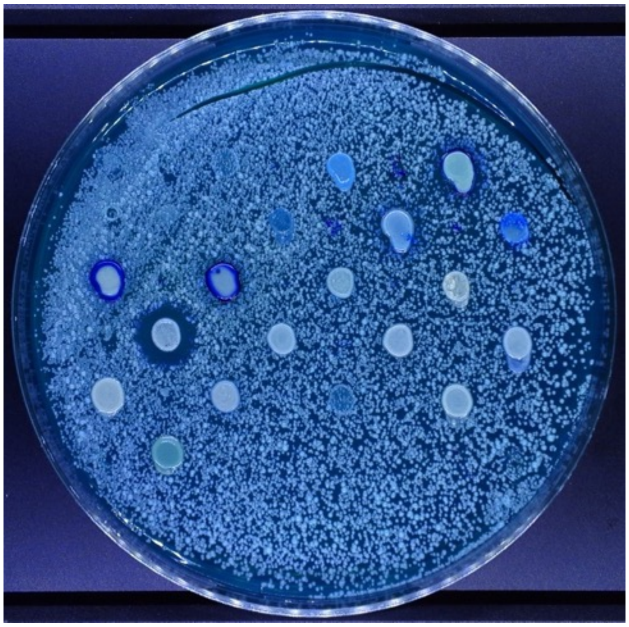

Lawn: DBVPG 6304

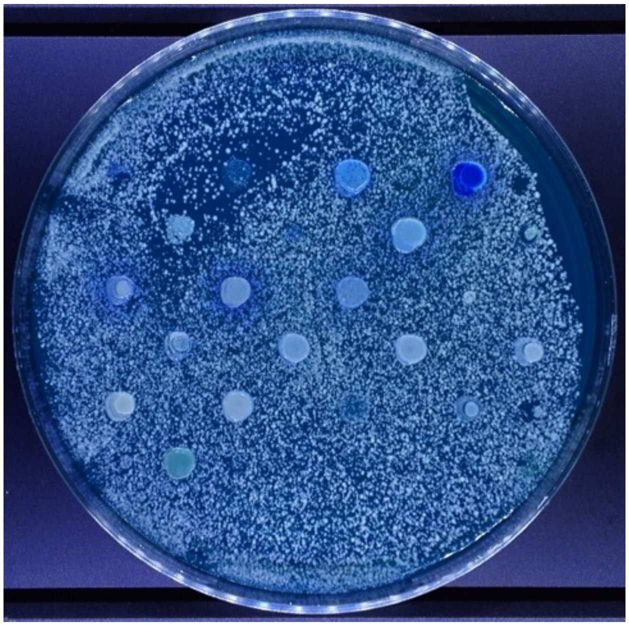

Lawn: Y-63717

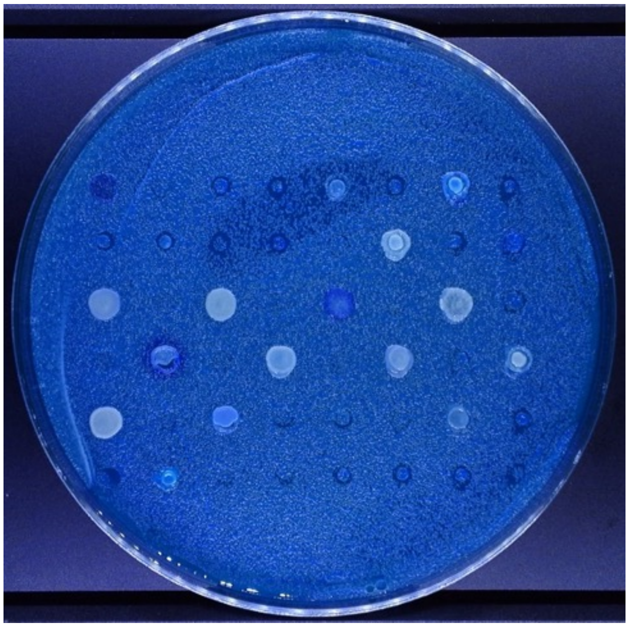

Lawn: Y8.5

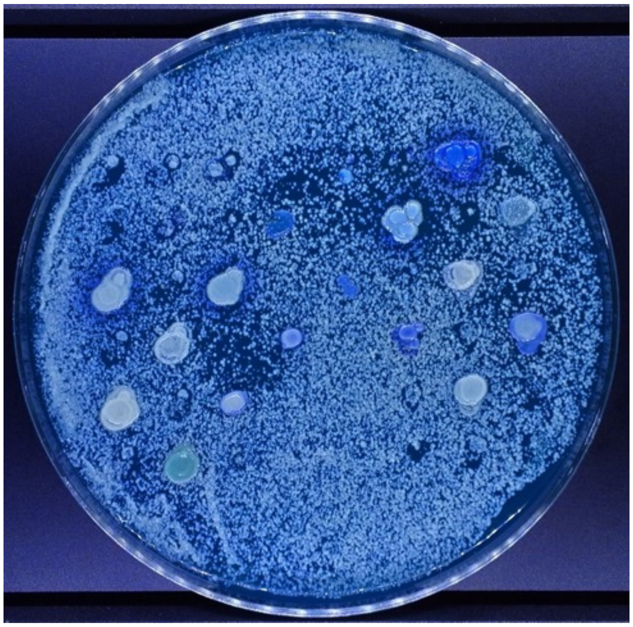

Lawn: Y-12602

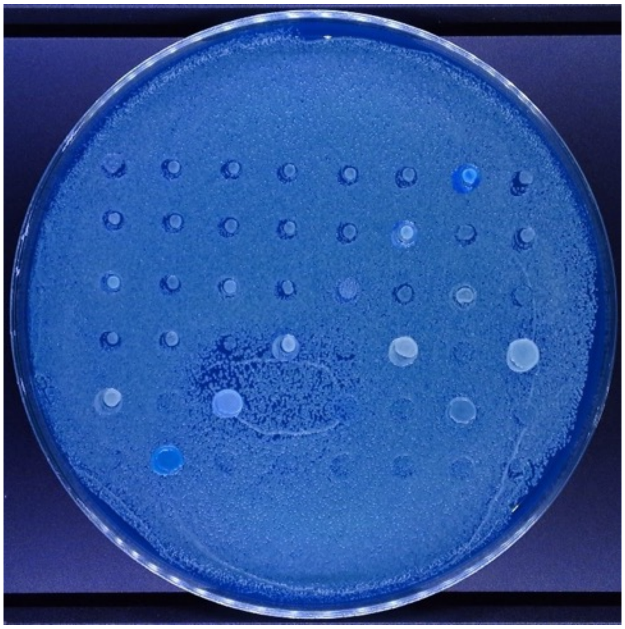

PLATE KEY (Pages 15-37)

| | 1 | 2 | 3 | 4 | 5 | 6 | 7 | 8 | 9 | 10 | 11 | 12 |
|---|---|---|---|---|---|---|---|---|---|---|---|---|
| A | - | - | - | - | - | - | - | - | - | - | - | - |
| B | - | n/a | n/a | n/a | n/a | n/a | n/a | n/a | n/a | n/a | Y-27788 | - |
| C | - | Y-27106 | n/a | Y-1891 | YB-432 | n/a | n/a | n/a | n/a | n/a | n/a | - |
| D | - | n/a | n/a | n/a | n/a | n/a | n/a | n/a | n/a | n/a | n/a | - |
| E | - | n/a | n/a | n/a | n/a | n/a | Y-2046 | Y-1344 | n/a | n/a | n/a | - |
| F | - | n/a | n/a | n/a | n/a | n/a | n/a | n/a | n/a | n/a | n/a | - |
| G | - | n/a | n/a | n/a | n/a | n/a | n/a | n/a | n/a | - | - | - |
| H | - | - | - | - | - | - | - | - | - | - | - | - |

Lawn: BJH001

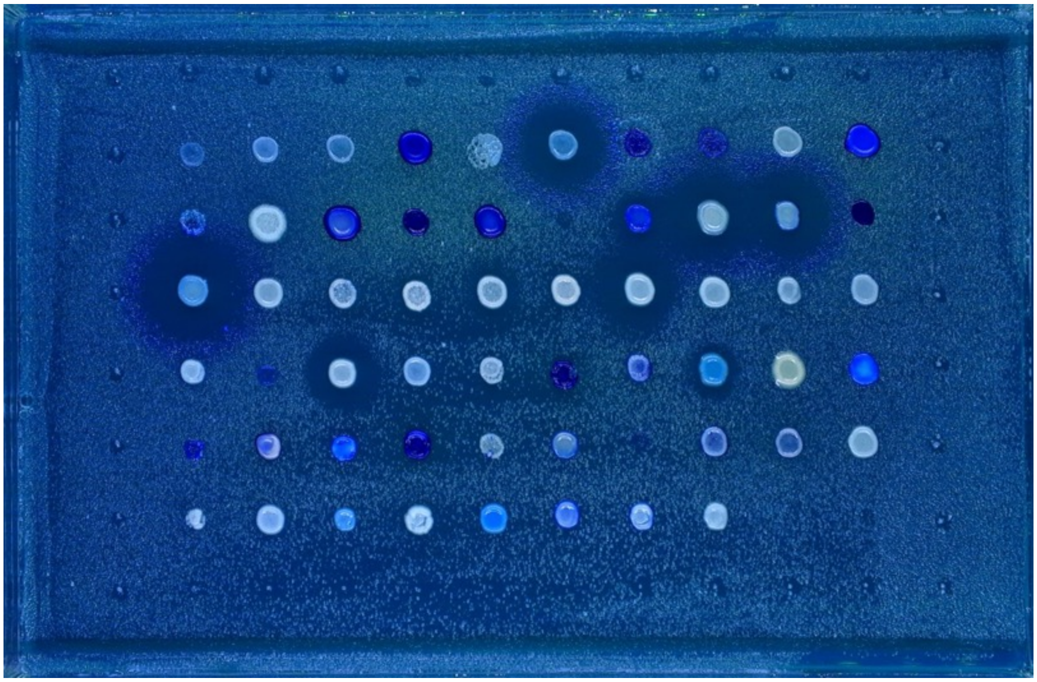

Lawn: NCYC 738

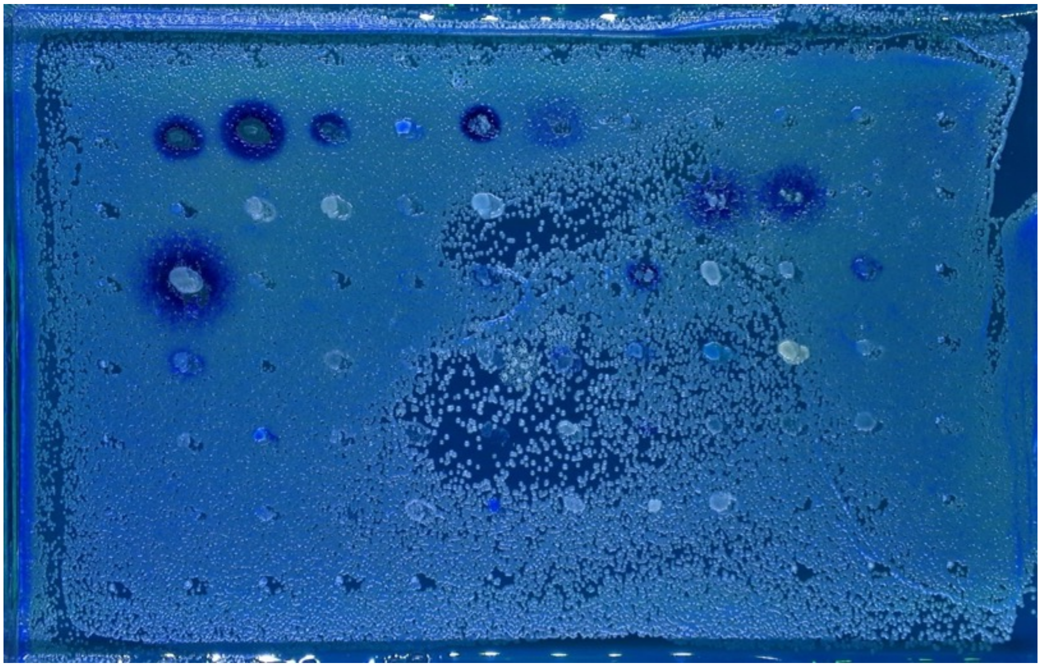

Lawn: DSM 70459

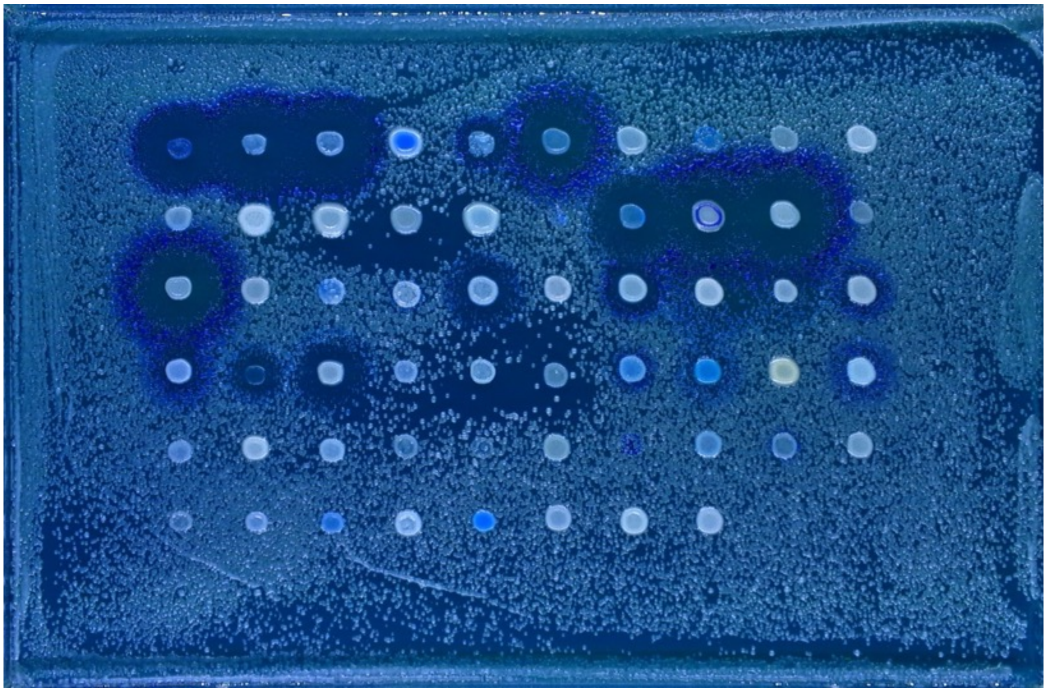

Lawn: DBVPG 1373

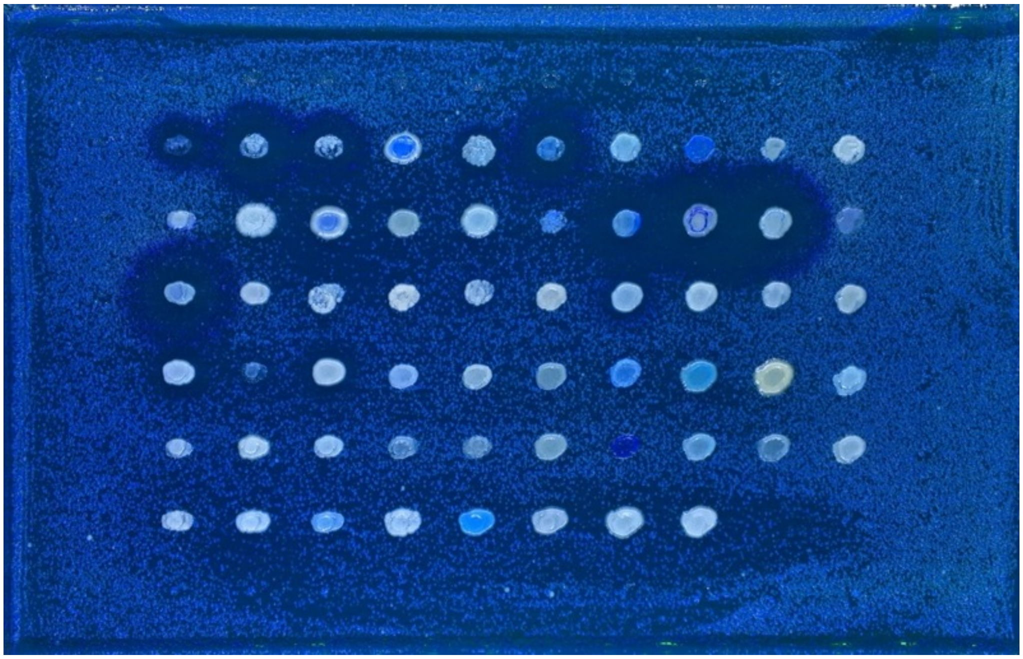

Lawn: Y-27788

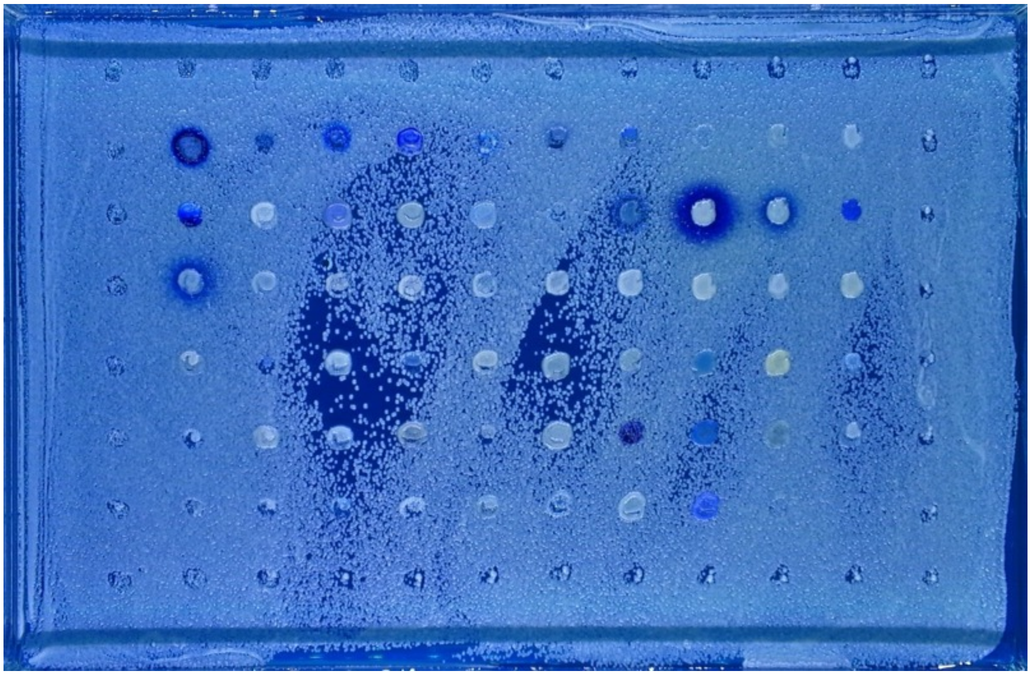

Lawn: Y-27106

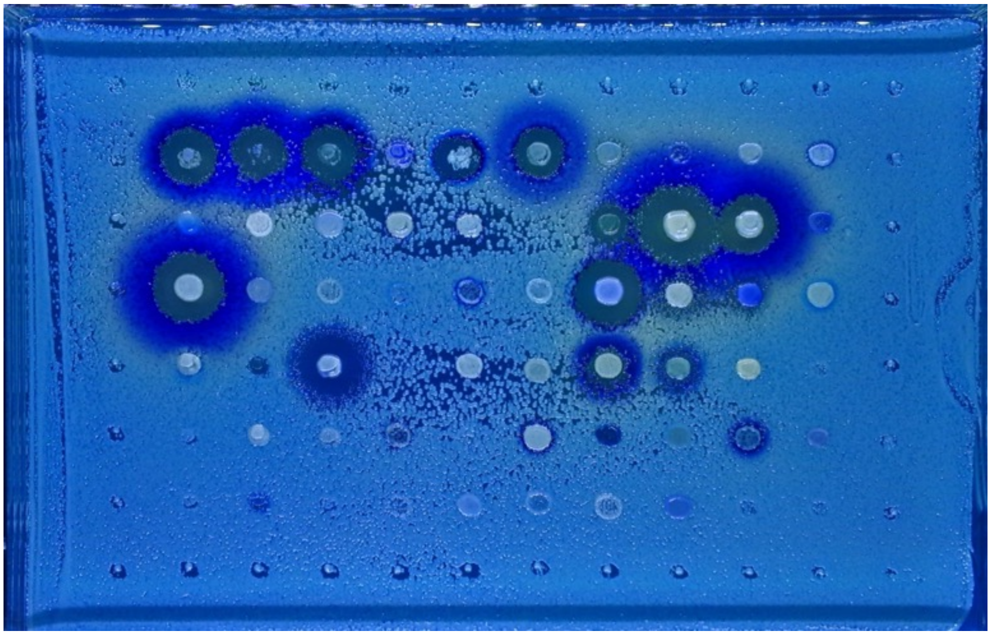

Lawn: Y-5509

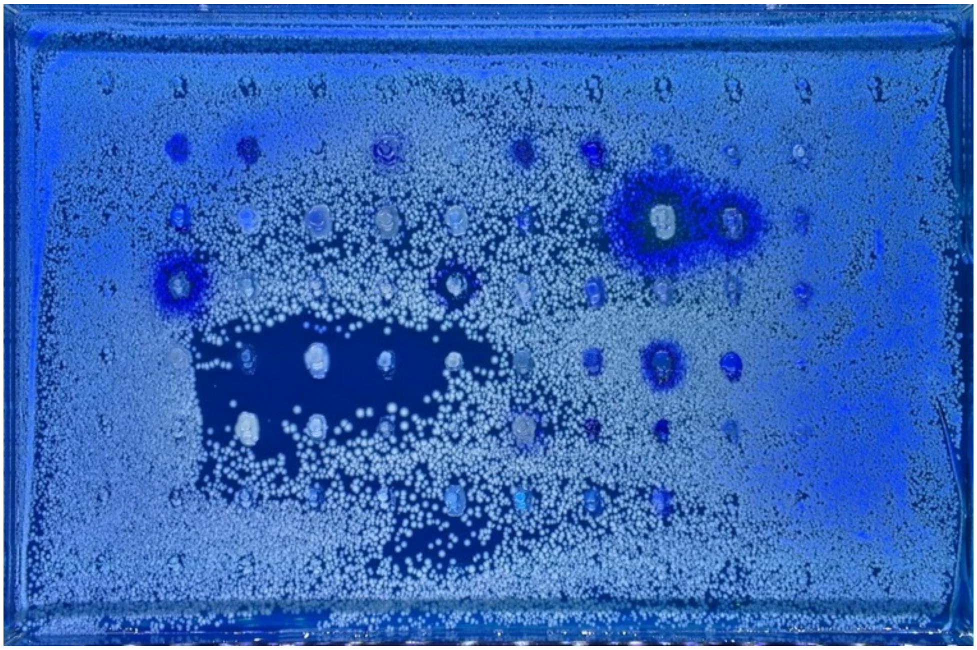

Lawn: Y-1891

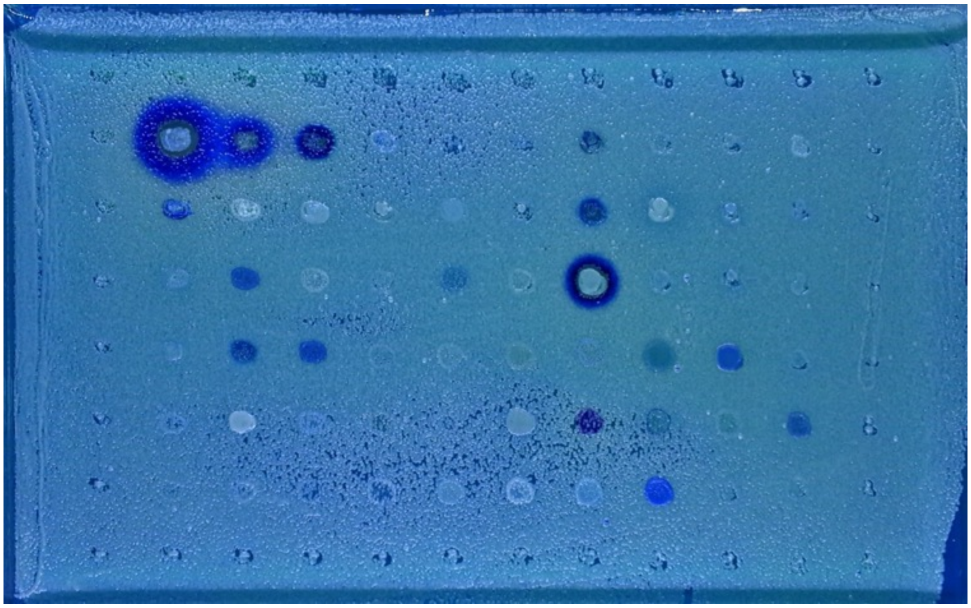

Lawn: YB-432

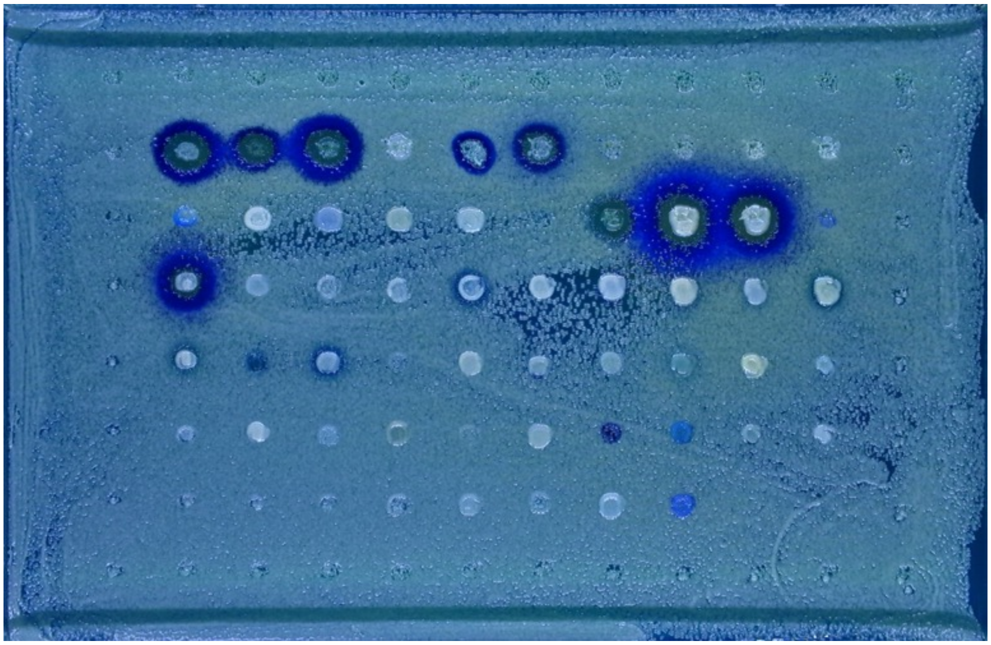

Lawn: NCYC 1001

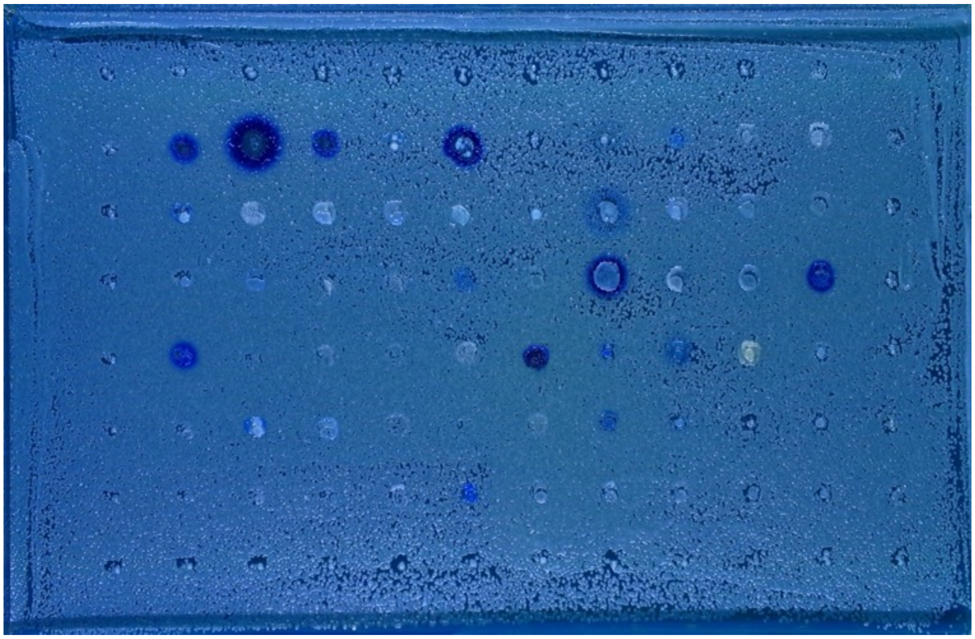

Lawn: NCYC 1006

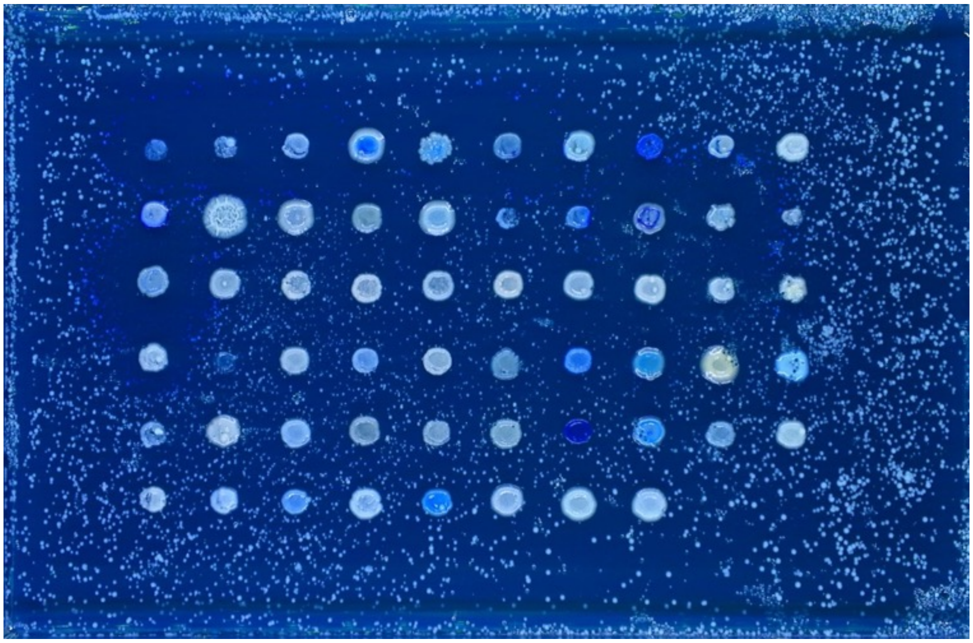

Lawn: MS 300c

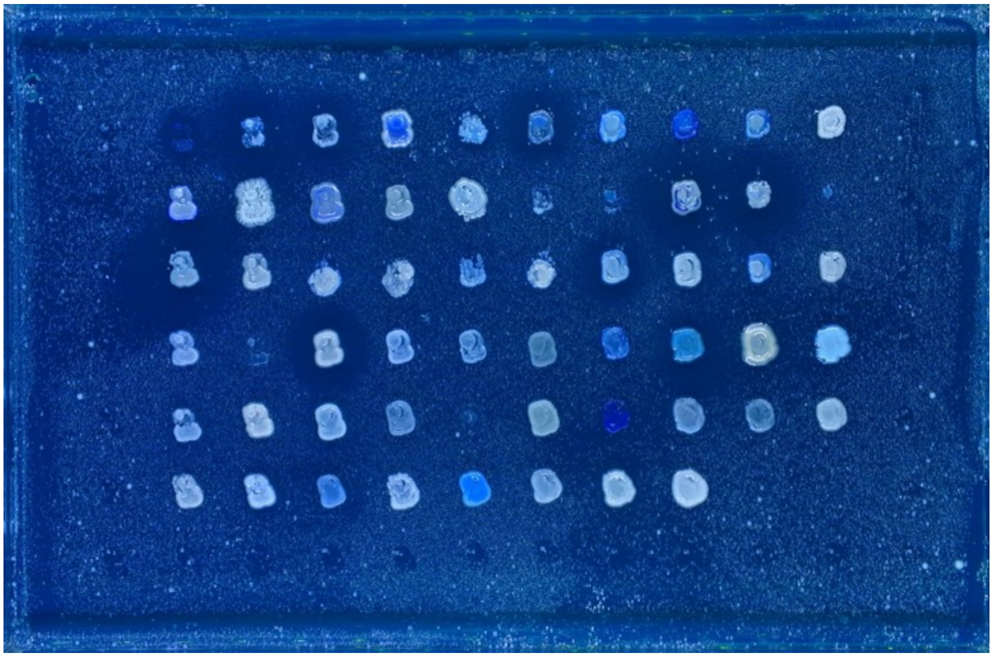

Lawn: CYC 1058

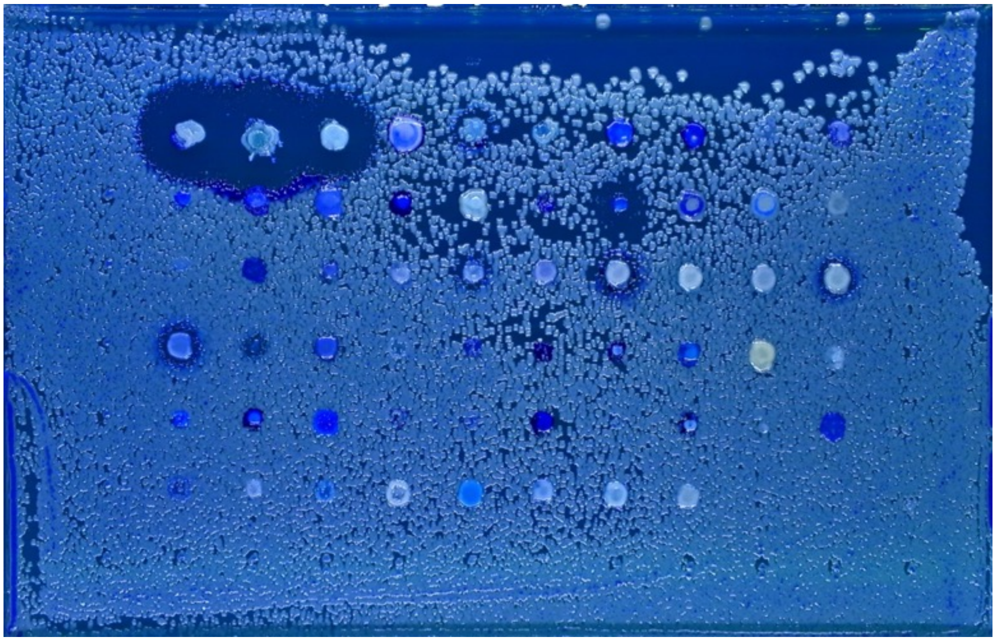

Lawn: CYC 1113

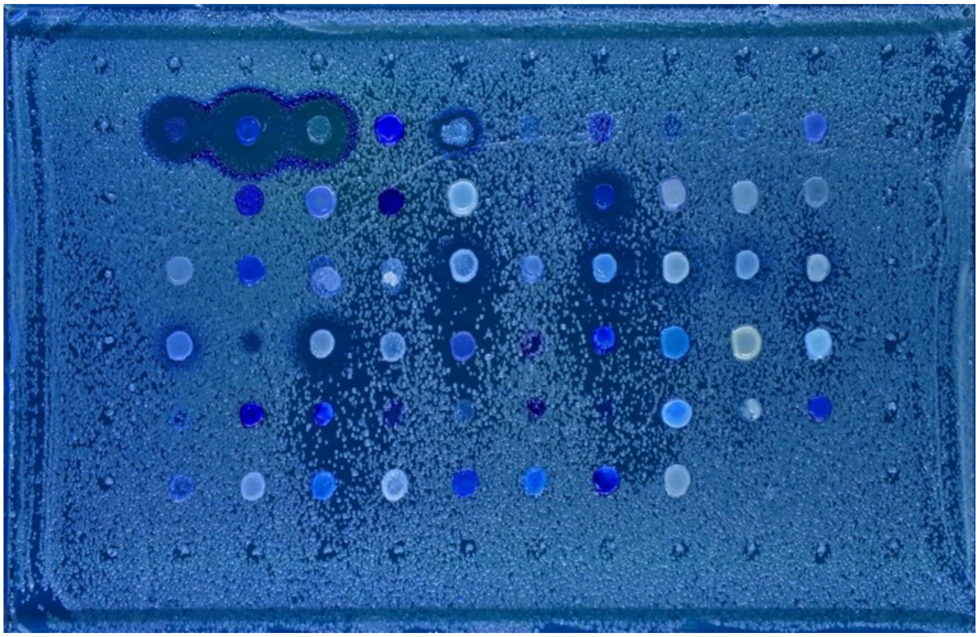

Lawn: CYC 1170

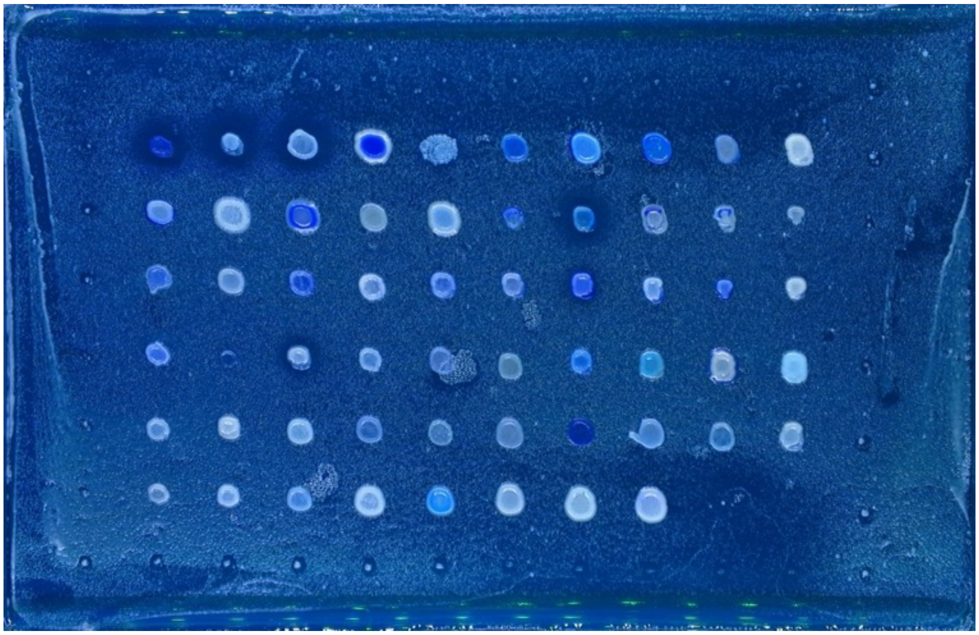

Lawn: CYC 1172

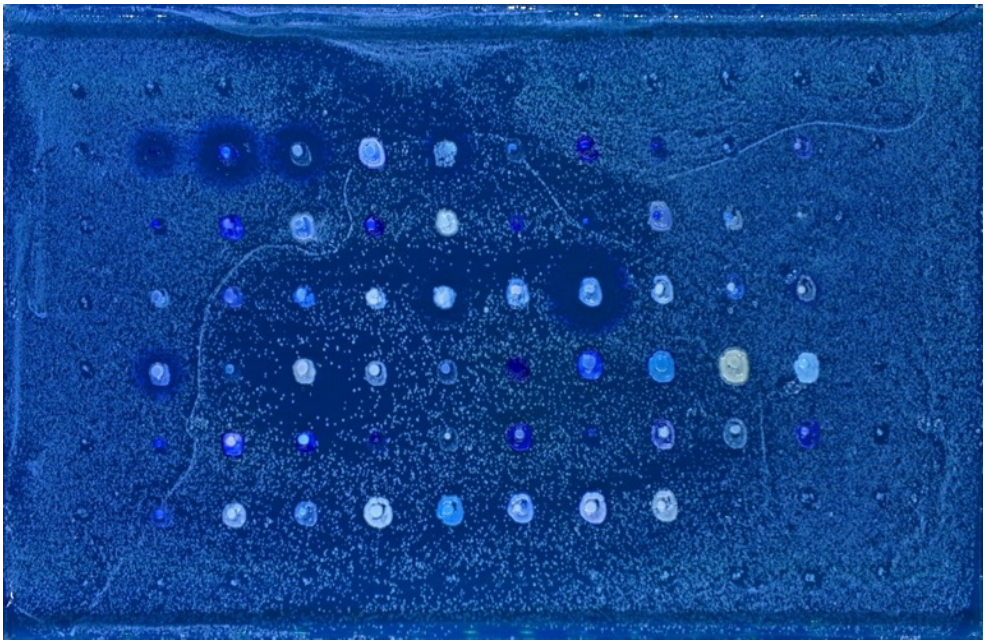

Lawn: Y-1088

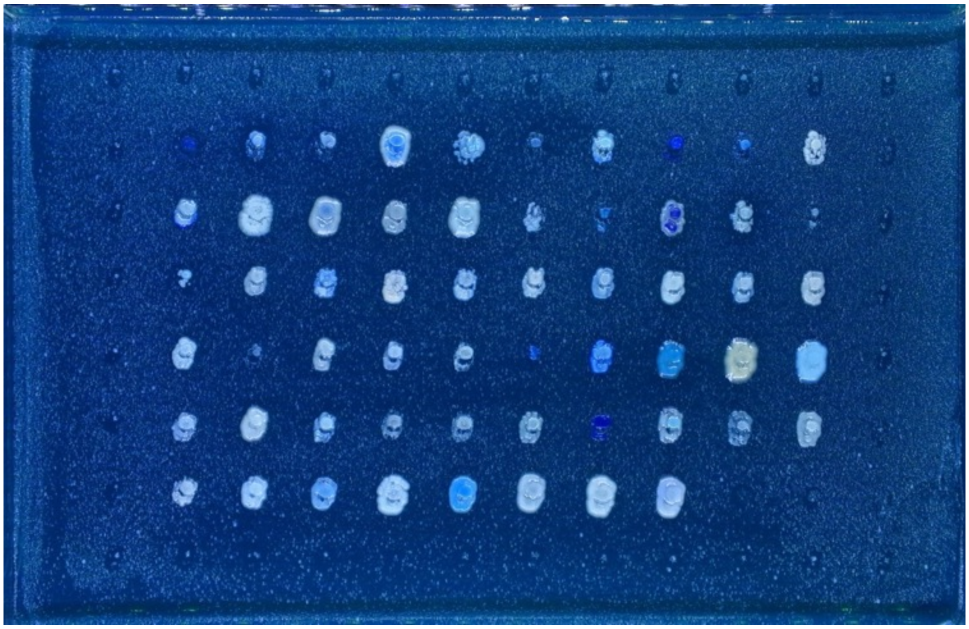

Lawn: Y-27342

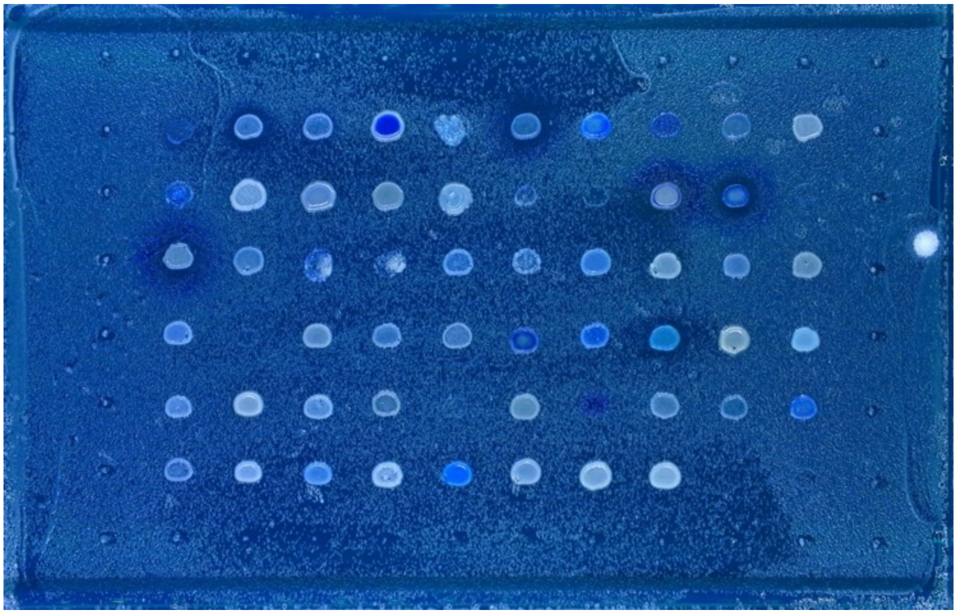

Lawn: Y-2046

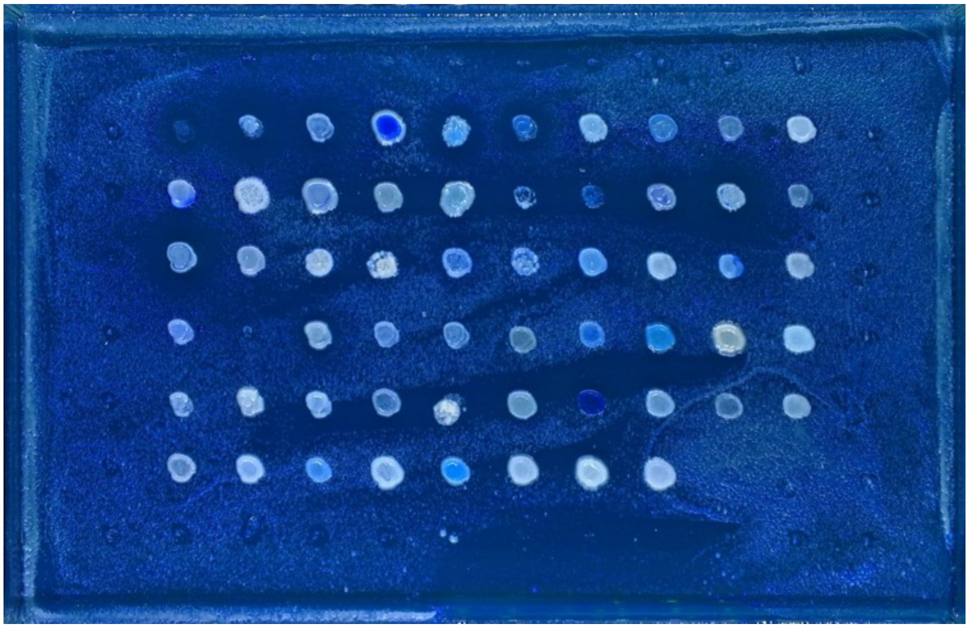

Lawn: Y-1344

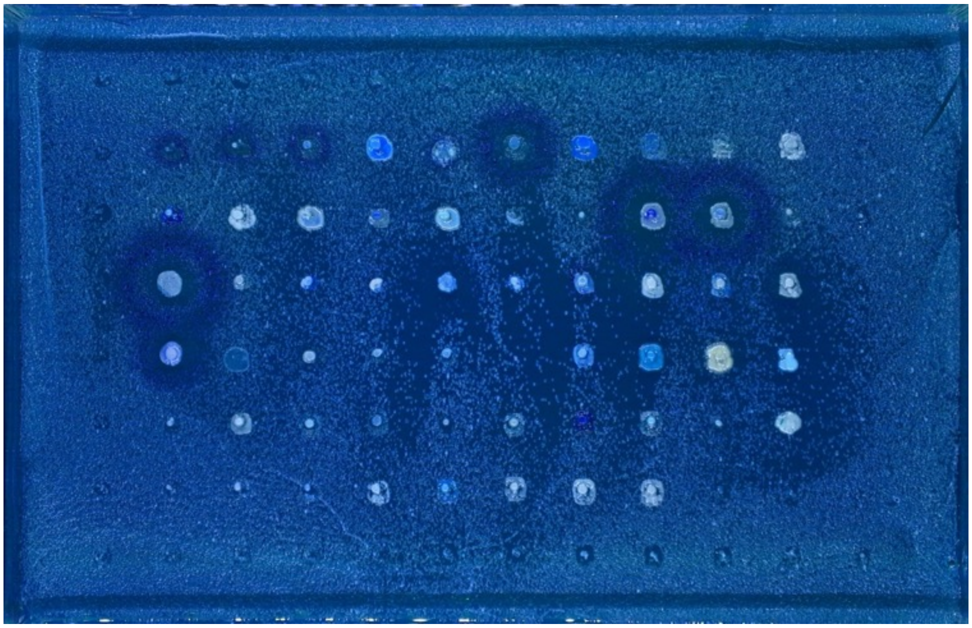

Lawn: NCYC 777

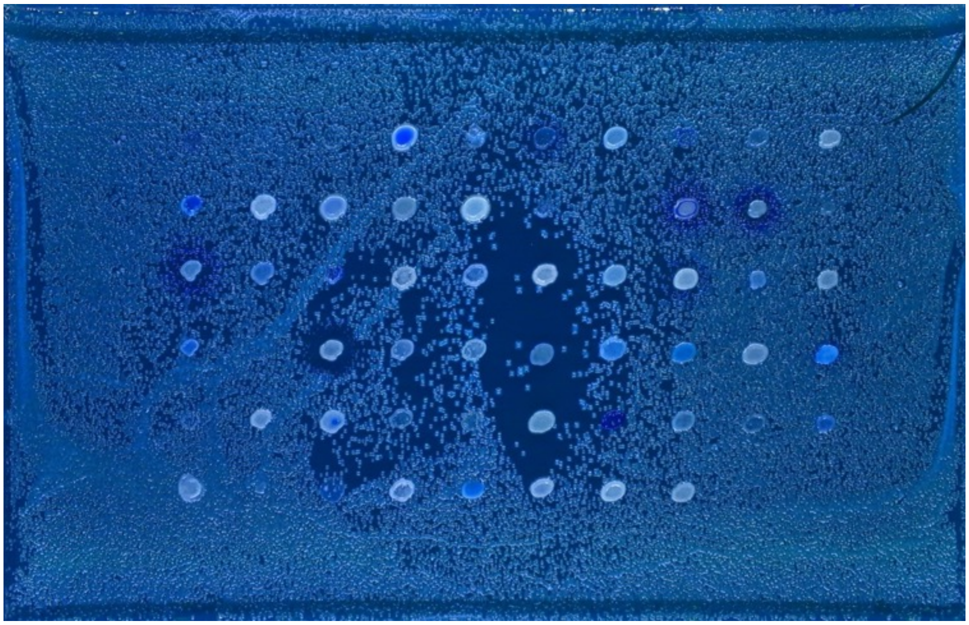

Lawn: NCYC 2898

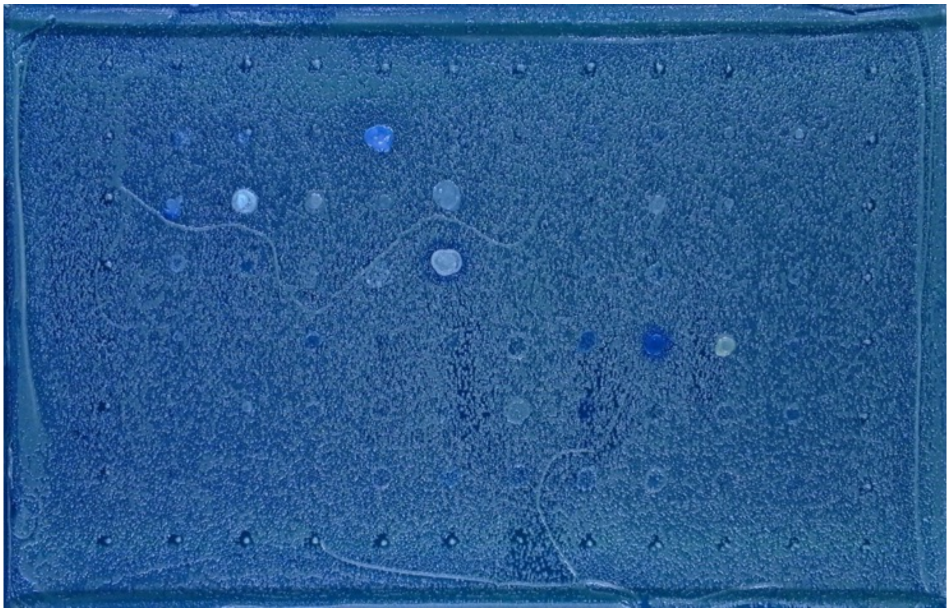

Lawn: 2729

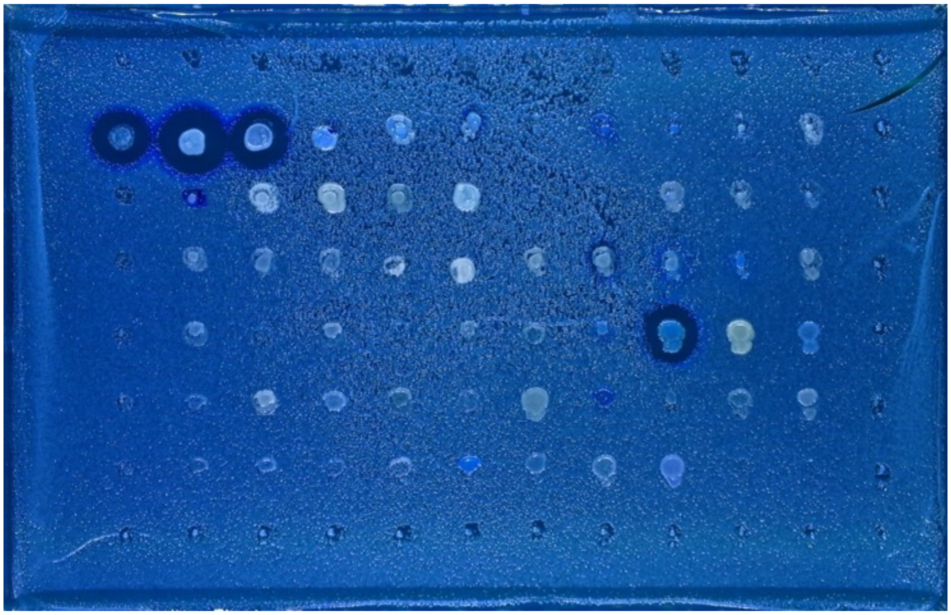

Lawn: FY4

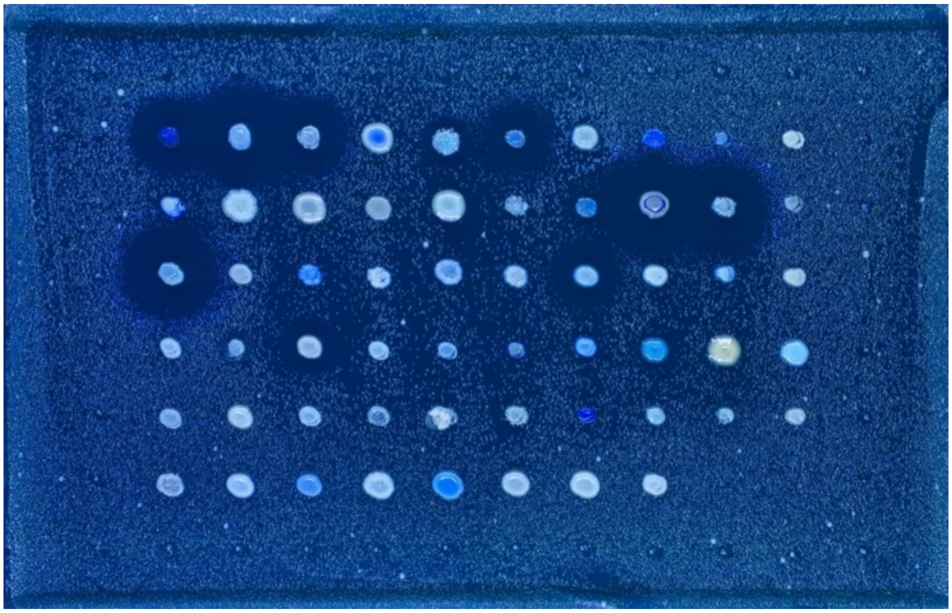

Lawn: K12

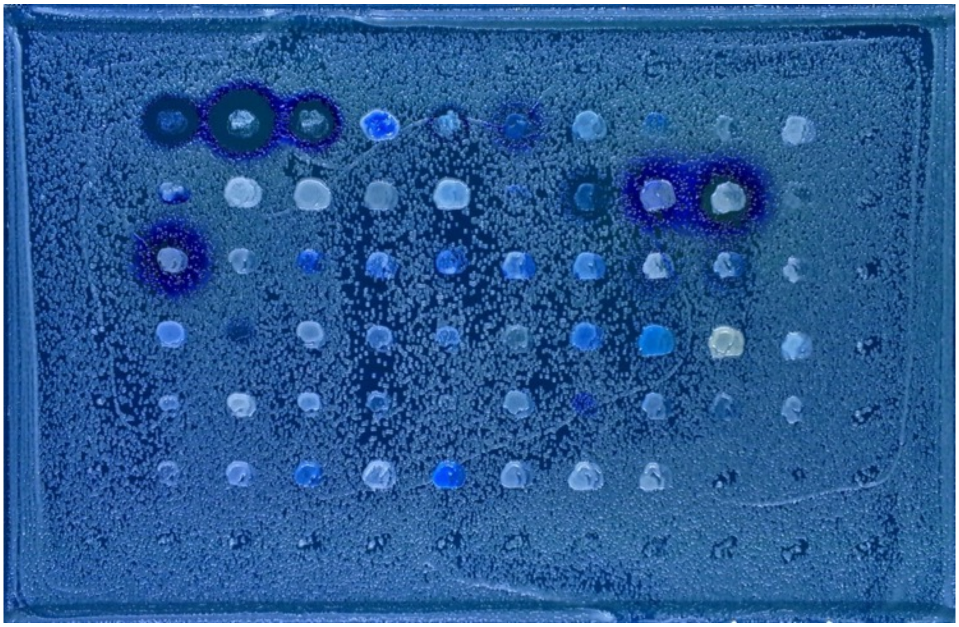

Lawn: BY4741

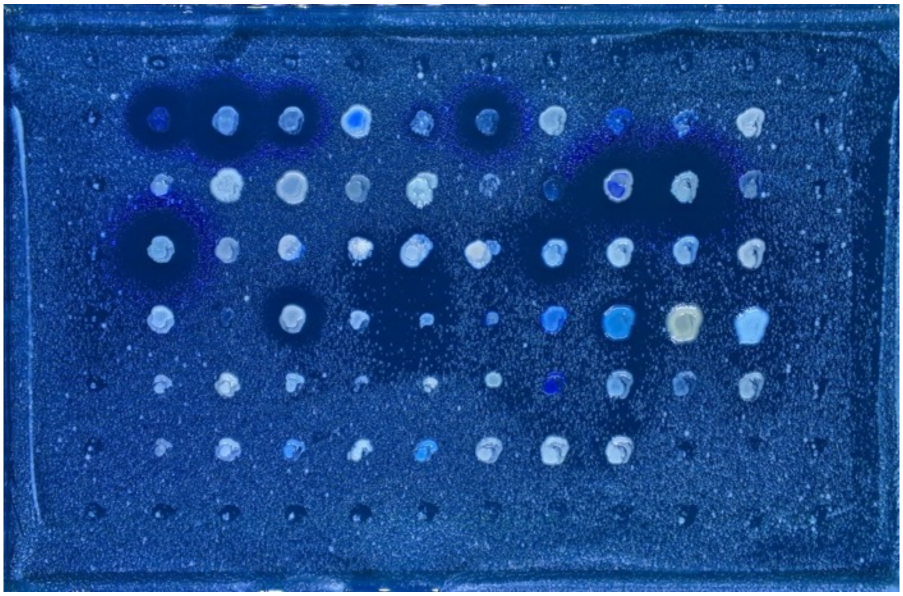

Lawn: DBVPG 6765

Lawn: YB-4237

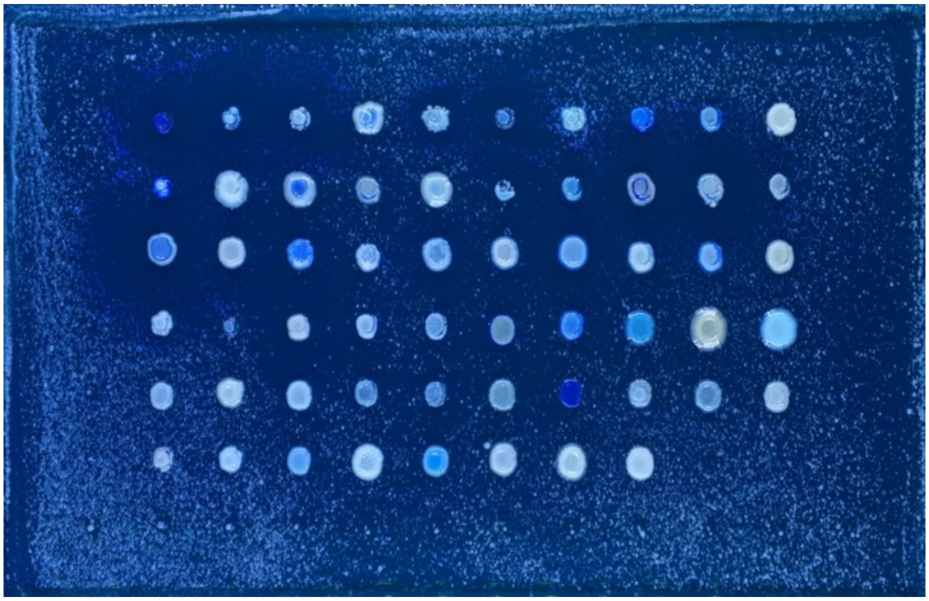

Lawn: CYC 1102

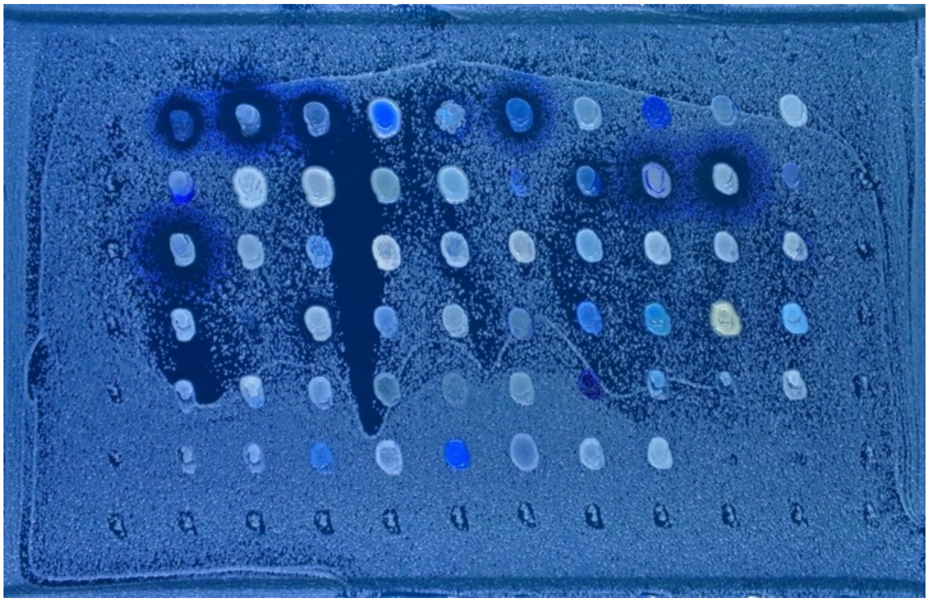

Lawn: YB-4237

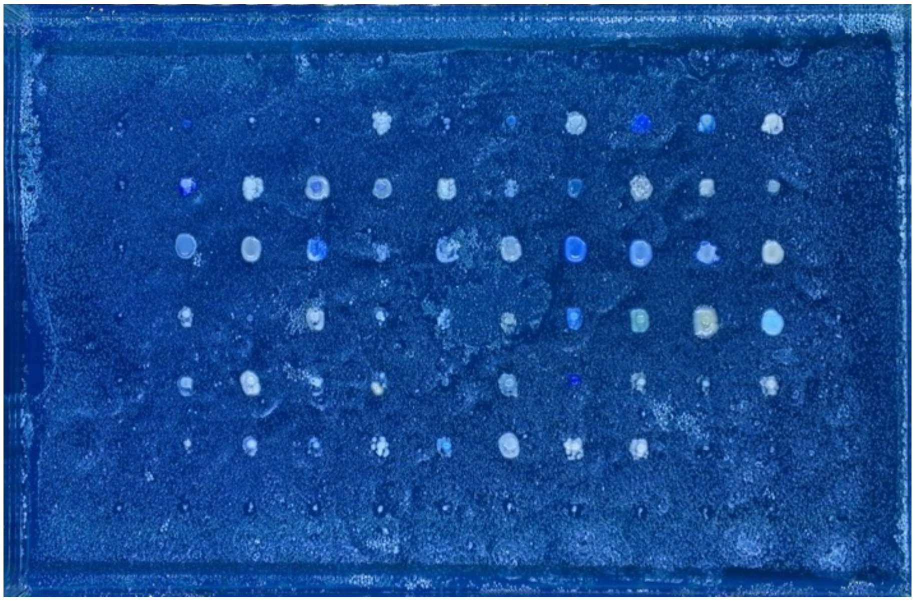

Lawn: CBS 432

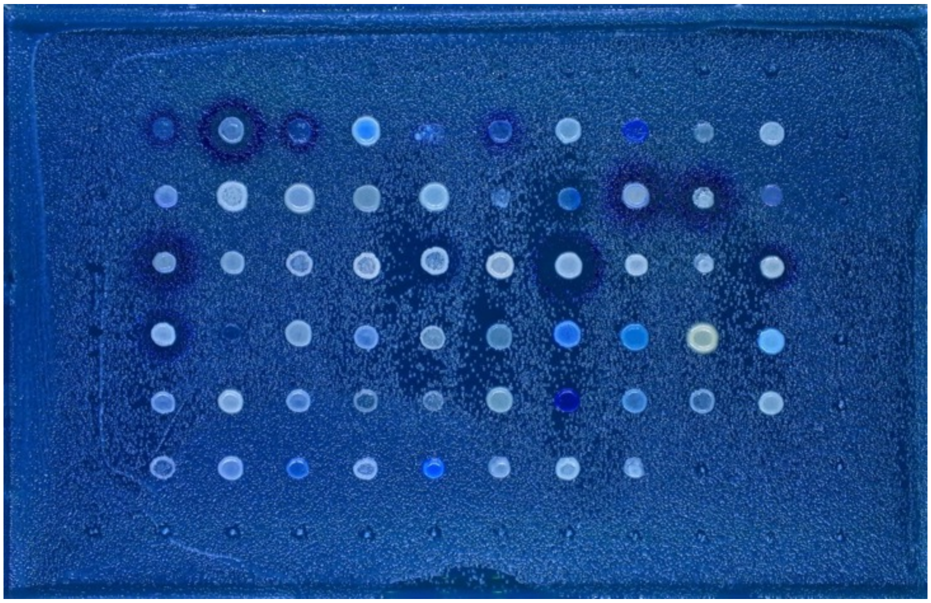

Lawn: A12C

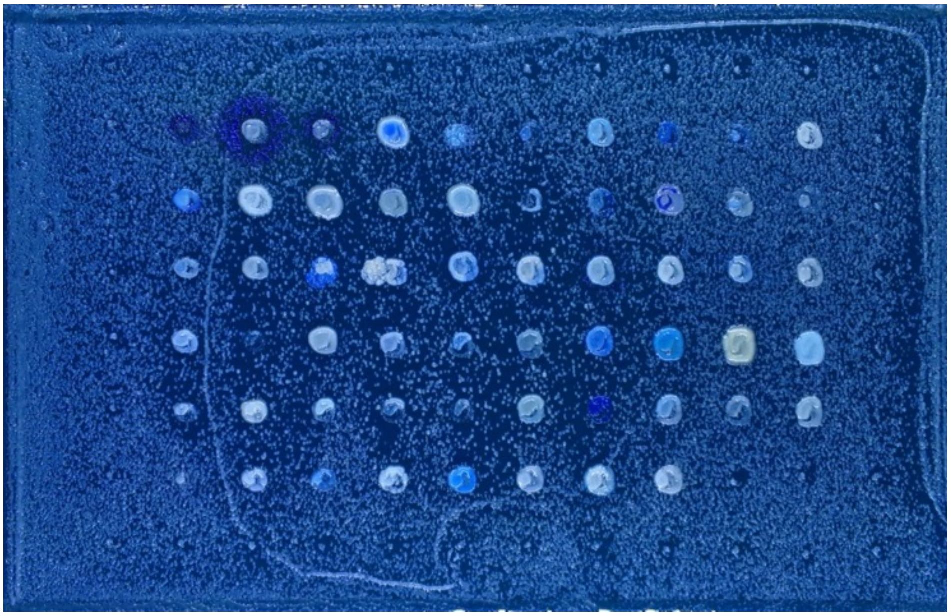

Lawn: NBRC 1815

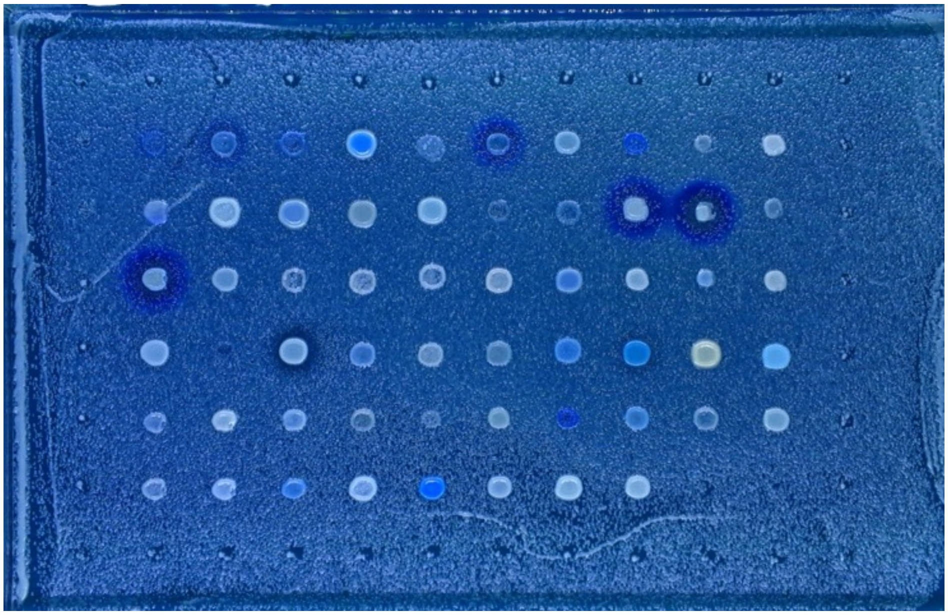

Lawn: SSS 211

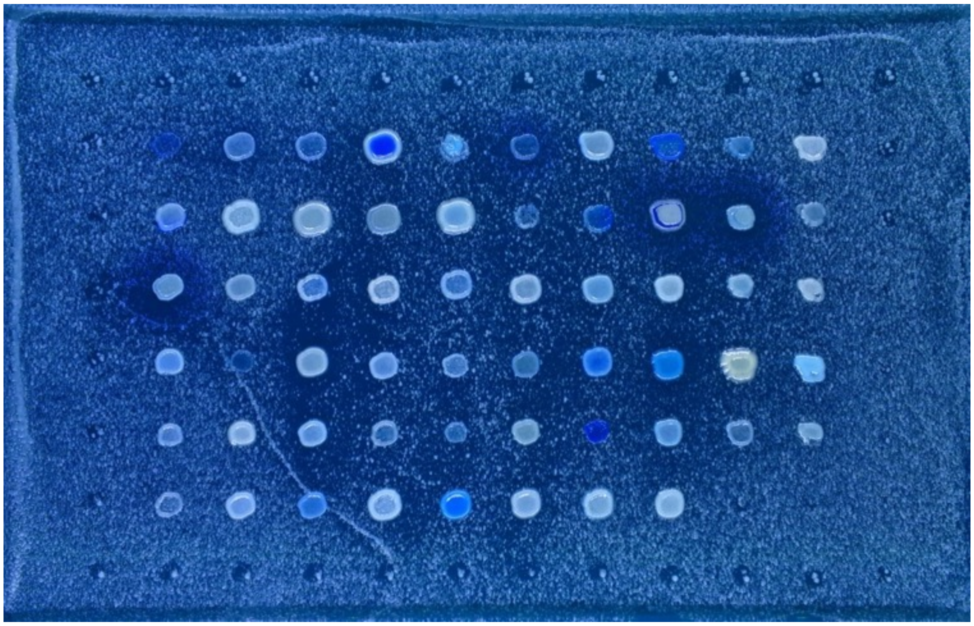

Lawn: NBRC 1802

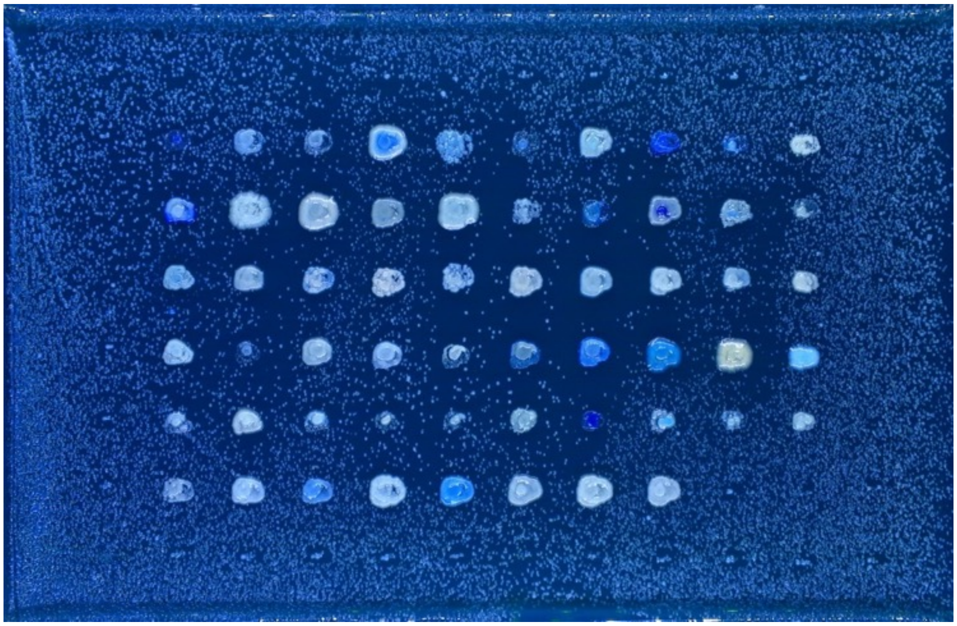

Lawn: CBS 7001

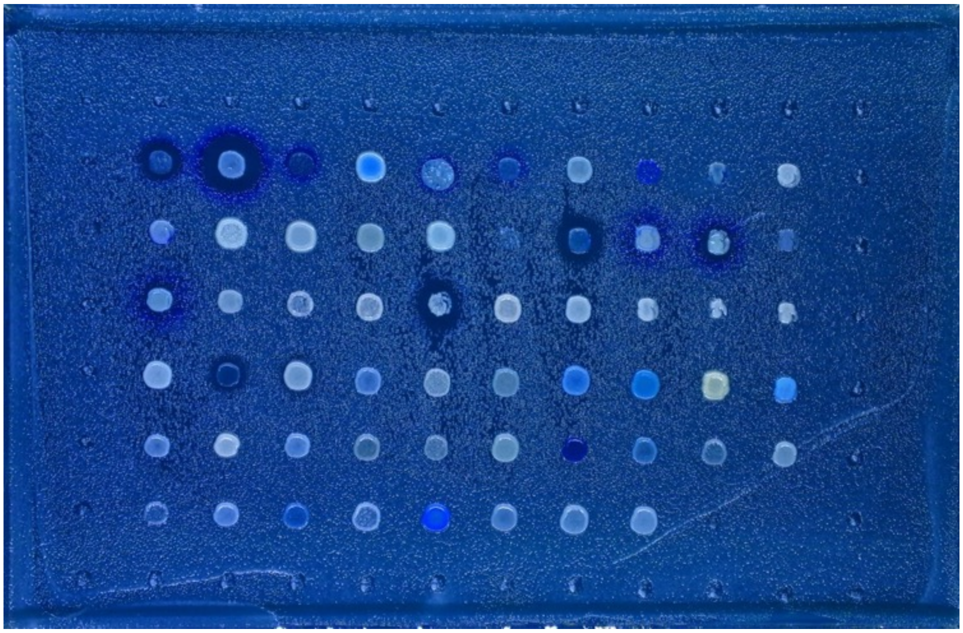

Lawn: SSS 104

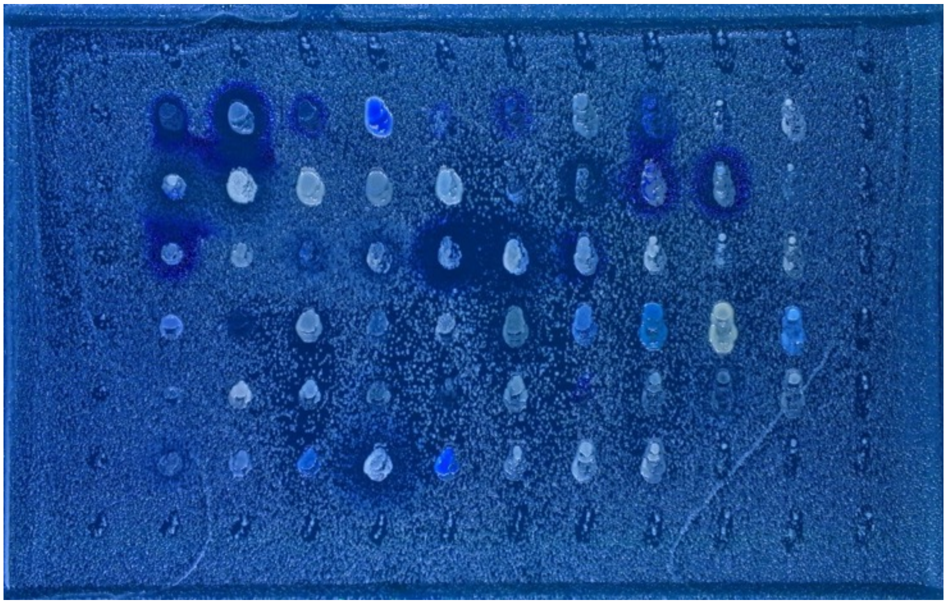

Lawn: Y-63711

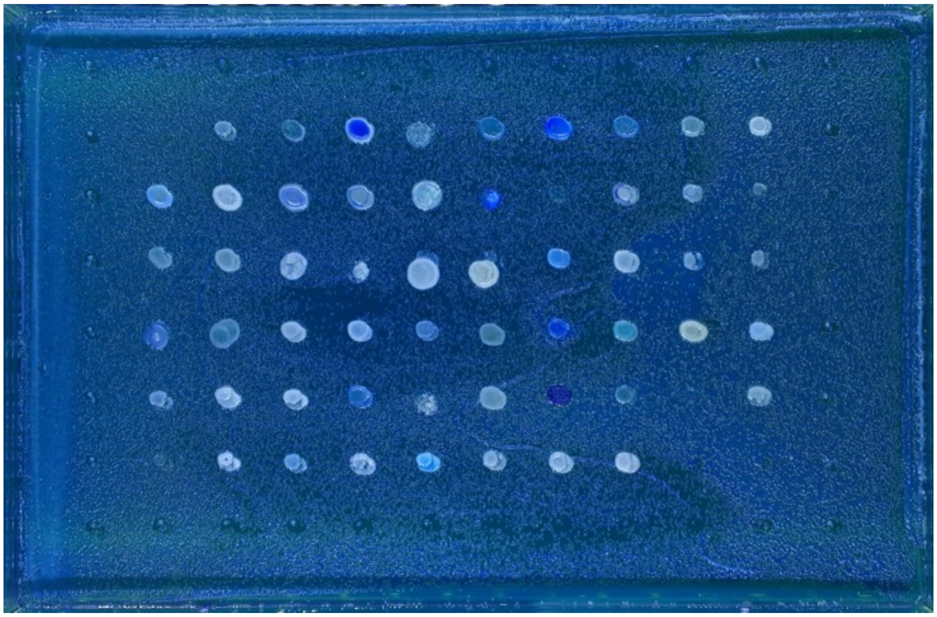

Lawn: Y-63716

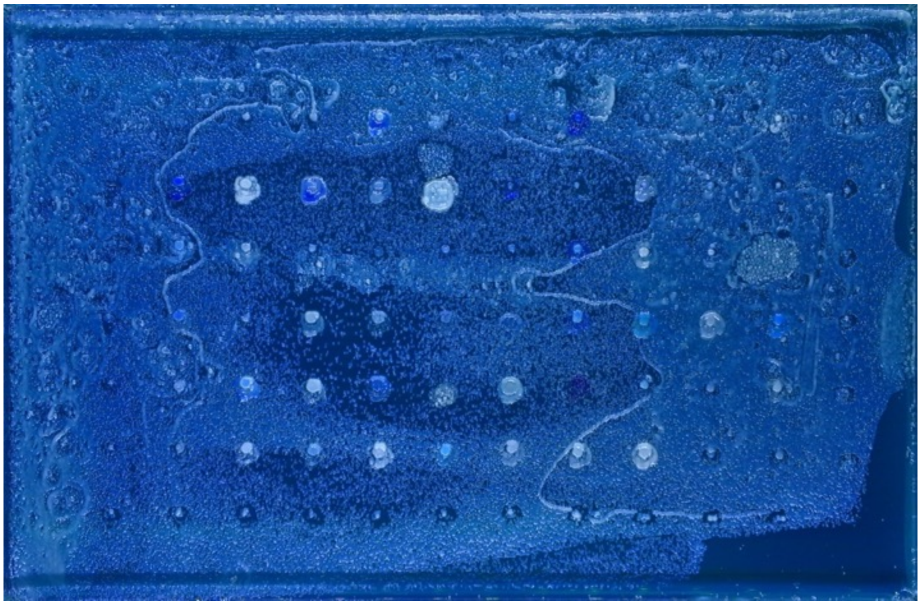

Lawn: YB-4565

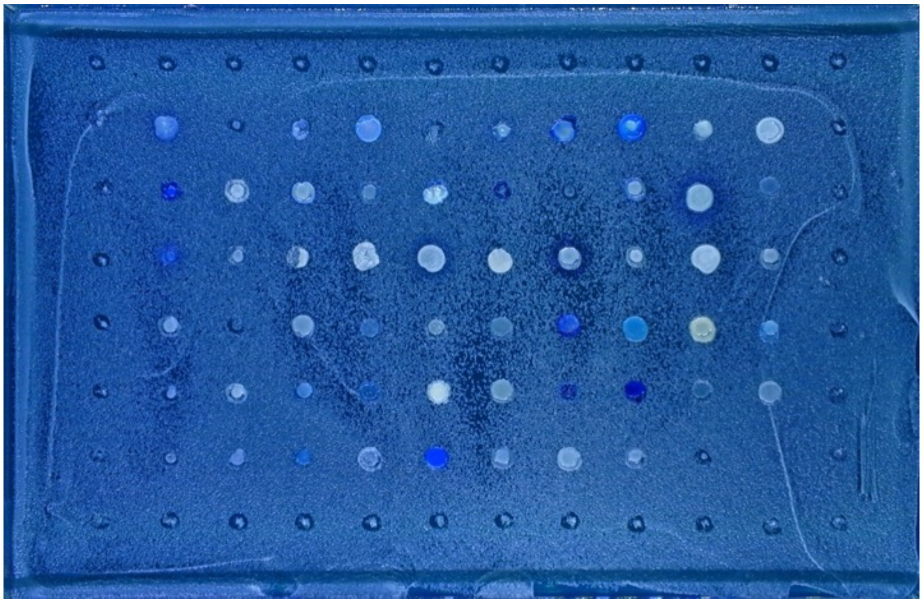

Lawn: 1116

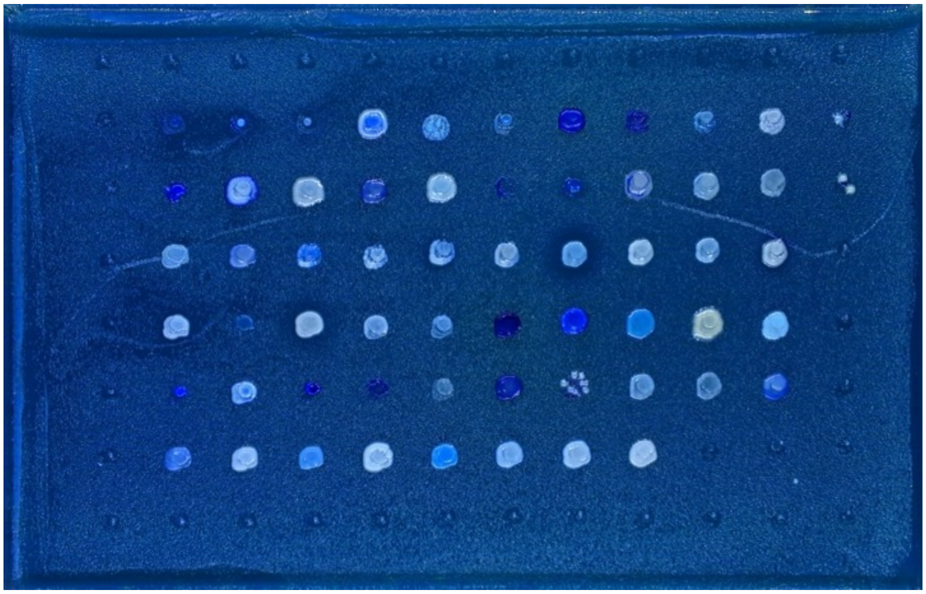

Lawn: Y-2429

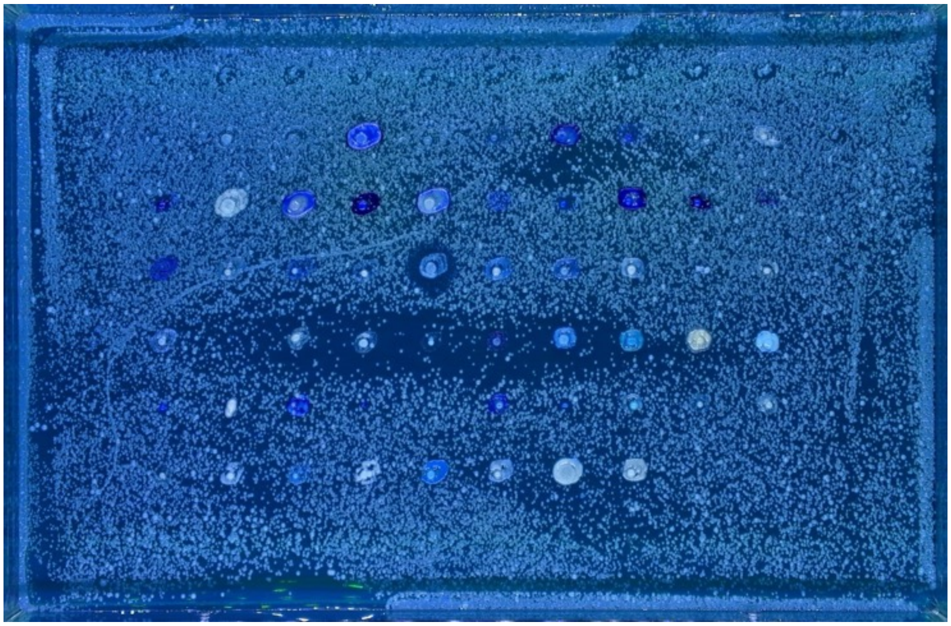

Lawn: DBVPG 6304

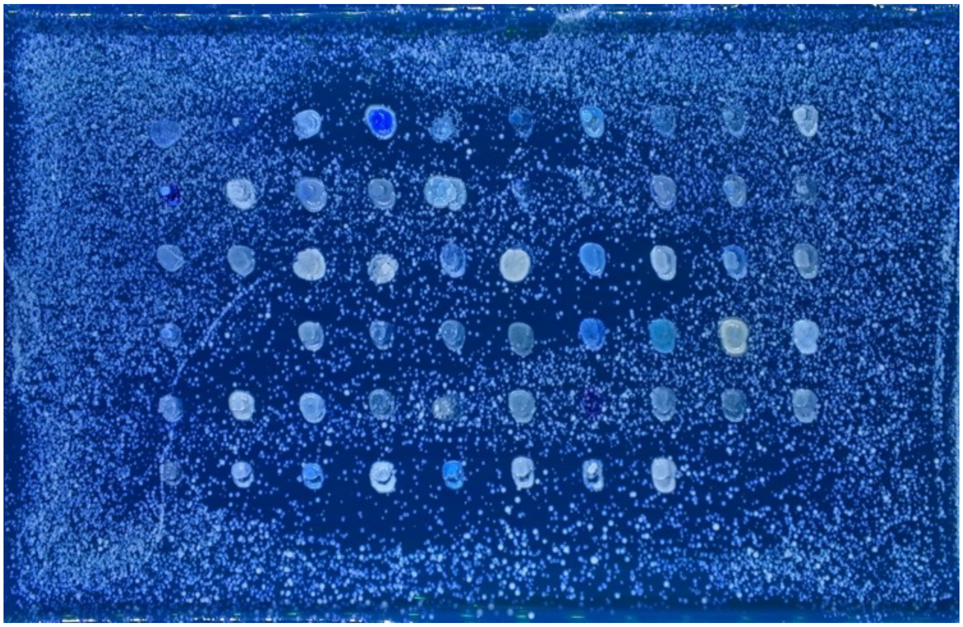

Lawn: Y-63717

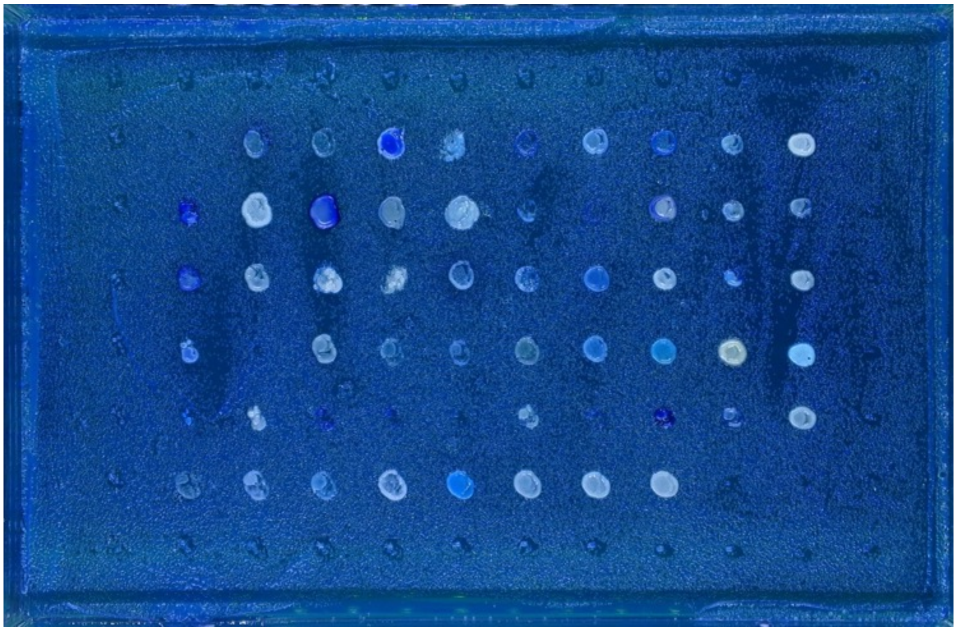

Lawn: Y8.5

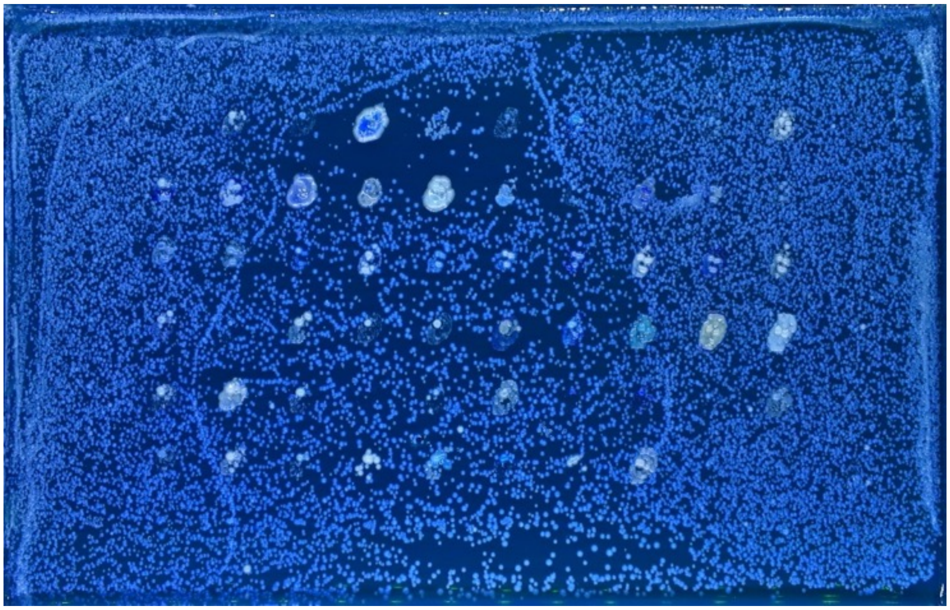

Lawn: Y-12602

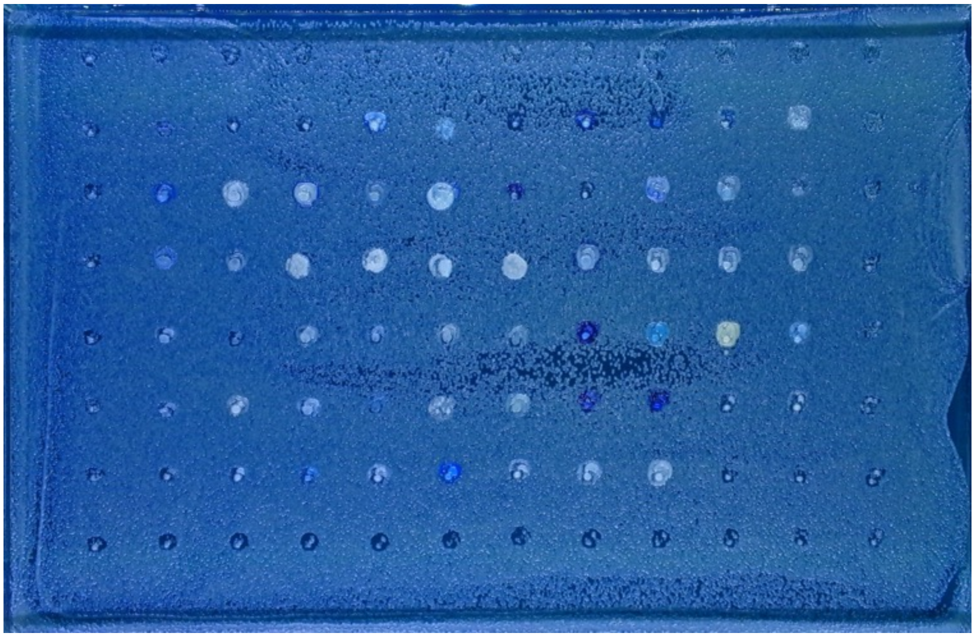
