## Supplementary material for "The Species-specific Acquisition and Diversification of a Novel Family of Killer Toxins in Budding Yeasts of the Saccharomycotina": File S4

Strain list

| Genus | Species | Strain | Source |
|---|---|---|---|
| *Saccharomyces* | *cerevisiae* | BY4741 | Rowley lab |
| *Saccharomyces* | *cerevisiae* | BHJ001 | Rowley lab |
| *Saccharomyces* | *paradoxus* | Y-63717 | NRRL(ARS) |
| *Saccharomyces* | *paradoxus* | A12C | Rowley lab |
| *Saccharomyces* | *cerevisiae* | DSM70459 | DSMZ |
| *Saccharomyces* | *cerevisiae* | NCYC190 | NCYC |
| *Saccharomyces* | *cerevisiae* | CYC1058 | CYC |
| *Saccharomyces* | *cerevisiae* | CYC1113 | CYC |
| *Saccharomyces* | *cerevisiae* | NCYC1001 | NCYC |
| *Saccharomyces* | *paradoxus* | Y8.5 | NCYC |
| *Naumovozyma* | *castelii* | NCYC_2898 | NCYC |
| *Naumovozyma* | *dairenensis* | NCYC_777 | NCYC |
| *Kazachstania* | *africana* | NCYC_2729 | NCYC |
| *Torulaspora* | *delbrueckii* | Y-866 | NRRL(ARS) |
| *Tetrapisispora* | *phaffii* | Y-8282 | NRRL(ARS) |
| *Pichia* | *membranifaciens* | Y-2026 | NRRL(ARS) |
| *Pichia* | *membranifaciens* | Y-6797 | NRRL(ARS) |
| *Pichia* | *membranifaciens* | NCYC_333 | NCYC |
| *Pichia* | *membranifaciens* | NCYC_2788 | NCYC |
| *Pichia* | *membranifaciens* | P43C007 | Okubara lab |
| *Saccharomyces* | *arboricolus* | Y-580 | NRRL(ARS) |
| *Saccharomyces* | *arboricolus* | Y-670 | NRRL(ARS) |
| *Saccharomyces* | *bayanus* | Y-846 | NRRL(ARS) |
| *Saccharomyces* | *bayanus* | Y-969 | NRRL(ARS) |
| *Saccharomyces* | *bayanus* | Y-972 | NRRL(ARS) |
| *Saccharomyces* | *bayanus* | Y-1374 | NRRL(ARS) |
| *Saccharomyces* | *bayanus* | Y-11845 | NRRL(ARS) |
| *Saccharomyces* | *bayanus* | Y-12624 | NRRL(ARS) |
| *Saccharomyces* | *bayanus* | Y-12646 | NRRL(ARS) |
| *Saccharomyces* | *bayanus* | Y-12648 | NRRL(ARS) |
| *Saccharomyces* | *bayanus* | Y-17034 | NRRL(ARS) |
| *Saccharomyces* | *bayanus* | Y-27339 | NRRL(ARS) |
| *Saccharomyces* | *bayanus* | Y-27470 | NRRL(ARS) |
| *Saccharomyces* | *bayanus* | Y-48770 | NRRL(ARS) |
| *Saccharomyces* | *bayanus* | Y-63707 | NRRL(ARS) |
| *Saccharomyces* | *bayanus* | Y-63718 | NRRL(ARS) |
| *Saccharomyces* | *cerevisiae* | YB-254 | NRRL(ARS) |
| *Saccharomyces* | *cerevisiae* | YB-432 | NRRL(ARS) |
| *Saccharomyces* | *cerevisiae* | Y-567 | NRRL(ARS) |
| *Saccharomyces* | *cerevisiae* | Y-851 | NRRL(ARS) |
| *Saccharomyces* | *cerevisiae* | Y-852 | NRRL(ARS) |
| *Saccharomyces* | *cerevisiae* | Y-897 | NRRL(ARS) |
| *Saccharomyces* | *cerevisiae* | Y-898 | NRRL(ARS) |
| *Saccharomyces* | *cerevisiae* | YB-908 | NRRL(ARS) |
| *Saccharomyces* | *cerevisiae* | Y-954 | NRRL(ARS) |

| | | | |
|---|---|---|---|
| *Saccharomyces* | *cerevisiae* | Y-975 | NRRL(ARS) |
| *Saccharomyces* | *cerevisiae* | Y-976 | NRRL(ARS) |
| *Saccharomyces* | *cerevisiae* | Y-977 | NRRL(ARS) |
| *Saccharomyces* | *cerevisiae* | Y-1018 | NRRL(ARS) |
| *Saccharomyces* | *cerevisiae* | Y-1089 | NRRL(ARS) |
| *Saccharomyces* | *cerevisiae* | Y-1285 | NRRL(ARS) |
| *Saccharomyces* | *cerevisiae* | Y-1301 | NRRL(ARS) |
| *Saccharomyces* | *cerevisiae* | Y-1370 | NRRL(ARS) |
| *Saccharomyces* | *cerevisiae* | Y-1428 | NRRL(ARS) |
| *Saccharomyces* | *cerevisiae* | Y-1429 | NRRL(ARS) |
| *Saccharomyces* | *cerevisiae* | Y-1430 | NRRL(ARS) |
| *Saccharomyces* | *cerevisiae* | Y-1436 | NRRL(ARS) |
| *Saccharomyces* | *cerevisiae* | Y-1438 | NRRL(ARS) |
| *Saccharomyces* | *cerevisiae* | Y-1536 | NRRL(ARS) |
| *Saccharomyces* | *cerevisiae* | Y-1540 | NRRL(ARS) |
| *Saccharomyces* | *cerevisiae* | YB-1773 | NRRL(ARS) |
| *Saccharomyces* | *cerevisiae* | Y-1891 | NRRL(ARS) |
| *Saccharomyces* | *cerevisiae* | Y-2044 | NRRL(ARS) |
| *Saccharomyces* | *cerevisiae* | Y-2045 | NRRL(ARS) |
| *Saccharomyces* | *cerevisiae* | Y-2046 | NRRL(ARS) |
| *Saccharomyces* | *cerevisiae* | Y-2204 | NRRL(ARS) |
| *Saccharomyces* | *cerevisiae* | Y-2205 | NRRL(ARS) |
| *Saccharomyces* | *cerevisiae* | Y-2429 | NRRL(ARS) |
| *Saccharomyces* | *cerevisiae* | Y-2430 | NRRL(ARS) |
| *Saccharomyces* | *cerevisiae* | Y-2432 | NRRL(ARS) |
| *Saccharomyces* | *cerevisiae* | Y-2434 | NRRL(ARS) |
| *Saccharomyces* | *cerevisiae* | YB-4237 | NRRL(ARS) |
| *Saccharomyces* | *cerevisiae* | YB-4255 | NRRL(ARS) |
| *Saccharomyces* | *cerevisiae* | YB-4634 | NRRL(ARS) |
| *Saccharomyces* | *cerevisiae* | YB-4635 | NRRL(ARS) |
| *Saccharomyces* | *cerevisiae* | Y-5508 | NRRL(ARS) |
| *Saccharomyces* | *cerevisiae* | Y-5509 | NRRL(ARS) |
| *Saccharomyces* | *cerevisiae* | Y-5510 | NRRL(ARS) |
| *Saccharomyces* | *cerevisiae* | Y-7327 | NRRL(ARS) |
| *Saccharomyces* | *cerevisiae* | Y-7328 | NRRL(ARS) |
| *Saccharomyces* | *cerevisiae* | Y-7567 | NRRL(ARS) |
| *Saccharomyces* | *cerevisiae* | Y-10988 | NRRL(ARS) |
| *Saccharomyces* | *cerevisiae* | Y-11875 | NRRL(ARS) |
| *Saccharomyces* | *cerevisiae* | Y-12842 | NRRL(ARS) |
| *Saccharomyces* | *cerevisiae* | Y-17009 | NRRL(ARS) |
| *Saccharomyces* | *cerevisiae* | Y-17898 | NRRL(ARS) |
| *Saccharomyces* | *cerevisiae* | Y-27105 | NRRL(ARS) |
| *Saccharomyces* | *cerevisiae* | Y-27106 | NRRL(ARS) |
| *Saccharomyces* | *cerevisiae* | Y-27437 | NRRL(ARS) |
| *Saccharomyces* | *cerevisiae* | Y-27788 | NRRL(ARS) |
| *Saccharomyces* | *cerevisiae* | Y-27796 | NRRL(ARS) |
| *Saccharomyces* | *cerevisiae* | y-63703 | NRRL(ARS) |
| *Saccharomyces* | *cerevisiae* | Y-63748 | NRRL(ARS) |

| | | | |
|---|---|---|---|
| *Saccharomyces* | *cerevisiae* | Y-63749 | NRRL(ARS) |
| *Saccharomyces* | *kudriavzevii* | Y-27340 | NRRL(ARS) |
| *Saccharomyces* | *kudriavzevii* | Y-27341 | NRRL(ARS) |
| *Saccharomyces* | *kudriavzevii* | Y-27342 | NRRL(ARS) |
| *Saccharomyces* | *kudriavzevii* | Y-27471 | NRRL(ARS) |
| *Saccharomyces* | *kudriavzevii* | Y-63704 | NRRL(ARS) |
| *Saccharomyces* | *kudriavzevii* | Y-63705 | NRRL(ARS) |
| *Saccharomyces* | *kudriavzevii* | Y-63706 | NRRL(ARS) |
| *Saccharomyces* | *paradoxus* | Y-788 | NRRL(ARS) |
| *Saccharomyces* | *paradoxus* | Y-863 | NRRL(ARS) |
| *Saccharomyces* | *paradoxus* | Y-911 | NRRL(ARS) |
| *Saccharomyces* | *paradoxus* | Y-1088 | NRRL(ARS) |
| *Saccharomyces* | *paradoxus* | Y-1344 | NRRL(ARS) |
| *Saccharomyces* | *paradoxus* | Y-1356 | NRRL(ARS) |
| *Saccharomyces* | *paradoxus* | Y-1548 | NRRL(ARS) |
| *Saccharomyces* | *paradoxus* | Y-1912 | NRRL(ARS) |
| *Saccharomyces* | *paradoxus* | Y-2038 | NRRL(ARS) |
| *Saccharomyces* | *paradoxus* | YB-2047 | NRRL(ARS) |
| *Saccharomyces* | *paradoxus* | YB-4137 | NRRL(ARS) |
| *Saccharomyces* | *paradoxus* | YB-4565 | NRRL(ARS) |
| *Saccharomyces* | *paradoxus* | Y-5688 | NRRL(ARS) |
| *Saccharomyces* | *paradoxus* | Y-6177 | NRRL(ARS) |
| *Saccharomyces* | *paradoxus* | Y-6179 | NRRL(ARS) |
| *Saccharomyces* | *paradoxus* | Y-11842 | NRRL(ARS) |
| *Saccharomyces* | *paradoxus* | Y-12602 | NRRL(ARS) |
| *Saccharomyces* | *paradoxus* | Y-17218 | NRRL(ARS) |
| *Saccharomyces* | *paradoxus* | Y-17353 | NRRL(ARS) |
| *Saccharomyces* | *paradoxus* | Y-63708 | NRRL(ARS) |
| *Saccharomyces* | *paradoxus* | Y-63709 | NRRL(ARS) |
| *Saccharomyces* | *paradoxus* | Y-63710 | NRRL(ARS) |
| *Saccharomyces* | *paradoxus* | Y-63711 | NRRL(ARS) |
| *Saccharomyces* | *paradoxus* | Y-63712 | NRRL(ARS) |
| *Saccharomyces* | *paradoxus* | Y-63713 | NRRL(ARS) |
| *Saccharomyces* | *paradoxus* | Y-63714 | NRRL(ARS) |
| *Saccharomyces* | *paradoxus* | Y-63715 | NRRL(ARS) |
| *Saccharomyces* | *paradoxus* | Y-63716 | NRRL(ARS) |

Primers

| Name | Sequence (5’-3’) | Notes |
|---|---|---|
| PRUI185 | AACATTTTCGGTTTGTATTACTTCTTATTCTCTAAAAATGAGAAATAGTACC | K1L amplification for cloning |
| PRUI186 | GTATCGTGATGACAGAGGCAGGGAGTGGGATCAAGCGCCAGTATCGCATTGGCTCC | |
| PRUI054 | caactgaaaacactccatctgtttcttacc | K1L amplification from P. membranifaciens NCYC333 |
| PRUI055 | ttaactcccagtatcacattcagtttcgtatgg | |
| PRUI052 | tgcaaggcttgaaaaatgaagttagc | K1L amplification from Naumovozyma dairenensis CBS 421 chr 7 |
| PRUI053 | catttgaaggcctatagggaagagg | |
| PRUI056 | tgactagaaactcatgtcgccaacc | K1L amplification from Kazachstania africana CBS2517 chr 1 |
| PRUI057 | aaagtgaaatttgcaagatcattagtaccc | |
| PRUI060 | tgttgaagctaaacagttttaacagagtgg | K1L amplification from Kazachstania africana CBS2517 chr 12 |
| PRUI061 | gtcgaacaagaagggaaaattacgg | |
| PRUI233 | GACCAGCGGATAAACAGTATGTGC | K1L amplification from Tetrapisispora phaffii CBS 4417 chr 2 |
| PRUI234 | TTTTACAAAAGAAAATGCAAGCAAGC | |
| PRUI235 | AACAACCGATTCATTAGCAATAGGC | K1L amplification from Tetrapisispora phaffii CBS 4417 chr 11 |
| PRUI236 | AACGCCTTCTTATTTAAGCGTCTCG | |
| PRUI237 | AACGCCTTCTTATTTAAGCGTCTCG | K1L amplification from Naumovozyma castellii CBS 4309 chr 6 |
| PRUI238 | TGTGTGAACCATTCTTCAACGTACC | |
| PRUI239 | ACCATATTGGGGTTATTTTCGTTCG | K1L amplification from Naumovozyma dairenensis CBS 421 chr 8 |
| PRUI240 | TTGTTGAAGACATAAAAACGCATCG | |
| PRUI241 | AAAAACGCATATTGAAGTTGTTCTCG | K1L amplification from Naumovozyma dairenensis CBS 421 chr 3 |
| PRUI242 | GTGCCTTAATGAAACTTAGTGGTTGG | |
| PRUI243 | atgagagttgttggaatttttactgttttgg | K1L amplification from Tetrapisispora phaffii CBS 4417 chr 1 |
| PRUI244 | ctaacttcctgtatcacatgaatccc | |
| PRUI245 | TGAGAAATAGTACCTTCACCCTTAATTTGA | Amplify 5' region of M1L from Y-63717 (Figure S3 RXN1) |
| PRUI246 | TTTGGCTCTGCACTTTACTCTATACTACG | |
| PRUI247 | AACTGGTAGTCCACGTCACTTACGG | Amplify 3' region of M1L from Y-63717 (Figure S3 RXN2) |
| PRUI248 | TTGTTCAGTATTCAGGACCTATTGTCG | |
| PRUI249 | ACAGGTTGTGGAACAGTATTTGTGG | Amplify central poly(A) region of M1L from Y-63717 (Figure S3 RXN3) |
| PRUI250 | GATGTCAGATATGCCAGCTTTTCC | |
| AMC001 | CCTATAGATATCTTAACAGC | Use for 5' RACE kit as GSP1 for top strand targeting |
| AMC002 | TAACAGAATTAACTACTCAG | Use for 5' RACE kit as GSP1 for bottom strand targeting |

Plasmid list

| Name | Description |
|---|---|
| pUI067 | pDONR221 w/ K1L Y-63717 |
| pUI109 | pAG426-Gal w/ T. phaffi chromosome 2 |
| pUI110 | pAG426-Gal w/ T. phaffi chromosome 11 |
| pUI111 | pAG426-Gal w/ T. phaffi chromosome 1 |
| pUI112 | pAG426-Gal w/ N. dair chromosome 8 |
| pUI113 | pAG426-Gal w/ N. dair chromosome 3 |
| pUI114 | pAG426-Gal w/ N. castelli chromosome 6 |
| pML115 | pAG426-Gal w/ N. dair chromosome 7 |
| pML116 | pAG426-Gal w/ P. memb |
| pML117 | pAG426-Gal w/ K. africana chromosome 1 |
| pML118 | pAG426-Gal w/ K. africana chromosome 12 |
| pUI119 | pAG426-Gal w/ S. paradoxus K1L |
